# GeroEngine: Generative single-cell aging trajectories reveal a bidirectionally traversable identity core and direction-specific inflammatory remodeling

**DOI:** 10.64898/2026.06.08.731002

**Authors:** Youngmin Bhak, Sungwon Jeon, Jong Bhak

## Abstract

Single-cell RNA sequencing (scRNA-seq) maps aging tissues at high resolution but is destructive, preventing longitudinal tracking; dropout and zero-inflation artifacts, amplified by shift-invariant linear simulations, confound age-associated variability. We developed GeroEngine, a technical-artifact-aware framework combining VAE-based trajectory simulation, LOPO cross-validation, linear baselines, reverse traversal, and reverse-directed network inference. In microglia and HSCs, the VAE reduced technical-artifact carryover while preserving trajectory heterogeneity and improving alignment to artifact-reduced reference manifolds. Consensus GeroTargets and GeroRegulators defined tissue-specific GeroNetworks organized into three pillars: lineage/replication identity collapse, a sex-dimorphic endocrine/stress core, and inflammatory remodeling. Forward and reverse simulations aligned to the common young*→*old aging axis revealed a sign-coherent, direction-specific program: identity/replication targets were bidirectionally recovered, whereas MHC/NF-*κ*B inflammatory programs were preferentially forward-recovered. These results support identity collapse as a deep traversable core of aging and nominate upstream homeostatic restoration over downstream inflammatory suppression.

## 1. Introduction

Biological aging is not a uniform decline, but a highly heterogeneous, non-linear process driven by stochastic transcriptomic shifts at the single-cell level.[1–4] Single-cell RNA sequencing (scRNA-seq) has revolutionized our ability to map tissue-specific aging trajectories, revealing cellular senescence patterns across critical systems such as the brain and bone marrow.[5] However, this methodology presents a fundamental limitation: scRNA-seq is inherently destructive. While we can capture high-resolution cross-sectional snapshots of diverse cell populations at distinct chronological ages, we cannot longitudinally track the exact same cell across its lifespan.[6] To overcome this physical limitation, researchers must develop *in silico* models that simulate longitudinal aging, taking a real young cell and predicting its probable future transcriptomic state. Yet forecasting these states is severely hindered by the inherent noise of scRNA-seq data, including zero-inflation and stochastic dropout,[7, 8] as well as the natural accumulation of transcriptional noise as cells age.[2, 9] Any viable simulation model must therefore capture the complex biology of aging while distinguishing putative regulatory shifts from technical artifacts, without discarding biologically meaningful stochastic variability that accumulates with age.

Deep generative models, particularly Variational Autoencoders (VAEs),[10] have emerged as a foundational standard for this challenge. By passing sparse count matrices through a non-linear latent bottleneck and explicitly modeling the underlying statistical distribution of the data, VAEs have been widely used to reduce dropout-associated technical artifacts and recover smoother representations of single-cell expression, although such reconstructions should not be interpreted as removing biological aging heterogeneity.[11, 12] A common baseline for modeling single-cell state shifts instead uses Principal Component Analysis (PCA)[13, 14] or a direct algebraic transition. However, because linear models are strictly shift-invariant, they are vulnerable to carrying source-cell technical artifacts and idiosyncratic measurement noise into the simulated aged state, rather than learning a context-dependent biological trajectory. When applied to simulate longitudinal aging, a linear vector acts as a rigid translation that blindly propagates the technical artifact and sparsity structure of the young cell into the aged state. As our benchmarking demonstrates, this inability to reduce technical artifact carryover produces simulated profiles that can be topologically distorted and less able to recapitulate the non-linear structure of the aging manifold.

While standard trajectory inference and optimal transport algorithms (e.g., scVelo, CellRank, Moscot) have revolutionized our ability to model continuous cellular differentiation,[15–17] they are not naturally suited to chronological aging across long, sparsely sampled age intervals.[16, 18] These models rely on continuous transitional states or RNA velocity gradients, which are often absent across long chronological gaps (e.g., 3 to 24 months); moreover, differentiation is a highly programmed funnel, whereas aging is driven by stochastic, system-wide functional decay. To reconstruct directional aging dynamics from static snapshots, we developed the GeroEngine, a unified, hypothesis-generating framework that integrates non-linear generative trajectory simulation (built upon the scGen architecture[19]) with strict cross-validation and reverse-directed network inference. We provide evidence that this framework reduces scRNA-seq technical artifacts while estimating a reproducible, population-level aging vector; cumulative exposome-driven variability is treated as part of the aging phenotype rather than as noise to remove. By doing so, it moves beyond predicting a future transcriptomic state to computationally approximating the directional aging trajectory (GeroVector), enabling systematic generation of hypotheses regarding the upstream regulatory drivers (GeroRegulators) that orchestrate the collapse of cellular identity.

In this study, we systematically benchmark generative versus linear simulation models across key aging tissues (Microglia and Hematopoietic Stem Cells [HSCs]). We demonstrate that VAEs preserve cellular heterogeneity and achieve superior global manifold fidelity against technical-artifact-reduced reference representations. Within a strict Leave-One-Pair-Out (LOPO) cross-validation framework, we provide evidence that generative inference can estimate a reproducible aging-associated trajectory while reducing dropout-associated artifacts and preserving age-associated cell-to-cell variability. We do not treat the elevated variance of true aged populations as removable noise; instead, the VAE reduces technical artifacts and is assessed for whether the learned aging trajectory is consistent with transcriptomic contraction and loss of cellular identity. Finally, we show that this technical-artifact-aware framework improves directional aging inference under cross-sex transfer, while revealing that accurate topological reconstruction in strongly sex-dimorphic tissues may require sex-specific decoders.

## 2. Results

### 2.1. VAE Preserves the Distributional Shape of Aging While Suppressing Technical Noise

At the pseudo-bulk population level, both the non-linear VAE and the linear models (PCA and Algebraic Linear Shift [ALS]) successfully captured the global mean expression state (Figure 1A) and the broad directionality of the aging vector, measured by the Pearson correlation of transcriptomic shifts (*R*_*shift*_; Figure 1B). However, macroscopic averages obscure single-cell topology: two-dimensional (2D) PCA visualizations (Figure 1H) revealed that linear models produced artificially uniform trajectories, forcing cells along rigid parallel tracks that failed to capture the distributional shape of the aged manifold.

**Figure 1.**
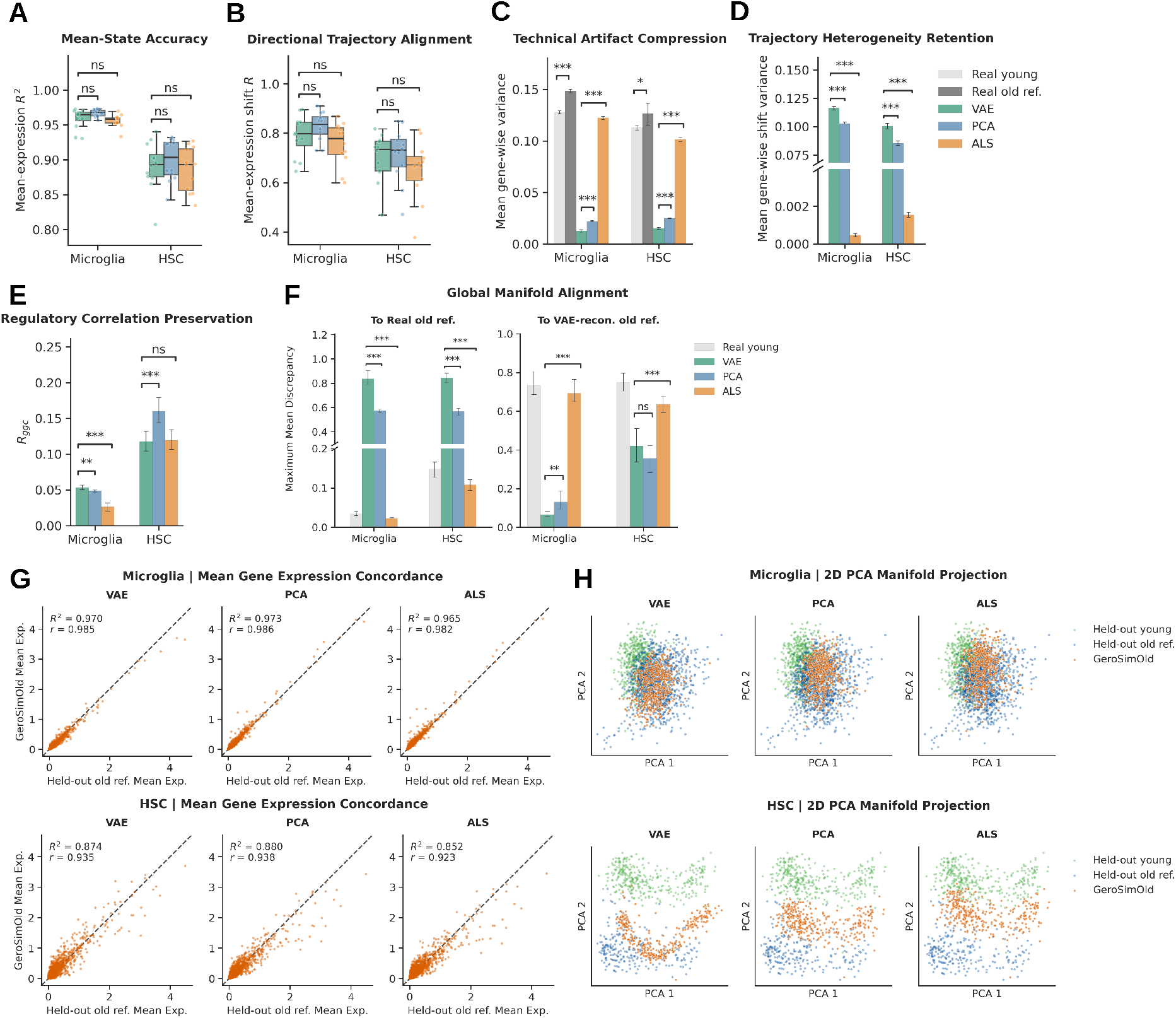
Benchmarking of longitudinal trajectory simulation fidelity. Metrics comparing the non-linear VAE against linear baselines (Principal Component Analysis [PCA] and Algebraic Linear Shift [ALS]) across 12 validation folds. (A) *R*^2^ of mean expression pseudo-bulks. (B) Pearson correlation of transcriptomic shifts *R*_*shift*_. (C) Mean of gene variances 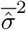, indicating reduced technical artifact carryover, interpreted together with preserved shift variance. (D) Mean of gene shift variances 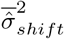 highlighting trajectory-heterogeneity retention. (E) Gene-gene correlation metrics evaluating network preservation. (F) Maximum Mean Discrepancy (MMD) scoring global manifold alignment. (G) 2D scatter of gene average correlation between target and predicted mean expressions. (H) 2D PCA embeddings visualizing simulated trajectories. Panels (G,H) display representative simulations using the 3 8 M and 24 58 M hold-out pair (LOPO fold 7 microglia; fold 5 HSCs). Significance: paired Wilcoxon signed-rank test with Benjamini-Hochberg false-discovery-rate (BH-FDR) correction (* *p <* 0.05, ** *p <* 0.01, *** *p <* 0.001, ns: non-significant). ref, reference; recon, reconstructed; pred, prediction; Exp, Expression.

To explain this discrepancy, we interrogated two aspects of simulation variance. Empirical baselines confirmed that biological aging naturally increases absolute transcriptomic variance *in vivo* (Microglia Real Young = 0.128 *±* 0.003 vs. Old = 0.148 *±* 0.003, *p <* 0.05; HSC Young = 0.113 *±* 0.004 vs. Old = 0.127 *±* 0.019, *p <* 0.05; Figure 1C). When simulating this process, the strictly shift-invariant ALS applied an identical scalar shift to every cell, practically eliminating natural stochasticity (Microglia 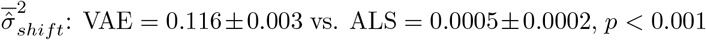), whereas the VAE preserved trajectory heterogeneity (Figure 1D). For the absolute variance of the final predicted state, PCA’s linear projection struggled to compress zero-inflated structural artifacts, retaining higher variance, while the non-linear VAE achieved significantly stronger artifact compression (Microglia 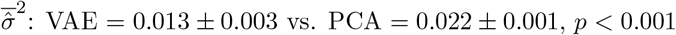; Figure 1C). Because 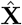 represents decoded continuous reconstructions rather than sparse counts, low 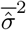 was interpreted only together with preserved shift variance, manifold alignment, and pathway-level concordance.

The VAE also preserved coordinated regulatory structure: by evaluating gene-gene co-expression against a technical-artifact-reduced reference rather than raw dropout-affected data, the VAE produced significantly higher global gene-gene correlation scores (*R*_*ggc*_; Microglia: VAE = 0.053 *±* 0.006 vs. PCA = 0.049 *±* 0.003, *p <* 0.01; Figure 1E). Likewise, while linear models scored artificially well on Maximum Mean Discrepancy (MMD) against raw zero-inflated old cells, the VAE achieved significantly superior alignment against a VAE-reconstructed (artifact-reduced) reference (Microglia VAE-reconstructed-reference MMD = 0.066 *±* 0.026 vs. PCA = 0.131 *±* 0.093, *p <* 0.01; Figure 1F). This trend was conserved in HSCs. Together, these metrics indicate that the VAE reduces structural dropout effects while preserving age-associated heterogeneity in the inferred manifold. We do not claim the VAE uniquely recovers an absolute biological ground truth; absent orthogonal modalities (e.g., scATAC-seq), our results instead indicate a consistent, technical-artifact-reduced representation useful for comparative, hypothesis-generating analysis.

### 2.2. VAE Captures a Transferable Aging Direction but Reveals Decoder Limits in Cross-Sex Topology

We stress-tested topological flexibility with out-of-distribution (OOD) validation on an unseen intermediate age and an unseen sex. When the 24-month vector was interpolated (*α ≈* 0.714) to predict the 18-month within-sex state, the VAE accurately mapped the intermediate trajectory, maintaining high macroscopic and topological alignment with the *in vivo* hold-out (Microglia *R*^2^ = 0.960 *±* 0.007, VAE-reconstructed-reference MMD = 0.099 *±* 0.078) and robustly preserving core aging heterogeneity across both tissues (SF 8).

For cross-sex generalization, we applied the male-derived vector to a young female baseline. The true aging vectors showed distinct cross-sex alignment: Microglia exhibited moderate positive cosine similarity (cos(*θ*) = 0.546 *±* 0.105), whereas HSCs were nearly antiparallel (cos(*θ*) = *−*0.228 *±* 0.038), indicating highly divergent, orthogonal sex-dimorphic shifts (SF 9F). Both linear baselines and the VAE independently maintained sex-dimorphic spatial separation (SF 9I), validating the underlying sex-specific baseline. Despite strong sex-dimorphic divergence in HSCs, the VAE captured a transferable directional component of aging, significantly outperforming linear baselines in transcriptomic shift concordance (HSC *R*_*shift*_: VAE = 0.261 *±* 0.094 vs. PCA = 0.129 *±* 0.091, *p <* 0.0001; SF 9B). This directional transfer, however, does not imply full topological generalization, as detailed below.

However, this revealed a topological limitation of generative decoding under extreme divergence. In moderately aligned Microglia, the VAE decoder generalized well, achieving tighter alignment than PCA (VAE-reconstructed-reference MMD: VAE = 0.112 *±* 0.065 vs. PCA = 0.182 *±* 0.112, *p <* 0.0001). In the highly orthogonal HSCs, forcing a female cell through a male-trained latent bottleneck distorted sex-specific network wiring, and the decoder-free PCA, operating on algebraic shifts, retained the original female topology more faithfully (HSC VAE-reconstructed-reference MMD: VAE = 0.689 *±* 0.157 vs. PCA = 0.572 *±* 0.083, *p <* 0.0001; SF 9G). Thus, while the directional vector of aging is conserved across sexes, accurate topological simulation across dimorphic tissues may require sex-specific decoders.

### 2.3. Bidirectional Validation: VAE Advantages Persist in Reverse Simulation

To test whether the GeroVector represents a coherent bidirectional axis rather than a one-directional fit, we applied the negated trajectory (*−***g**) to held-out old cells and benchmarked the reverse-simulated young profiles (GeroSimYoung) against the held-out young reference using the identical metric suite (Figure 2). Because the negated vector trivially displaces the population toward the young centroid, we focused on distributional metrics not guaranteed to transfer across directions. The full forward-direction pattern was recapitulated in reverse: the VAE again achieved significantly stronger artifact compression than the linear baselines and preserved trajectory heterogeneity, while ALS collapsed to near-zero shift variance. Critically, against the VAE-reconstructed (artifact-reduced) young reference, the VAE attained the lowest MMD in both tissues, whereas linear models scored deceptively well only against the raw young target. These results indicate that the VAE’s distributional advantages are direction-symmetric, supporting the interpretation that the GeroVector captures a genuine, traversable aging axis. The partial gene-level overlap between forward and reverse signatures, analyzed in detail below, reveals that the replicative/identity program is more bidirectionally reversible under linear latent perturbation than the downstream inflammatory cascade, concordant with the “primacy of identity loss” architecture.

**Figure 2.**
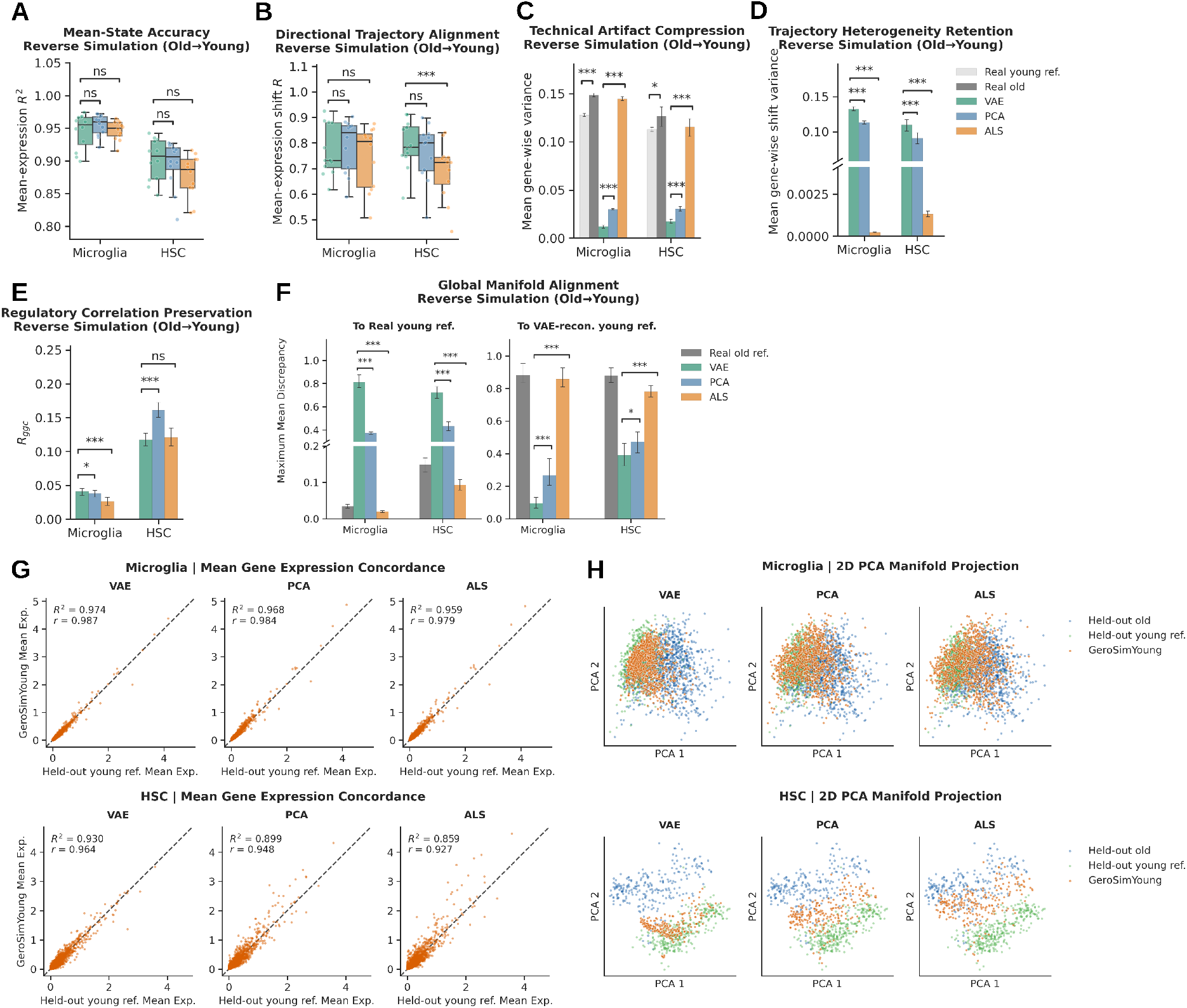
Reverse Simulation (Old*→*Young): VAE advantages are direction-symmetric. Reverse-direction benchmarking generated by applying the negated GeroVector (*−***g**) to held-out old cells and evaluating GeroSimYoung against the held-out young reference across 12 LOPO folds; panels correspond to Figure 1. Because the reverse trajectory applies the negated forward vector, population-level movement toward the young state is expected by construction; the informative comparisons are the distributional metrics, in which the VAE significantly outperforms linear baselines against the artifact-reduced reference. Panels (G,H) display representative simulations using the 3 9 M/24 60 M pair for microglia (fold 2) and 3 8 M/24 58 M for HSCs (fold 5). Significance: paired Wilcoxon signed-rank test with BH-FDR correction (* *p <* 0.05, ** *p <* 0.01, *** *p <* 0.001, ns: non-significant).

### 2.4. VAE-Derived Aging Signatures Are Highly Specific and Biologically Relevant

Applying the strict 12-fold LOPO consensus threshold yielded a reproducible signature of 38 microglial and 61 HSC consensus aging targets (GeroTargets; Supplementary Data; representative targets in Table 1). These computationally derived targets closely aligned with known hallmarks of aging and with our upstream GeroRegulator and Gene Set Enrichment Analysis (GSEA) predictions. In HSCs, the collapse of cell-cycle and replication pathways corresponded with consensus down-regulation of the minichromosome maintenance (MCM) helicase complex (*Mcm2/3/5/6*) and DNA polymerase (*Pold1*),[20, 21] while hallmark inflammatory myeloid skewing appeared as up-regulation of Major Histocompatibility Complex (MHC) class II molecules (*H2-Aa, H2-Eb1*) and *Cd74*.[22] In microglia, the model identified consensus down-regulation of the Activator Protein 1 (AP-1) complex (*Jun, Junb*)[23] alongside upstream cytokine/Janus kinase–signal transducer and activator of transcription (JAK–STAT) receptors (*Jak1, Lifr, Il13ra1*),[24–28] together with the classical microglial aging signature of up-regulated *Apoe* and MHC class I/II molecules (*H2-K1, H2-D1, H2-Ab1, Cd74*).[29–33]

**Table 1.** High-Confidence Consensus GeroTargets Identified by the VAE. Aging signatures defined using a strict 12*/*12 cross-validation threshold (38 microglial, 61 HSC targets). Representative genes are grouped by core biological function; complete lists in Supplementary Data.

| Cell type<br>(Tissue) | Trajectory | Representative Consensus GeroTargets |
| --- | --- | --- |
| Microglia<br>(Brain myeloid) | Up-regulated | **Inflammatory/MHC Response:** *Tnfsf13b*, *H2-Q6*, *H2-K1*, *H2-D1*, *H2-Ab1*, *Cd74*, *Apoe*, *Lyz2* |
|  | Down-regulated | **Upstream Receptor/Signaling Cascade:** *Lifr*, *Il13ra1*, *Jak1*, *Jun*, *Junb*<br>**Immune/Phagocytic Homeostasis:** *Rab31*, *C3ar1* |
| HSCs<br>(Marrow) | Up-regulated | **Myeloid Skewing/Inflammation:** *H2-Aa*, *H2-Eb1*, *H2-Ab1*, *Cd74*<br>**Metabolic/Stress Response:** *Apoe*, *Dusp1*, *Egr1*, *Fos*, *Junb*, *Jun*, *Nfkbia* |
|  | Down-regulated | **DNA Replication/Cell Cycle:** *Mcm2*, *Mcm3*, *Mcm5*, *Mcm6*, *Pold1*, *Mki67*<br>**Stemness/Progenitor Markers:** *Flt3*, *Kit*, *Mpo* |

GSEA revealed a profound divergence in how linear versus non-linear models capture age-related shifts. In microglia, while all models captured the MHC response, the VAE additionally prioritized lineage-specific upstream terms, myeloid-centric pathways (Osteoclast differentiation, Leishmaniasis) and, uniquely, the Cytokine-cytokine receptor interaction (odds ratio [OR] = 6.5) and JAK–STAT (OR = 10.2) cascades, by capturing upstream receptors (*Lifr, Il13ra1, Jak1*) that linear baselines missed or de-prioritized (ST 1 and SR 2.1). In HSCs, the same paradigm held: the VAE retained the inflammatory component while prioritizing putative upstream drivers of stem-cell exhaustion as its top hits, isolating DNA replication with a far higher odds ratio (OR = 84.6 vs. PCA 52.0, ALS 41.5; capturing the MCM complex and *Pold1*) and elevating Hematopoietic cell lineage and pre-leukemic transformation pathways through the loss of *Kit, Flt3*, and *Mpo* (ST 2 and SR 2.2). Thus the VAE’s non-linear manifold prioritizes core hallmarks of replicative senescence and progenitor exhaustion alongside broader inflammatory remodeling, mitigating over-reliance on MHC-dominated downstream signals.

### 2.5. Master Regulators and the Three Conserved Pillars of Cellular Aging

A reverse-directed Personalized PageRank (PPR) algorithm seeded by the 12/12 consensus GeroTargets, followed by Kneedle thresholding and tissue-adaptive topological filters, reconstructed the core tissue-specific regulator–target networks (GeroNetworks; Table 2, Figure 3). In Microglia, the algorithm prioritized *Rela, Nfkb1*, and *Irf1* connected to MHC targets (*Cd74, H2-D1*), a sexually dimorphic core (*Esr1, Ar*) converging on the AP-1 complex and *Lifr*, and identity regulators (*Spi1, Tal1*) mapping to loss of homeostatic receptors (*Rab31, C3ar1*). In HSCs, the *E2f* /*Tp53* /*Myc* modules drove the downregulation of the MCM complex and *Mki67*, while myeloid master factors (*Spi1, Cebpa*) and stress hubs (*Creb1*) drove up-regulated inflammatory targets (*Cd74, Nfkbia, Cd9*). This structural alignment between mathematically derived GeroRegulators and empirical GeroTargets supports the VAE’s capacity to reconstruct tissue-specific regulatory hierarchies of cellular senescence.

**Table 2.**
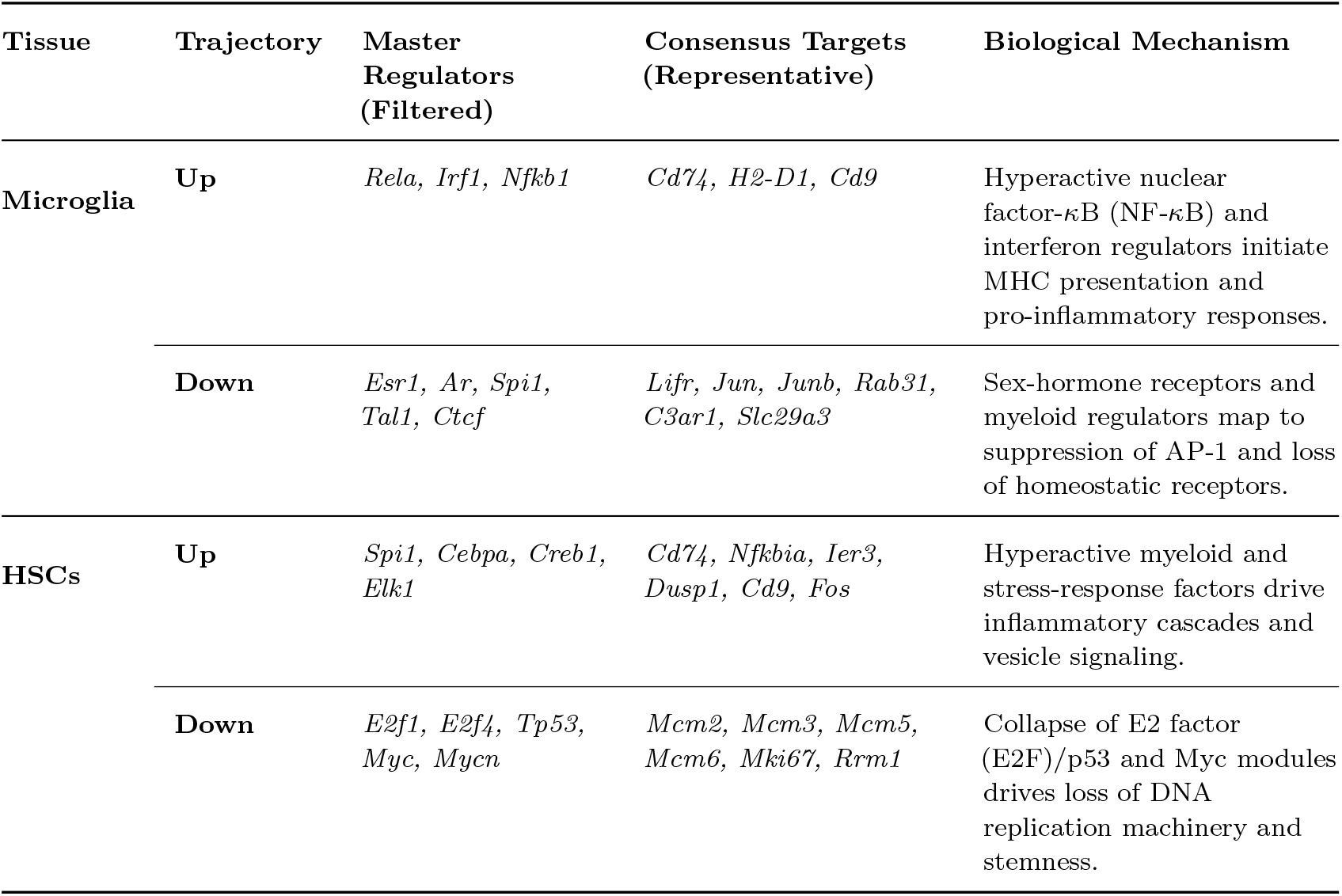
Mechanistic Alignment of VAE-Derived GeroRegulators and Consensus GeroTargets. Upstream master regulators (tissue-adaptive thresholds) align with the stringent consensus targets (12*/*12 folds) they putatively regulate, reconstructing robust regulatory hierarchies across distinct functional aging pillars.

**Table 3.**
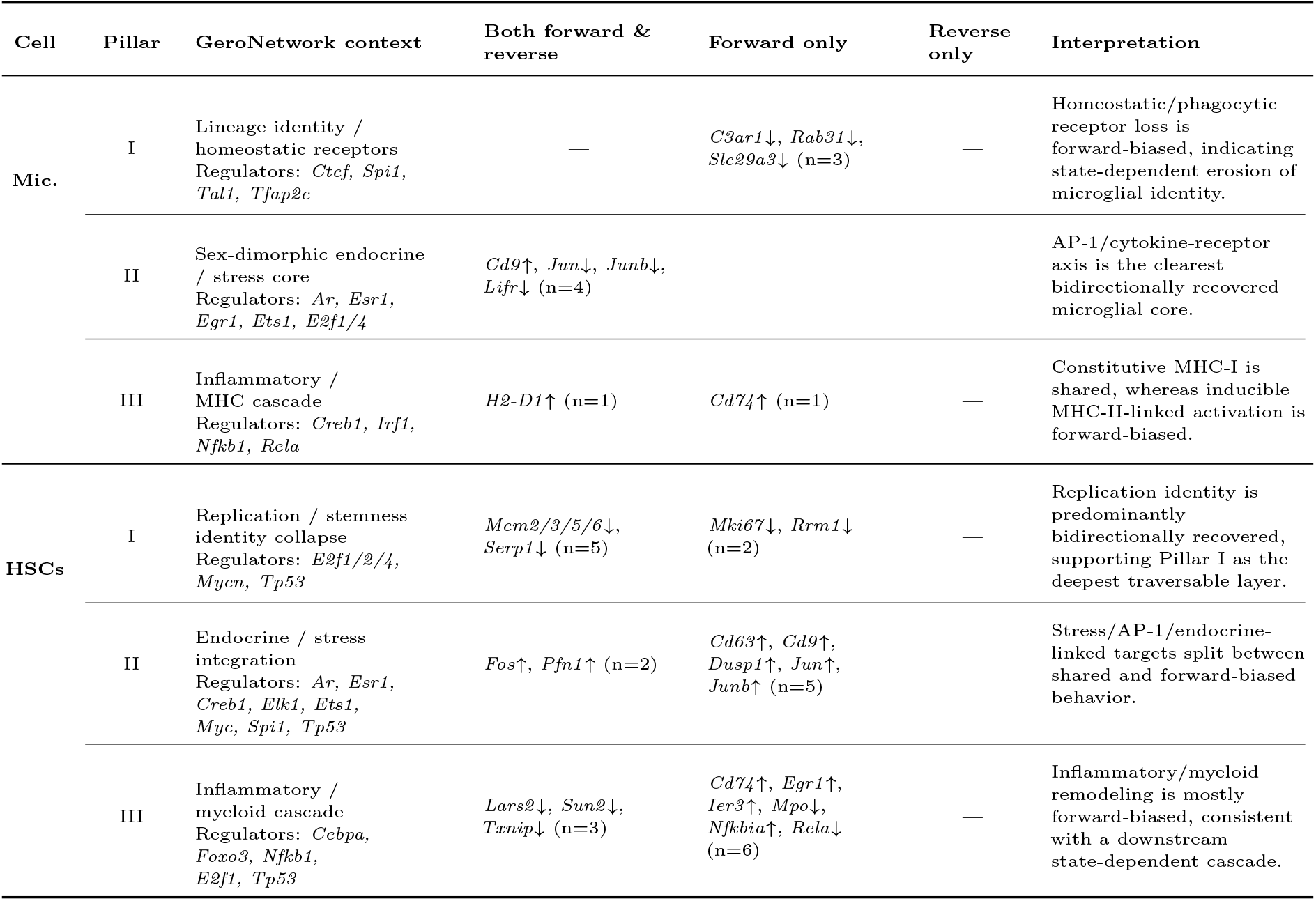
Forward–reverse recovery of GeroNetwork three-pillar targets. Consensus GeroTargets were classified as recovered in forward simulation only, reverse simulation only, or both directions after expressing both gene-effect signatures on the common young*→*old aging axis. Rows show only targets assigned to the displayed three-pillar GeroNetwork core; peripheral non-pillar consensus targets are listed in Supplementary Data. Pillar membership denotes GeroNetwork topology, whereas forward–reverse class denotes simulation recovery behavior.

**Figure 3.**
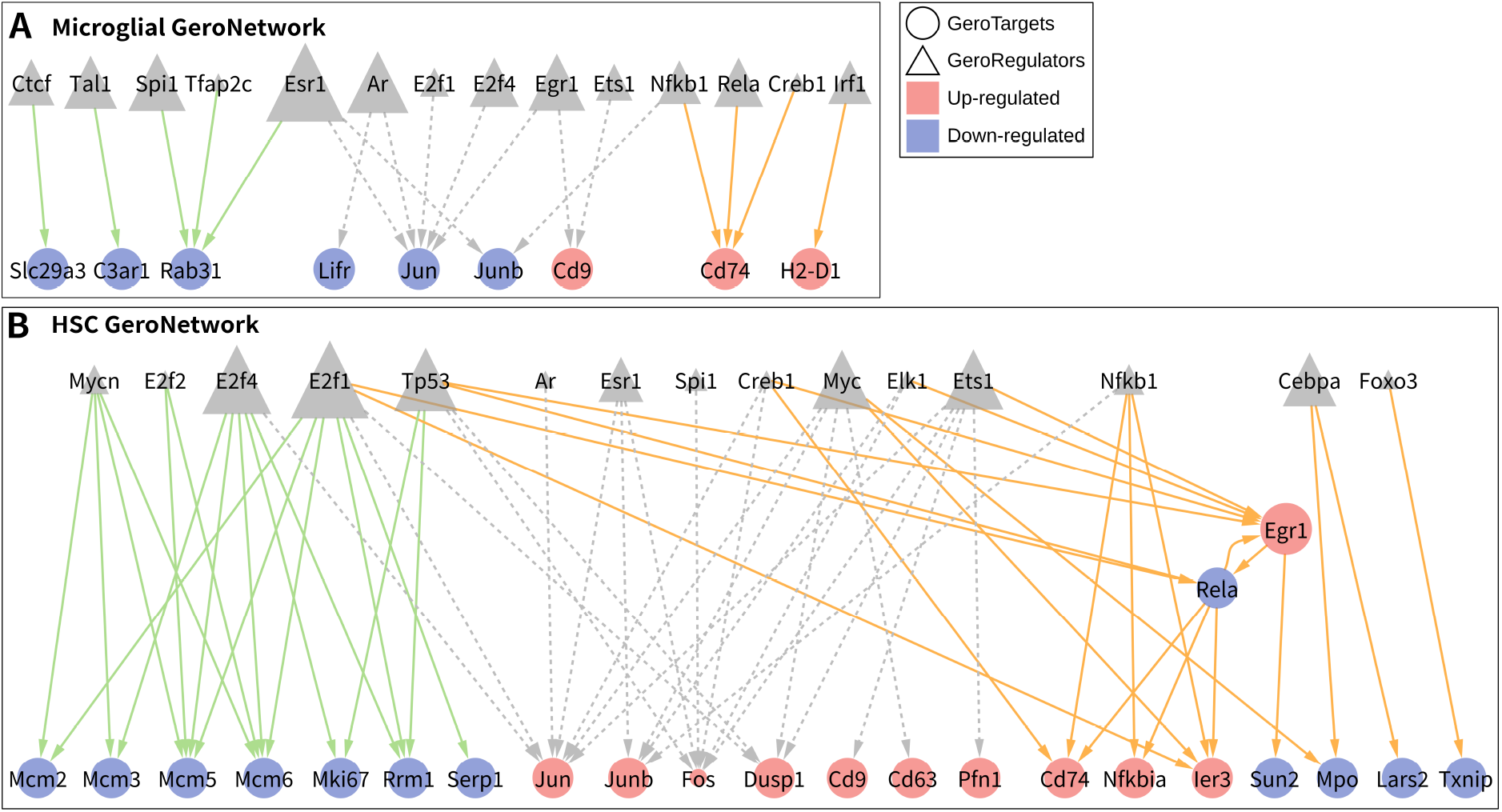
Topological Master Regulators Map Tissue-Specific and Shared Aging Trajectories. Hierarchical regulatory networks for (A) Microglia and (B) HSCs constructed from strict consensus GeroTargets (circles); upstream Master GeroRegulators (triangles) are scaled by topological score. Layout and edge styling delineate three pillars: lineage-specific homeostasis and stemness collapse (solid green), the centrally conserved sexually dimorphic stress core (dashed gray), and the systemic inflammatory/myeloid cascade (solid orange).

Structural analysis of the directed GeroNetworks (Figure 3) revealed that the hierarchy segregates into three conserved functional pillars across both tissues, mapping the flow from upstream regulators to downstream targets. **Pillar I (Lineage-Specific Homeostasis and Identity Collapse)** encapsulates loss of cellular identity: in microglia, suppression of identity factors (*Spi1, Tal1, Tfap2c, Ctcf*) drives downregulation of phagocytic/sensory targets (*Rab31, C3ar1, Slc29a3*); in HSCs, *E2f1/E2f4* hubs with *Tp53* /*Mycn* drive a coordinated shutdown of the MCM complex and *Mki67*, representing replicative senescence. **Pillar II (The Sexually Dimorphic Stress Core)** is a dense central integration hub orchestrated by sex-hormone receptors (*Esr1, Ar*) converging on AP-1 (*Jun, Junb, Fos*); strikingly, the regulatory sign of this conserved core inverts by tissue, suppressive in microglia (downregulating AP-1 and the neuroprotective *Lifr*) versus hyperactivating in HSCs (upregulating AP-1, *Dusp1*, and vesicle markers *Cd9* /*Cd63*). Because these hubs were inferred from a male-derived trajectory, their centrality provides a mechanistic hypothesis for the cross-sex OOD degradation observed in HSCs. **Pillar III (The Systemic Inflammatory and Myeloid Cascade)** captures inflammaging: in microglia, *Irf1* /*Nfkb1* /*Rela* link directly to MHC presentation (*H2-D1, Cd74*), whereas in HSCs a multi-tiered cascade routes myeloid/stress regulators (*Cebpa, Spi1, Foxo3*) through an intermediate *Egr1 →Rela* feed-forward loop to the terminal inflammatory state, providing topological evidence that aging stem cells undergo aberrant myeloid priming before a terminal inflammatory phenotype.

### 2.6. Forward–Reverse Target Symmetry Reveals a Bidirectional Identity Core

The GeroNetwork hierarchy predicts that lineage/replication identity programs (Pillar I) act as deep upstream drivers while the inflammatory/MHC cascade (Pillar III) emerges as a downstream, state-dependent consequence. A direct gene-level test is whether the deep identity programs are more bidirectionally traversable than the downstream inflammatory programs.

For gene-level comparison, forward and reverse signatures were both expressed on the common young*→*old aging axis: forward effects were computed as simulated aged profiles (GeroSimOld) minus real young, whereas reverse effects were computed as real old minus GeroSimYoung. Under this convention, a bidirectionally recovered aging gene is expected to retain the same up- or down-regulated sign in both analyses.

Using this aging-axis convention, the VAE shared a sign-consistent but incomplete core (22 shared targets in microglia, 29 in HSCs; 38% Jaccard in both), with no sign-inconsistent genes detected in any model, tissue, or condition, confirming a sign-coherent bidirectional axis (Figure 4A).

**Figure 4.**
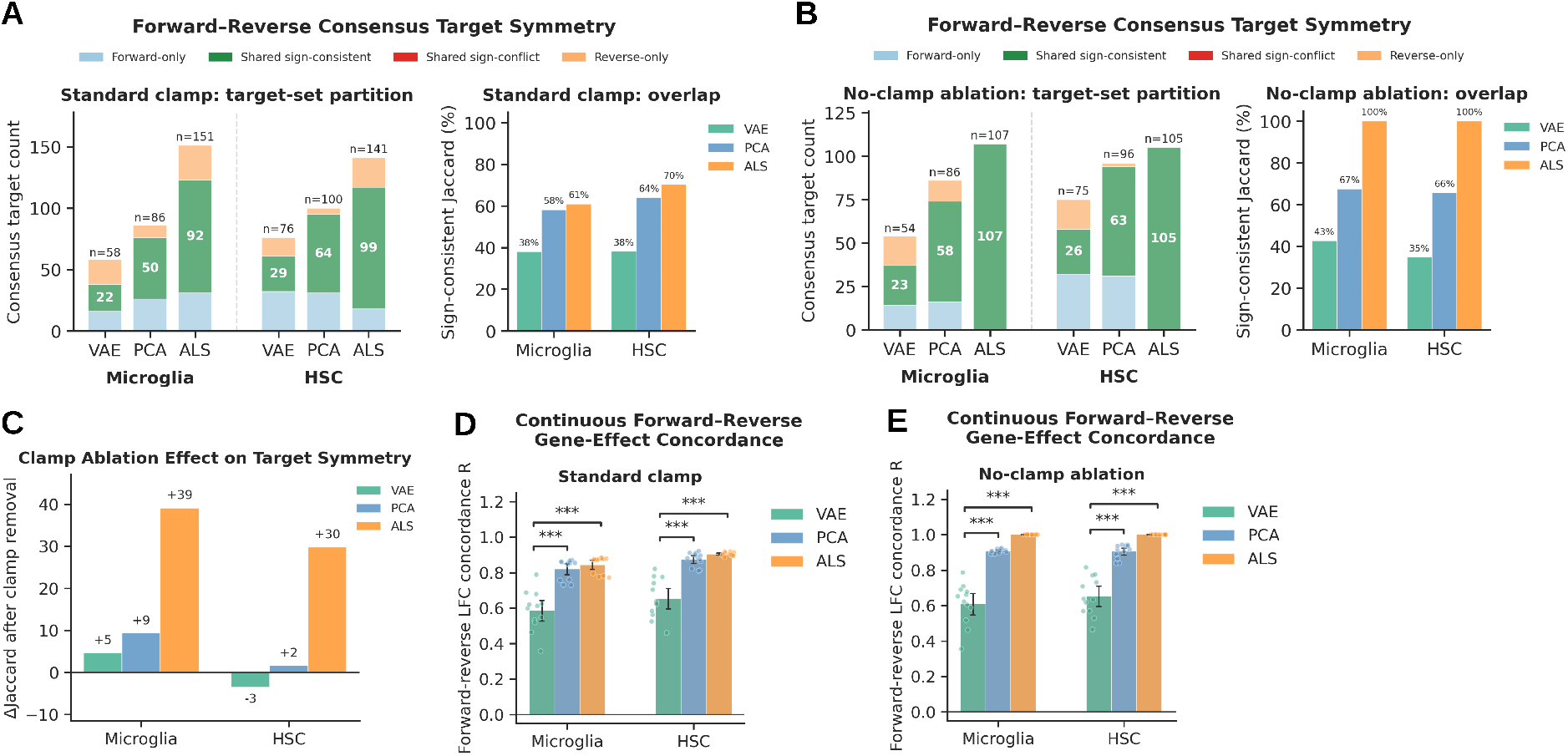
Forward–reverse target symmetry and clamp ablation reveal VAE-specific state-dependent manifold traversal. Forward young*→*old and reverse old*→*young consensus targets were compared after expressing both gene-effect signatures on the common young*→*old aging axis. Forward effects were defined as GeroSimOld minus real young, and reverse effects were defined as real old minus GeroSimYoung. (A) Under the standard clamped pipeline, ALS, PCA, and the VAE each showed sign-consistent overlap, but the VAE retained a smaller shared core and a larger direction-specific component (counts shown within each bar; *n* above each bar indicates the union size). (B) Removing the non-negativity clamp restored complete same-axis forward–reverse symmetry in ALS, produced a modest increase in PCA overlap, and minimally changed VAE overlap. (C) ΔJaccard following clamp removal. ALS recovered to complete same-axis symmetry (+39 percentage points Microglia; +30 HSC), PCA recovered partially (+9, +2), and the VAE was essentially unchanged (+5, *−*3), identifying intrinsic non-linear decoding as the dominant source of the VAE’s direction-specific recovery. (D–E) Continuous threshold-free forward–reverse log_2_FC concordance (Pearson *r*) across all HVGs per LOPO fold, under standard clamp (D) and no-clamp ablation (E). Dots, per-fold values; bars, mean *±* SD over 12 folds. ALS reached *r* = 1.00 after clamp removal, PCA approached *r ≈* 0.91, and the VAE remained at *r ≈* 0.60–0.65, consistent with the discrete-set result and confirming that the asymmetry is not a consensus-threshold artifact. The no-clamp condition permits negative reconstructed expression values and is used only as a mechanism-isolating ablation, not as a biologically realizable simulated transcriptomic state. Across all 12 conditions (3 models *×* 2 tissues *×* 2 clamp settings), no shared sign-inconsistent genes were detected after aging-axis alignment, confirming the sign coherence of the learned bidirectional aging axis. Jaccard overlap is defined as shared sign-consistent genes divided by the union of forward and reverse consensus target sets.

This partial overlap is consistent with state-dependent decoder geometry rather than simple inferior reversibility: two control analyses (a no-clamp ablation and a threshold-free continuous log_2_FC concordance across all Highly Variable Genes [HVGs]) rule out both output rectification and consensus thresholding as the cause and isolate intrinsic non-linear, state-dependent decoder geometry as the source (Figure 4; full mechanistic dissection and statistics in SR 2.3).

Biologically, the VAE partitions matched the GeroNetwork hierarchy: the HSC shared bidirectional core was dominated by replication/progenitor-identity machinery (MCM complex, *Pold1, Flt3*), precisely the Pillar I program anchored by *E2f* /*Tp53* /*Mycn*, while the forward-only set was dominated by the Pillar III inflammatory cascade (MHC class II, AP-1/immediate-early/NF-*κ*B regulators, *Apoe*). Across both tissues, constitutive MHC class I components were bidirectionally reversible whereas inducible MHC class II was forward-specific. This dissociation provides convergent gene-level support for the primacy of identity loss: Pillar I identity/replication programs constitute the deep, bidirectionally traversable core of the learned trajectory, while Pillar III programs are state-dependent downstream consequences.

## 3. Discussion

Biological aging is inherently stochastic and heterogeneous; individual cells within the same tissue do not age at identical rates or follow identical transcriptomic trajectories. Simulating how a young population transitions into an aged state is therefore essential for identifying the upstream regulators of senescence, yet traditional methods struggle with the severe technical noise and dropouts of scRNA-seq. We established the GeroEngine as a robust *in silico* platform to overcome these limitations.

Our empirical evaluation confirms that absolute transcriptomic variance is significantly higher in the true aged state than in the young baseline.[9] This age-associated variability poses a computational challenge: standard linear models may propagate source-cell sparsity forward, artificially maintaining variance without distinguishing technical carryover from reproducible biological shifts. By passing profiles through a non-linear latent bottleneck, the VAE reduces scRNA-seq-specific artifacts while age-related biological heterogeneity is retained as part of the phenotype. We emphasize that the shared directional component of aging can be represented in a more compact latent manifold, but the dispersion of *in vivo* aged populations remains biologically meaningful and should not be interpreted as noise removed by the model.

This capacity to reduce technical artifact carryover while estimating a reproducible directional trajectory lets us conceptualize aging not strictly as a programmed clock, but as a cumulative erosion of function. We propose that the heightened variance of the aged state reflects the molecular footprint of continuous exposure to the cumulative environmental exposome.[34] Over a lifespan, these stressors act as chronic triggers that progressively degrade the epigenome, desensitize receptors, and disrupt signaling cascades;[35] while the accumulation of damage is stochastic, the phenotypic outcome is directional, the cell loses its youthful identity and specialized functions. The ultimate objective of anti-aging therapeutics, therefore, cannot merely be brute-force suppression of downstream stress variability or inflammation, but must aim to systematically restore eroded cellular identity.

The VAE-derived GeroNetworks provide a computational framework to hypothesize such interventions, mapping the inferred hierarchical flow from upstream regulators to downstream effectors and unveiling several conserved principles. First, they establish the **Primacy of Identity Loss**: the tripartite architecture delineates the inferred primary drivers (Pillar I) from likely downstream consequences (Pillar III inflammaging), with the most topologically dominant regulators linked not to generic inflammation but to the dismantling of the cell’s core function. Second, they reveal **Structural Conservation with Directional Divergence**: the manifold relies on a stable geometry across distinct germ layers, using the same sexually dimorphic endocrine hubs (*Esr1, Ar* ; Pillar II), yet the regulatory sign of this conserved core is strictly context-dependent. Finally, the centrality of endocrine (*Esr1, Ar*) and systemic stress (*Nfkb1, Myc*) regulators bridges cell-autonomous senescence and systemic organismal aging, consistent with the hypothesis that intracellular deterioration is orchestrated by shifts in systemic endocrine signals.

Our bidirectional analysis showed the VAE exhibits substantially lower forward–reverse gene overlap than the linear baselines, a gap that persists after removing both the non-negativity rectification and the consensus threshold. We interpret this lower overlap not as inferior reversibility but as the diagnostic signature of a model that explicitly encodes state-dependent biology: ALS is state-independent by construction, PCA introduces mild state dependence through lossy projection, and the VAE is strongly state-dependent because a fixed latent displacement **g** produces different gene-space shifts depending on the starting cellular state (full state-dependence analysis in SDi 3.1). Biologically, once a cell has acquired the aged inflammatory state, negating the population vector does not deterministically restore every gene, because the inflammatory program engages downstream of the more state-independent identity machinery. The deeper a program lies in the hierarchy, the more bidirectionally reversible it appears, precisely what the consensus partition shows, with the MCM complex, *Pold1*, and *Flt3* forming the shared core while the MHC class II/AP-1/NF-*κ*B module is recovered only forward. This reframes the partial overlap as a depth-of-hierarchy readout and prioritizes Pillar I master regulators (*E2f, Mycn*; *Spi1, Tal1*) as candidates more likely to yield broad, coherent reversal than downstream inflammatory hubs alone. We interpret reverse simulation throughout as a computational analog of rejuvenation-like latent traversal, not as evidence of validated biological rejuvenation.

Guided by these principles, the Topological GeroRegulators emerge as prioritized, experimentally testable candidate intervention nodes at the points where cumulative damage may translate into functional collapse. In HSCs, these results nominate the *E2f* and *Mycn* modules as candidates for testing whether replication-associated identity can be restored; in Microglia, they nominate *Spi1, Tal1*, and *Rela*-linked modules for testing sensory-identity and inflammatory-state modulation. While strictly computational and requiring experimental validation, these hubs offer a prioritized hypothesis space for testing whether tissue-specific pharmacological or epigenetic modulation can restore aspects of youthful cellular function.

## 4. Methods

### 4.1. Overview of the GeroEngine Pipeline

Following single-cell preprocessing, we employed a modified 12-fold LOPO cross-validation strategy in which all possible combinations of one young and one old mouse were iteratively held out as the test set while the remaining cross-sectional cohort was used for training (full preprocessing, feature selection, and cross-validation design in SM 1.1). For each fold, a deep VAE was trained on the background cohort to calculate a fold-specific population-level latent aging vector, GeroVector 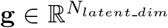 Applying **g** to the latent representations of hold-out young cells 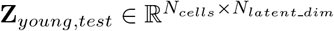 produced predicted latent states that were decoded back into gene-expression space to generate *in silico* mouse-matched longitudinal profiles GeroSimOld 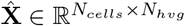 (Figure 5).

**Figure 5.**
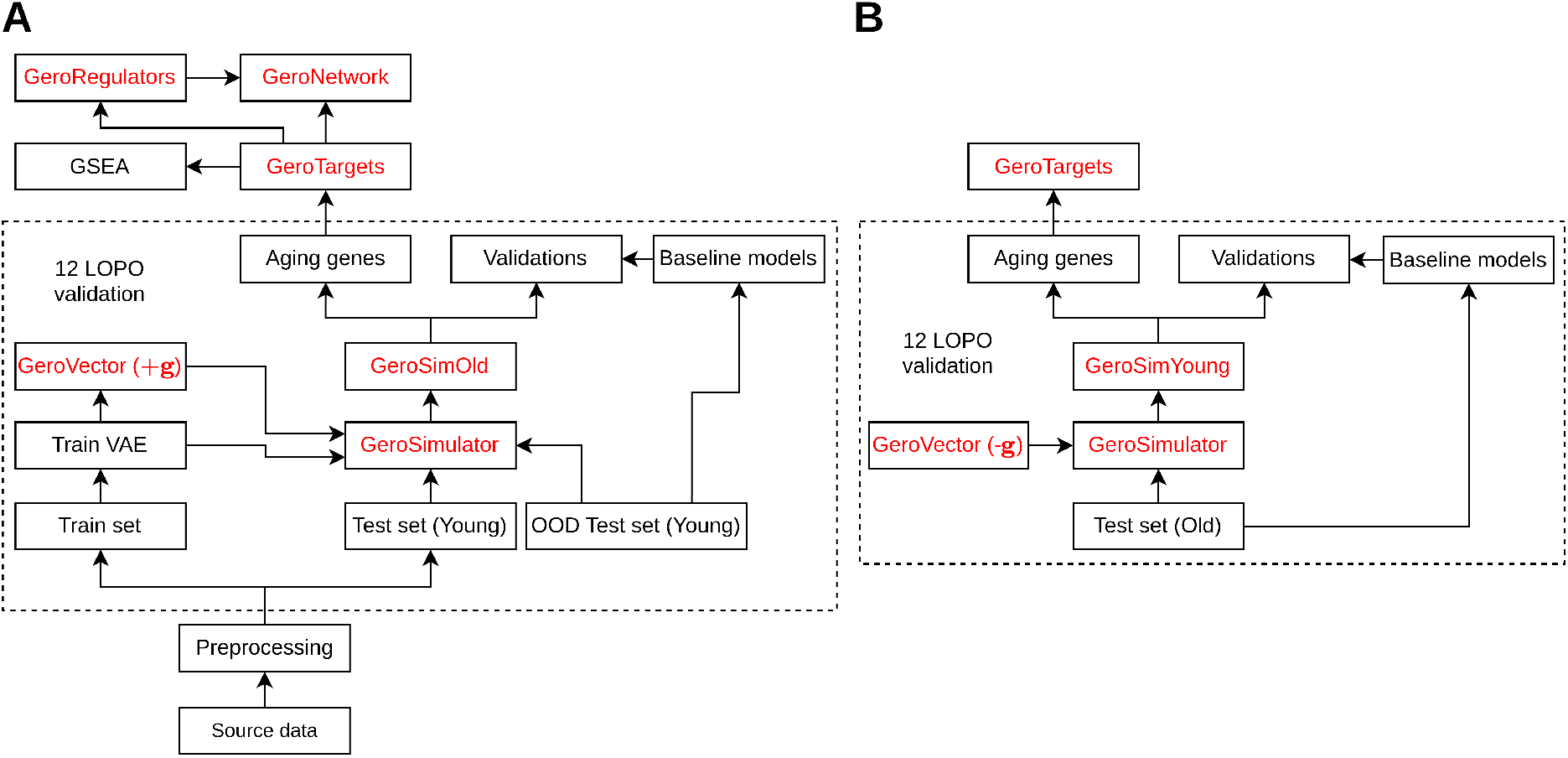
Overview of the GeroEngine computational pipeline. **(A)** Forward (young*→*old) simulation. Source transcriptomic data undergoes preprocessing and is split into training, test, and Out-Of-Distribution (OOD) sets. Within a stringent 12-fold Leave-One-Pair-Out (LOPO) cross-validation framework (dashed box), a Variational Autoencoder (VAE) is trained on the background cohort to establish a latent trajectory of aging (GeroVector, +**g**). The GeroSimulator applies this vector to held-out young test cells (and OOD young cells) to generate simulated aged profiles (GeroSimOld), benchmarked against linear baseline models in the validation step. Downstream inference extracts high-confidence consensus GeroTargets, which are subjected to Gene Set Enrichment Analysis (GSEA) and topological network extraction to identify upstream GeroRegulators, reconstructing tissue-specific GeroNetworks. **(B)** Reverse (old*→*young) simulation. Using the same fold-specific trained models, the negated trajectory (*−***g**) is applied to held-out old test cells to generate reverse-simulated young profiles (GeroSimYoung), benchmarked against the same baseline models and processed through the identical aging-gene and consensus-target pipeline to assess sign-consistent recovery on the common young*→*old aging axis. Red text denotes core modules and latent outputs generated by the GeroEngine; black text indicates standard processing and baseline steps.

After quantitative validation of the predicted state space, differential expression analysis was performed on paired real-young and GeroSimOld profiles. To eliminate inter-individual batch effects, differentially expressed targets were intersected across all 12 folds, establishing a definitive consensus aging signature (GeroTargets). Upstream genes (GeroRegulators) were inferred using a reverse-directed Personalized PageRank (PPR) algorithm[36, 37] mapped onto literature-curated regulatory graphs (OmniPath[38] and DoRothEA[39]). Topological GeroRegulators were established by intersecting the dynamically shifting potential aging gene pool with the PPR-derived regulatory hubs, and integrated with GeroTargets to construct a unified GeroNetwork mapping the putative regulatory hierarchy of single-cell aging.

### 4.2. Data, VAE Architecture, and Trajectory Generation

Raw count matrices were acquired from the FACS-sorted Tabula Muris Senis atlas.[40] The primary, sex-controlled cohort was restricted to 3-month-old (Young) and 24-month-old (Old) male mice, comprising 6434 microglial cells and 2113 HSCs after quality control. Two independent OOD validation cohorts were established: an unseen-age cohort (18-month males) and an unseen-sex cohort (3- and 18-month females). Highly Variable Genes (HVGs) were identified on the raw male cohort using a standardized variance cutoff of 1.3 (3182 microglial, 2908 HSC transcripts); all cohorts were then library-size normalized and log-transformed, and subsetted to the primary HVG feature space to prevent leakage. A global random seed of 42 ensured reproducibility. Detailed preprocessing, quality-control thresholds, cohort sizes, and the LOPO design are provided in SM 1.1.

We adopted the scGen VAE framework, implemented via `scvi-tools`,[10, 19, 41, 42] extended with strict LOPO consensus filtering and individual-agnostic trajectory weighting. The encoder and decoder each comprise two 800-unit hidden layers mapping to a 100-dimensional latent space, with the decoder reconstructing in the ln(*x* + 1) domain (negative values clamped to zero at inference). A hyperparameter ablation across latent dimensionalities and KL penalties showed exceptional robustness, with most configurations yielding *R*_*shift*_ *>* 0.70 on unseen folds (SF 3); we selected a balanced architecture (*N*_*latentdim*_ = 100, *β*_*KL*_ = 10^−3^) that avoids the heavy regularization which artificially crushes transcriptomic variance, so the simulated transcriptomic contraction remains an emergent property rather than a tuning artifact. Full architecture, optimization, and ablation details are in SM 1.2.

The GeroVector **g** was computed via a strict mouse-balanced averaging algorithm in which the unweighted mean latent vector was first computed per mouse and then averaged across mice, ensuring every subject contributed equally to the systemic aging signature (SA 1). GeroSimOld profiles were then generated by encoding hold-out young cells, adding **g** in latent space, and decoding with non-negativity clamping (SEq 1, SEq 2).

Forward and reverse simulations were generated by applying opposite latent displacements while using the same fold-specific encoder, decoder, and training-derived GeroVector. In the forward young*→*old direction, held-out young cells were encoded, shifted by +**g**_*fold*_, and decoded to generate GeroSimOld:

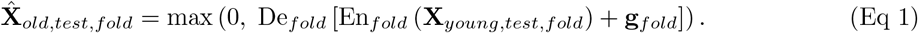

In the reverse old*→*young direction, held-out old cells were encoded, shifted by *−***g**_*fold*_, and decoded to generate GeroSimYoung:

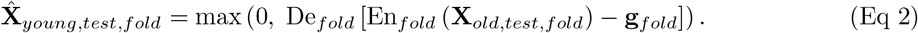

### 4.3. Linear Baselines, Reverse and OOD Simulation

To evaluate the necessity of non-linear inference, we established two deterministic linear baselines: Algebraic Linear Shift (ALS), which adds a mouse-balanced expression-space aging vector directly to young test cells, and PCA, which applies a mouse-balanced aging vector within a 100-component linear subspace before inverse-transforming. All aging vectors were computed strictly within training partitions using mouse-balanced means to prevent leakage and recovery-rate bias (SM 1.3).

To test whether the learned trajectory behaves as a coherent bidirectional axis, we performed reverse-direction (Old*→*Young) simulation using the same fold-specific trained models and training-derived vectors; no model was retrained and no held-out cells were used to recompute trajectories. Reverse VAE predictions were generated by encoding held-out old cells, applying the negated vector *−***g**, and decoding, yielding GeroSimYoung profiles (PCA and ALS reverse simulations defined analogously; SM 1.4). Because reverse simulation explicitly applies the negated forward vector, population-level movement toward the young centroid is expected by construction; we therefore interpreted reverse simulation using distributional metrics, continuous gene-effect concordance, and forward–reverse consensus-target overlap rather than centroid movement alone.

For gene-level GeroTarget extraction and forward–reverse comparison, both simulated directions were expressed on the common young*→*old aging axis. Forward aging effects were defined as the pseudo-bulked difference between GeroSimOld and the paired real-young source state:

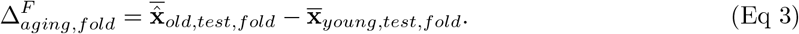

Reverse effects were aligned to the same young*→*old axis by comparing the real-old source state against GeroSimYoung:

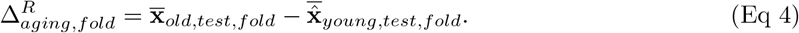

Thus, positive and negative GeroTarget directions always indicate up- or down-regulation on the young*→*old aging axis, regardless of whether the simulated profile was generated in the forward or reverse direction. Bidirectionally shared genes were considered sign-consistent when 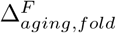 and 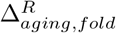 had the same direction after consensus filtering.

For OOD validation, the 24-month aging vector was linearly interpolated by *α* = (18 *−* 3)*/*(24 *−* 3) *≈* 0.714 to predict the unseen 18-month state, and the male-derived vector was applied to the young female baseline to stress-test cross-sex topological flexibility. For all OOD analyses, no PCA basis or trajectory vector was estimated from OOD evaluation cells; all model components were derived exclusively from the primary male LOPO training cohort, and OOD cells were used only as inputs transformed through these training-derived mappings (SM 1.6).

For each LOPO fold, the female aging direction was obtained by encoding the female OOD cells with that fold’s forward-trained VAE and computing the mouse-balanced old-young latent centroid difference **g**_*female*_, using the identical procedure used for the male vector **g**. Cross-sex alignment was then quantified as the cosine similarity cos(**g, g**_*female*_) per fold, yielding one value per fold (12 total; mean *±* SD), visualized as a bar chart with per-fold strip points.

### 4.4. Quantitative Evaluation Metrics

We assessed simulation fidelity using a suite of metrics evaluating macroscopic alignment, directional accuracy, technical-artifact compression, trajectory-heterogeneity preservation, and global manifold topology (full algebraic definitions and a consolidated visual summary in SM 1.7 and SF 7). Briefly: (i) the coefficient of determination (*R*^2^) on pseudo-bulked profiles measures population-level mean-state accuracy;[19, 43] (ii) the Pearson correlation of transcriptomic shifts (*R*_*shift*_) verifies that the model captures the true directionality of aging rather than static identity;[44] (iii) the mean of gene variances (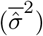) detects propagation of zero-inflation-associated technical artifacts, interpreted as reduced carryover only when accompanied by preserved shift variance and manifold fidelity;[8, 12] (iv) the mean of gene shift variances (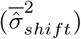) detects algorithmic mode collapse, with values near zero indicating a rigid uniform trajectory; (v) gene-gene co-expression concordance (*R*_*ggc*_) quantifies preservation of multivariate regulatory structure;[12, 45, 46] and (vi) Maximum Mean Discrepancy (MMD) with a multi-scale Gaussian RBF kernel grades global manifold alignment against both a raw *in vivo* reference and a VAE-reconstructed (artifact-reduced) reference.[47, 48]

These metrics must be interpreted holistically: because raw scRNA-seq data is highly zero-inflated, deterministic linear projections can achieve artificially high *R*^2^ and *R*_*ggc*_ simply by preserving the static technical artifacts of the starting population. Distinguishing true trajectory inference from artifact propagation therefore requires accurate dynamic directionality (high 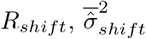) alongside reduced artifact carryover, non-collapsed heterogeneity, and convergence on a technical-artifact-reduced topological reference (low VAE-reconstructed-reference MMD).

### 4.5. Consensus Targets, Regulator Inference, and Statistics

A rank-based consensus approach was used to extract aging signatures. Pseudo-bulked young and predicted-old expressions were converted to log_2_ fold change, segregated by directionality with a strict zero baseline, and the top 100 up- and down-regulated genes ranked per fold; only genes ranking within the top 100 across all 12 folds (100% consensus) were finalized as GeroTargets. Downstream GSEA queried the KEGG 2019 Mouse and GO Biological Process 2021 libraries via gseapy with the full tissue HVG set as background.[49–53]

GeroRegulators were isolated through a sequential filtering pipeline on directed OmniPath/DoRothEA graphs: a reverse-directed PPR algorithm seeded by consensus GeroTargets prioritized upstream hubs, which were retained only if they intersected the active aging gene pool, passed structural cleaning (Kneedle elbow thresholding[54], UniProt/MGI symbol validation[55]), possessed a directed path of length *L ≤* 2 to GeroTargets, and met network-size-adaptive out-degree and PageRank thresholds (full pipeline and tissue-specific thresholds in SM 1.8 and SM 1.9). Networks were visualized in Cytoscape.[56] Because evaluations were strictly paired across the 12 LOPO folds, statistical significance between models was determined by paired Wilcoxon signed-rank tests with Benjamini-Hochberg FDR correction; data are mean *±* SD across folds (*a priori* thresholds: * *p <* 0.05, ** *p <* 0.01, *** *p <* 0.001, **** *p <* 0.0001).[57–60]

## 5. Limitations of the Study

Several limitations must be acknowledged. First, empirical validation was restricted to two cell types (microglia, HSCs) from a single murine atlas; future studies must validate the framework across broader solid-tissue architectures and human cohorts. Second, our benchmarking focused on the mathematical divergence between linear baselines and a VAE, and did not benchmark against the broader landscape of modern generative models (e.g., scVI[11] or multi-task frameworks), which future iterations should evaluate. Third, although the VAE embeds profiles into a non-linear manifold, the temporal perturbation itself is a strictly linear vector addition (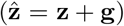); we therefore frame the trajectory as a first-order approximation, and future work should explore non-linear models such as Neural ODEs[61] directly in the latent space. Relatedly, because the reverse analysis applies the negated forward vector, population-level convergence is expected by construction, and the partial asymmetric overlap does not constitute evidence of biological rejuvenation. Finally, the GeroRegulators are computational predictions from curated databases; the conclusion that “erosion of cellular identity” is the primary driver of senescence remains a computationally generated hypothesis requiring *in vitro* and *in vivo* validation (e.g., targeted CRISPR-Cas9 perturbation).[62, 63]

## 6. Conclusion

Accurately modeling how a young cell transitions into an aged state is a foundational step toward prioritizing experimentally testable hypotheses for aging intervention. We addressed the physical (destructive sampling) and mathematical (shift-invariant linear models) limitations of longitudinal single-cell tracking by developing the GeroEngine. Through non-linear generative inference, we reduced dropout- and zero-inflation-associated technical artifacts and estimated reproducible transcriptomic shifts associated with senescence, while treating cumulative exposome-driven heterogeneity as part of the biological aging state rather than noise to be removed.

Our topological analysis reframes the molecular pathogenesis of aging. The tripartite GeroNetwork architecture supports a model in which a primary driver of senescence is the directional erosion of lineage-specific identity, replicative exhaustion in HSCs and homeostatic sensory collapse in microglia, rather than uniform decay or generic inflammation. The conserved, sexually dimorphic endocrine core (*Esr1, Ar*), inferred from male-derived trajectories, is consistent with a mechanistic bridge linking intracellular identity loss to systemic organismal signals. Forward–reverse traversal, analyzed after aligning both gene-effect signatures to the common young*→*old aging axis, demonstrated that the learned aging axis is sign-coherent but direction-specific: a shared identity/replication core was recovered in both directions, whereas inflammatory remodeling remained preferentially forward-recovered, providing convergent gene-level support for the primacy of identity loss. By redefining cellular aging as a loss of specialized identity, our framework suggests that future anti-aging strategies should test whether prioritizing upstream regulatory hubs can restore aspects of cellular homeostasis, rather than merely suppressing downstream inflammatory symptoms. All conclusions herein are strictly restricted to the analyzed murine cell types and datasets and should not be generalized to systemic human aging without rigorous, orthogonal experimental validation.

## Supporting information

Supplementary manuscript

## 7. Data and Code Availability

The scRNA-seq data supporting the findings of this study are publicly available from the Tabula Muris Senis consortium (FACS-sorted data). All custom Python code, environment specifications, trained model files, preprocessed data files, and analysis pipelines used to generate the GeroEngine models and produce the figures in this manuscript have been deposited in a public GitHub repository https://github.com/ymb943/research_geroengine and are freely available under an MIT open-source license.

## Abbreviations

ALS: Algebraic Linear Shift
AP-1: Activator Protein 1
BH-FDR: Benjamini–Hochberg false discovery rate correction
E2F: E2 factor transcription factor family
FACS: Fluorescence-activated cell sorting
FDR: False discovery rate
GO: Gene Ontology
GRN: Gene regulatory network
GSEA: Gene Set Enrichment Analysis
HSC: Hematopoietic stem cell
HVG: Highly variable gene
JAK–STAT: Janus kinase–signal transducer and activator of transcription
KEGG: Kyoto Encyclopedia of Genes and Genomes
KL: Kullback–Leibler divergence
log_2_ FC: Logarithm base 2 fold change
LOPO: Leave-One-Pair-Out
MAD: Median absolute deviation
MCM: Minichromosome maintenance
MGI: Mouse Genome Informatics
MHC: Major histocompatibility complex
MHC-I: Major histocompatibility complex class I
MHC-II: Major histocompatibility complex class II
MMD: Maximum Mean Discrepancy
MSE: Mean squared error
NF-*κ*B: Nuclear factor-*κ*B
ODE: Ordinary differential equation
OOD: Out-of-distribution
OR: Odds ratio
PCA: Principal Component Analysis
PCC: Pearson correlation coefficient
PPR: Personalized PageRank
RBF: Radial basis function
SA: Supplementary Algorithm
scATAC-seq: Single-cell assay for transposase-accessible chromatin using sequencing
scRNA-seq: Single-cell RNA sequencing
SD: Standard deviation
SDi: Supplementary Discussion
SEq: Supplementary Equation
SF: Supplementary Figure
SM: Supplementary Methods
SR: Supplementary Results
ST: Supplementary Table
VAE: Variational Autoencoder

## 10.Terms

### GeroEngine-Specific Terms

**GeroEngine**: Computational framework for technical-artifact-aware generative simulation of single-cell aging trajectories and downstream regulator inference

**GeroSimulator**: VAE-based module that applies the learned latent aging direction to source cells and decodes simulated aged or reverse-simulated young profiles

**GeroVector**: Fold-specific population-level latent aging direction, denoted **g** or **g**_*fold*_, learned from mouse-balanced old-minus-young centroids in the VAE latent space

**GeroSimOld**: Simulated aged expression profile generated by applying +**g** to held-out young cells and decoding the shifted latent state

**GeroSimYoung**: Reverse-simulated young expression profile generated by applying *−***g** to held-out old cells and decoding the shifted latent state

**GeroTarget**: Consensus aging-associated target gene recovered across LOPO folds from simulated transcriptomic shifts

**GeroRegulator**: Putative upstream regulator inferred from consensus GeroTargets using reverse-directed network analysis

**GeroNetwork**: Tissue-specific regulator–target network integrating GeroTargets and GeroRegulators

### Mathematical Notation

#### Constants and Hyperparameters

*N*_*hvg*_: Constant. Number of highly variable genes retained during preprocessing

*N*_*cells*_: Constant. Number of cells in a given matrix

*N*_*latentdim*_: Constant. Dimensionality of the VAE latent space and the matched PCA subspace

*β*_*KL*_: Constant. Scaling factor for the Kullback–Leibler divergence regularization term

*p*_*dropout*_: Constant. Dropout probability used for VAE regularization

*ϵ*_*bn*_: Constant. Epsilon value for batch normalization in the VAE model

*α*: Constant. Temporal scaling factor used for 18-month chronological interpolation, *α* = (18*−*3)*/*(24*−*3) *≈*0.714

*L*: Integer. Directed regulatory path length used during GeroRegulator filtering

*n*: Integer. Union size of forward and reverse consensus target sets when displayed above overlap bars

#### Expression Space (Vectors and Matrices)

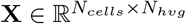: Matrix. Empirical preprocessed gene expression of cells

**X**_*young,test,fold*_, **X**_*old,test,fold*_: Matrices. Empirical held-out young and old expression matrices for a given fold

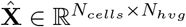 : Matrix. Predicted simulated gene expression of cells

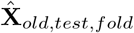: Matrix. GeroSimOld profile generated by forward young*→*old simulation

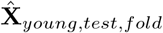: Matrix. GeroSimYoung profile generated by reverse old*→*young simulation

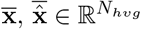 : Vectors. Empirical and predicted pseudo-bulked mean expression vectors

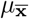: Scalar. Global mean of the target expression vector across the evaluated gene subset

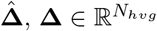: Vectors. Predicted and empirical mean transcriptomic shift vectors

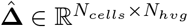: Matrix. Cell-wise predicted transcriptomic shift matrix

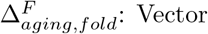: Vector. Forward aging-axis effect, defined as 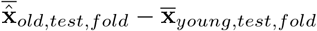

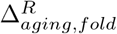: Vector. Reverse simulation effect re-expressed on the common young*→*old aging axis, defined as 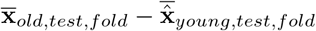

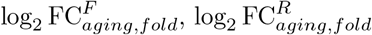: : Vectors. Forward and reverse-aligned log_2_ fold changes, computed by dividing 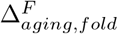 and 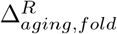 by ln 2

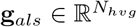: Vector. ALS aging trajectory vector in the original gene-expression space

#### Latent Space and Cosine Similarity

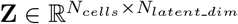 : Matrix. Encoded gene expression of cells in VAE latent or PCA space

**Z**_*young,test*_, **Z**_*young,test,fold*_: Matrices. Encoded latent states of held-out young cells

**Z**_*old,test*_, **Z**_*old,test,fold*_: Matrices. Encoded latent states of held-out old cells

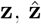: Vectors. Original and shifted latent representations of an individual cell

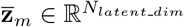 : Vector. Cell-level centroid, or mean latent vector, for mouse *m*

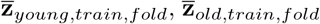: Vectors. Mouse-balanced young and old training centroids used to compute **g**_*fold*_

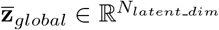 : Vector. Unweighted mouse-level global centroid in latent space

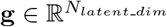 : Vector. Reference male GeroVector for a fold unless otherwise specified; computed as the mouse-balanced old-minus-young latent centroid difference in the primary male training cohort

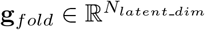 : Vector. Explicit fold-indexed GeroVector

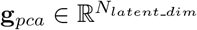 : Vector. Aging trajectory direction in the linear principal-component subspace

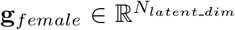: Vector. Female OOD aging direction for a fold; computed by encoding female OOD cells with that fold’s forward-trained VAE and taking the mouse-balanced old-minus-young latent centroid difference

*θ*: Scalar. Angle between two latent aging vectors

cos(*θ*): Scalar. Cosine similarity used in Results to summarize cross-sex vector alignment

cos(**g, g**_*female*_): Scalar. Fold-wise cross-sex cosine similarity, computed as ^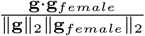^ and summarized across 12 LOPO folds as mean *±* SD with per-fold strip points

#### Evaluation Metrics (Variances and Networks)

*R*^2^: Scalar. Coefficient of determination measuring macroscopic alignment of mean expression profiles *R*_*shift*_: Scalar. Pearson correlation coefficient between predicted and empirical transcriptomic shift vectors

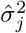: Scalar. Absolute variance of gene *j* across simulated cells

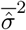: Scalar. Mean absolute gene variance across retained HVG features

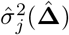: Scalar. Variance of predicted transcriptomic shifts for gene *j*

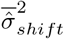: Scalar. Mean shift variance across retained HVG features

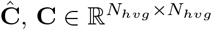 : Matrices. Predicted and empirical pairwise gene-gene Pearson correlation matrices

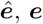: Vectors. Flattened network edge vectors containing strictly upper-triangular elements of Ĉ and **C**

*R*_*ggc*_: Scalar. Pearson correlation coefficient between flattened network edge vectors 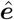 and ***e***

*r*: Scalar. Pearson correlation coefficient when used for continuous forward–reverse log_2_FC concordance

ΔJaccard: Scalar. Change in Jaccard overlap after clamp removal relative to the standard clamped pipeline

MMD: Scalar. Empirical Maximum Mean Discrepancy, computed using the biased V-statistic estimator

*k*(**u, v**): Function. Multi-scale Gaussian radial basis function kernel used for MMD

*σ*_*base*_: Scalar. Base bandwidth parameter for the RBF kernel, established using the median heuristic

*↑, ↓*: Symbols. Up-regulation and down-regulation on the common young*→*old aging axis

#### Topology, Algorithms, and Sets

En(*·*): Function. Encoder of the trained VAE De(*·*): Function. Decoder of the trained VAE

PCA(*·*): Function. Projection of expression data into the fitted PCA subspace

PCA^−1^(*·*): Function. Inverse projection from the PCA subspace back to the original gene-expression space

max(0, *·*): Function. Non-negativity clamp applied to decoded or inverse-transformed simulated expression values

*𝒟*: Dataset of scRNA-seq profiles with metadata such as age and mouse ID

*ℳ*: List. Stores the calculated mean latent vectors **z**_*m*_ for individual mouse subjects

## References

[1] Ailsa M. Jeffries, Tianxiong Yu, Jennifer S. Ziegenfuss, Allie K. Tolles, Christina E. Baer, Cesar Bautista Sotelo, Yerin Kim, Zhiping Weng, and Michael A. Lodato. Single-cell transcriptomic and genomic changes in the ageing human brain. Nature, 646(8085):657–666, Oct 2025. ISSN 1476-4687. doi: 10.1038/s41586-025-09435-8. URL 10.1038/s41586-025-09435-8.

[2] Josh Bartz, Hannim Jung, Karen Wasiluk, Lei Zhang, and Xiao Dong. Progress in discovering transcriptional noise in aging. International Journal of Molecular Sciences, 24(4), 2023. ISSN 1422-0067. doi: 10.3390/ijms24043701. URL https://www.mdpi.com/1422-0067/24/4/3701.

[3] Jincheng Wang, Yuchen Sang, Shengxian Jin, Xuezheng Wang, Gajendra Kumar Azad, Mark A. McCormick, Brian K. Kennedy, Qing Li, Jianbin Wang, Xiannian Zhang, Yi Zhang, and Yanyi Huang. Single-cell rna-seq reveals early heterogeneity during aging in yeast. Aging Cell, 21(11): e13712, 2022. doi: 10.1111/acel.13712. URL https://onlinelibrary.wiley.com/doi/abs/10.1111/acel.13712.e13712ACE-21-0054.R1.

[4] Peiru Wu, Xuyu Zhao, Zixin Chen, Jingying Huang, Tengteng Dai, Jianxin Zhou, Luyao Xiao, Luonan Chen, Robert Chunhua Zhao, and Jiao Wang. Identifying tipping points during healthy brain aging through single-nucleus transcriptomic analysis. Advanced Science, 12(41):e05779, 2025. doi: 10.1002/advs.202505779. URL https://advanced.onlinelibrary.wiley.com/doi/abs/10.1002/advs.202505779.

[5] Yunjin Li, Qixia Wang, Yuan Xuan, Jian Zhao, Jin Li, Yuncai Tian, Geng Chen, and Fei Tan. Investigation of human aging at the single-cell level. Ageing Res. Rev., 101(102530):102530, November 2024.

[6] Tabula Muris Consortium. A single-cell transcriptomic atlas characterizes ageing tissues in the mouse. Nature, 583(7817):590–595, July 2020.

[7] Qinhuan Luo, Yongzhen Yu, and Tianying Wang. Denoising single-cell RNA-seq data with a deep learning-embedded statistical framework. BMC Bioinformatics, 26(1):282, November 2025.

[8] Peng Qiu. Embracing the dropouts in single-cell RNA-seq analysis. Nat. Commun., 11(1):1169, March 2020.

[9] Martin Enge, H. Efsun Arda, Marco Mignardi, John Beausang, Rita Bottino, Seung K. Kim, and Stephen R. Quake. Single-Cell Analysis of Human Pancreas Reveals Transcriptional Signatures of Aging and Somatic Mutation Patterns. Cell, 171(2):321–330.e14, October 2017. ISSN 0092-8674. doi: 10.1016/j.cell.2017.09.004. URL https://pmc.ncbi.nlm.nih.gov/articles/PMC6047899/.

[10] Diederik P Kingma and Max Welling. Auto-encoding variational bayes, 2022. URL https://arxiv.org/abs/1312.6114.

[11] Romain Lopez, Jeffrey Regier, Michael B Cole, Michael I Jordan, and Nir Yosef. Deep generative modeling for single-cell transcriptomics. Nature Methods, 15(12):1053–1058, December 2018.

[12] Gö kcen Eraslan, Lukas M Simon, Maria Mircea, Nikola S Mueller, and Fabian J Theis. Single-cell RNA-seq denoising using a deep count autoencoder. Nat. Commun., 10(1):390, January 2019.

[13] Karl Pearson. Liii. on lines and planes of closest fit to systems of points in space. The London, Edinburgh, and Dublin Philosophical Magazine and Journal of Science, 2(11):559–572, 1901. doi: 10.1080/14786440109462720. URL 10.1080/14786440109462720.

[14] H Hotelling. Analysis of a complex of statistical variables into principal components, 1933.

[15] Volker Bergen, Marius Lange, Stefan Peidli, F. Alexander Wolf, and Fabian J. Theis. Generalizing rna velocity to transient cell states through dynamical modeling. Nature Biotechnology, 38(12): 1408–1414, August 2020. ISSN 1546-1696. doi: 10.1038/s41587-020-0591-3. URL 10.1038/s41587-020-0591-3.

[16] Marius Lange, Volker Bergen, Michal Klein, Manu Setty, Bernhard Reuter, Mostafa Bakhti, Heiko Lickert, Meshal Ansari, Janine Schniering, Herbert B. Schiller, Dana Pe’Er, and Fabian J. Theis. CellRank for directed single-cell fate mapping. Nature Methods, 19:159–170, 2022. doi: 10.1038/s41592-021-01346-6. CRIS-Team Scopus Importer:2023-03-08.

[17] Dominik Klein, Giovanni Palla, Marius Lange, Michal Klein, Zoe Piran, Manuel Gander, Laetitia Meng-Papaxanthos, Michael Sterr, Lama Saber, Changying Jing, Aimée Bastidas-Ponce, Perla Cota, Marta Tarquis-Medina, Shrey Parikh, Ilan Gold, Heiko Lickert, Mostafa Bakhti, Mor Nitzan, Marco Cuturi, and Fabian J Theis. Mapping cells through time and space with moscot. Nature, 638(8052): 1065–1075, February 2025.

[18] Cole Boyle. Trajectory inference for aging time courses of single-cell data. PhD thesis, University of British Columbia, 2026. URL https://open.library.ubc.ca/collections/ubctheses/24/items/1.0451843.

[19] Mohammad Lotfollahi, F Alexander Wolf, and Fabian J Theis. scgen predicts single-cell perturbation responses. Nature Methods, 16(8):715–721, August 2019.

[20] Matthew L. Bochman and Anthony Schwacha. The mcm complex: Unwinding the mechanism of a replicative helicase. Microbiology and Molecular Biology Reviews, 73(4):652–683, 2009. doi: 10.1128/mmbr.00019-09. URL https://journals.asm.org/doi/abs/10.1128/mmbr.00019-09.

[21] Wikipedia. POLD1 — Wikipedia, the free encyclopedia. http://en.wikipedia.org/w/index.php?title=POLD1&oldid=1329681568, 2026. [Online; accessed 01-May-2026].

[22] Che-Feng Chang, Brittany A. Goods, Michael H. Askenase, Hannah E. Beatty, Artem Osherov, Jonathan H. DeLong, Matthew D. Hammond, Jordan Massey, Margaret Landreneau, J. Christopher Love, and Lauren H. Sansing. Divergent functions of tissue-resident and blood-derived macrophages in the hemorrhagic brain. Stroke, 52(5):1798–1808, 2021. doi: 10.1161/STROKEAHA.120.032196. URL https://www.ahajournals.org/doi/abs/10.1161/STROKEAHA.120.032196.

[23] Philipp Novoszel, Barbara Drobits, Martin Holcmann, Cristiano De Sa Fernandes, Roland Tschismarov, Sophia Derdak, Thomas Decker, Erwin F Wagner, and Maria Sibilia. The AP-1 transcription factors c-jun and JunB are essential for CD8α conventional dendritic cell identity. Cell Death & Differentiation, 28(8):2404–2420, August 2021.

[24] Nicos A Nicola and Jeffrey J Babon. Leukemia inhibitory factor (LIF). Cytokine Growth Factor Rev., 26(5):533–544, October 2015.

[25] Yutaka Takahashi, Michiko Takahashi, Nick Carpino, Shiann-Tarng Jou, Jyh-Rong Chao, Satoshi Tanaka, Yasufumi Shigeyoshi, Evan Parganas, and James N Ihle. Leukemia inhibitory factor regulates trophoblast giant cell differentiation via janus kinase 1-signal transducer and activator of transcription 3-suppressor of cytokine signaling 3 pathway. Mol. Endocrinol., 22(7):1673–1681, July 2008.

[26] Hetvi Gandhi, Remigiusz Worch, Kristina Kurgonaite, Martin Hintersteiner, Petra Schwille, Christian Bökel and Thomas Weidemann. Dynamics and interaction of interleukin-4 receptor subunits in living cells. Biophys. J., 107(11):2515–2527, December 2014.

[27] Siti Muhamad Nur Husna, Norasnieda Md Shukri, Sharifah Emilia Tuan Sharif, Hern Tze Tina Tan, Noor Suryani Mohd Ashari, and Kah Keng Wong. IL-4/IL-13 axis in allergic rhinitis: Elevated serum cytokines levels and inverse association with tight junction molecules expression. Front. Mol. Biosci., 9:819772, March 2022.

[28] Ryan Ferrao, Heidi J A Wallweber, Hoangdung Ho, Christine Tam, Yvonne Franke, John Quinn, and Patrick J Lupardus. The structural basis for class II cytokine receptor recognition by JAK1. Structure, 24(6):897–905, June 2016.

[29] Timothy R Hammond, Connor Dufort, Lasse Dissing-Olesen, Stefanie Giera, Adam Young, Alec Wysoker, Alec J Walker, Frederick Gergits, Michael Segel, James Nemesh, Samuel E Marsh, Arpiar Saunders, Evan Macosko, Florent Ginhoux, Jinmiao Chen, Robin J M Franklin, Xianhua Piao, Steven A McCarroll, and Beth Stevens. Single-Cell RNA sequencing of microglia throughout the mouse lifespan and in the injured brain reveals complex Cell-State changes. Immunity, 50(1): 253–271.e6, January 2019.

[30] Hadas Keren-Shaul, Amit Spinrad, Assaf Weiner, Orit Matcovitch-Natan, Raz Dvir-Szternfeld, Tyler K Ulland, Eyal David, Kuti Baruch, David Lara-Astaiso, Beata Toth, Shalev Itzkovitz, Marco Colonna, Michal Schwartz, and Ido Amit. A unique microglia type associated with restricting development of alzheimer’s disease. Cell, 169(7):1276–1290.e17, June 2017.

[31] Md Akkas Ali, Md Hasanul Banna Siam, Donald Vardaman, 3rd, Chase Bolding, J Nicholas Brazell, Ashleigh D Whatley, Christopher A Risley, Harrison Tidwell, Syed Nakib Hossain, Juhi Samal, Ashley S Harms, Mallikarjun Patil, and Daniel J Tyrrell. High-dimensional single-cell analysis reveals coordinated age-dependent neuroinflammatory microglia-t cell circuits in the brain. December 2025.

[32] Ayaka Sugai, Hinase Moridono, Merve Bilgic, Yukiko Gotoh, and Yusuke Kishi. Age-associated microglia populations identified from several single cell transcriptome data. bioRxiv, 2025. doi: 10.1101/2025.07.28.665226. URL https://www.biorxiv.org/content/early/2025/07/31/2025.07.28.665226.

[33] Ignazio Antignano, Yingxiao Liu, Nina Offermann, and Melania Capasso. Aging microglia. Cellular and Molecular Life Sciences, 80(5):126, April 2023.

[34] Christopher P Wild. Complementing the genome with an “exposome”: the outstanding challenge of environmental exposure measurement in molecular epidemiology. Cancer epidemiology, biomarkers & prevention, 14(8):1847–1850, 2005.

[35] Carlos Ló pez-Otín, Maria A Blasco, Linda Partridge, Manuel Serrano, and Guido Kroemer. The hallmarks of aging. Cell, 153(6):1194–1217, 2013.

[36] Lawrence Page, Sergey Brin, Rajeev Motwani, and Terry Winograd. The pagerank citation ranking : Bringing order to the web. In The Web Conference, 1999. URL https://api.semanticscholar.org/CorpusID:1508503.

[37] Aric A. Hagberg, Daniel A. Schult, and Pieter J. Swart. Exploring network structure, dynamics, and function using networkx. In Gäel Varoquaux, Travis Vaught, and Jarrod Millman, editors, Proceedings of the 7th Python in Science Conference, pages 11–15, Pasadena, CA USA, 2008.

[38] Dénes Türei, Tamás Korcsmáros, and Julio Saez-Rodriguez. OmniPath: guidelines and gateway for literature-curated signaling pathway resources. Nat. Methods, 13(12):966–967, November 2016.

[39] Luz Garcia-Alonso, Christian H. Holland, Mahmoud M. Ibrahim, Denes Turei, and Julio Saez-Rodriguez. Benchmark and integration of resources for the estimation of human transcription factor activities. Genome Research, 29(8):1363–1375, 2019. doi: 10.1101/gr.240663.118. URL http://genome.cshlp.org/content/genome/29/8/1363.

[40] Tabula Muris Consortium. A single-cell transcriptomic atlas characterizes ageing tissues in the mouse. Nature, 583(7817):590–595, July 2020.

[41] Adam Gayoso, Romain Lopez, Galen Xing, Pierre Boyeau, Valeh Valiollah Pour Amiri, Justin Hong, Katherine Wu, Michael Jayasuriya, Edouard Mehlman, Maxime Langevin, Yining Liu, Jules Samaran, Gabriel Misrachi, Achille Nazaret, Oscar Clivio, Chenling Xu, Tal Ashuach, Mariano Gabitto, Mohammad Lotfollahi, Valentine Svensson, Eduardo da Veiga Beltrame, Vitalii Kleshchevnikov, Carlos Talavera-Ló pez Lior Pachter, Fabian J Theis, Aaron Streets, Michael I Jordan, Jeffrey Regier, and Nir Yosef. A python library for probabilistic analysis of single-cell omics data. Nature Biotechnology, 40(2):163–166, February 2022.

[42] Adam Paszke, Sam Gross, Francisco Massa, Adam Lerer, James Bradbury, Gregory Chanan, Trevor Killeen, Zeming Lin, Natalia Gimelshein, Luca Antiga, Alban Desmaison, Andreas Kö pf Edward Yang, Zach DeVito, Martin Raison, Alykhan Tejani, Sasank Chilamkurthy, Benoit Steiner, Lu Fang, Junjie Bai, and Soumith Chintala. Pytorch: An imperative style, high-performance deep learning library, 2019. URL https://arxiv.org/abs/1912.01703.

[43] Jordan W Squair, Matthieu Gautier, Claudia Kathe, Mark A Anderson, Nicholas D James, Thomas H Hutson, Rémi Hudelle, Taha Qaiser, Kaya J E Matson, Quentin Barraud, Ariel J Levine, Gioele La Manno, Michael A Skinnider, and Grégoire Courtine. Confronting false discoveries in single-cell differential expression. Nat. Commun., 12(1):5692, September 2021.

[44] Yusuf Roohani, Kexin Huang, and Jure Leskovec. Predicting transcriptional outcomes of novel multigene perturbations with GEARS. Nature Biotechnology, 42(6):927–935, June 2024.

[45] David van Dijk, Roshan Sharma, Juozas Nainys, Kristina Yim, Pooja Kathail, Ambrose J Carr, Cassandra Burdziak, Kevin R Moon, Christine L Chaffer, Diwakar Pattabiraman, Brian Bierie, Linas Mazutis, Guy Wolf, Smita Krishnaswamy, and Dana Pe’er. Recovering gene interactions from single-cell data using data diffusion. Cell, 174(3):716–729.e27, July 2018.

[46] Wenpin Hou, Zhicheng Ji, Hongkai Ji, and Stephanie C Hicks. A systematic evaluation of single-cell RNA-sequencing imputation methods. Genome Biol., 21(1):218, August 2020.

[47] Arthur Gretton, Karsten M. Borgwardt, Malte J. Rasch, Bernhard Schölkopf, and Alexander Smola. A kernel two-sample test. Journal of Machine Learning Research, 13(25):723–773, 2012. URL http://jmlr.org/papers/v13/gretton12a.html.

[48] Mingsheng Long, Yue Cao, Jianmin Wang, and Michael Jordan. Learning transferable features with deep adaptation networks. In Francis Bach and David Blei, editors, Proceedings of the 32nd International Conference on Machine Learning, volume 37 of Proceedings of Machine Learning Research, pages 97–105, Lille, France, 07–09 Jul 2015. PMLR. URL https://proceedings.mlr.press/v37/long15.html.

[49] Zhuoqing Fang, Xinyuan Liu, and Gary Peltz. Gseapy: a comprehensive package for performing gene set enrichment analysis in python. Bioinformatics, 39(1):btac757. 01 2023. ISSN 1367-4811. doi: 10.1093/bioinformatics/btac757. URL 10.1093/bioinformatics/btac757.

[50] M Kanehisa and S Goto. KEGG: kyoto encyclopedia of genes and genomes. Nucleic Acids Res., 28 (1):27–30, January 2000.

[51] Carol J Bult, Judith A Blake, Cynthia L Smith, James A Kadin, Joel E Richardson, and Mouse Genome Database Group. Mouse genome database (MGD) 2019. Nucleic Acids Res., 47(D1): D801–D806, January 2019.

[52] Michael Ashburner, Catherine A Ball, Judith A Blake, David Botstein, Heather Butler, J Michael Cherry, Allan P Davis, Kara Dolinski, Selina S Dwight, Janan T Eppig, Midori A Harris, David P Hill, Laurie Issel-Tarver, Andrew Kasarskis, Suzanna Lewis, John C Matese, Joel E Richardson, Martin Ringwald, Gerald M Rubin, and Gavin Sherlock. Gene ontology: tool for the unification of biology. Nature Genetics, 25(1):25–29, May 2000.

[53] The Gene Ontology Consortium. The gene ontology knowledgebase in 2026. Nucleic Acids Research, 54(D1):D1779–D1792, 01 2026. ISSN 1362-4962. doi: 10.1093/nar/gkaf1292. URL 10.1093/nar/gkaf1292.

[54] Ville Satopaa, Jeannie Albrecht, David Irwin, and Barath Raghavan. Finding a “kneedle” in a haystack: Detecting knee points in system behavior. In Proceedings of the 2011 31st International Conference on Distributed Computing Systems Workshops, ICDCSW ‘11, page 166–171, USA, 2011. IEEE Computer Society. ISBN 9780769543864. doi: 10.1109/ICDCSW.2011.20. URL 10.1109/ICDCSW.2011.20.

[55] The UniProt Consortium. Uniprot: the universal protein knowledgebase in 2025. Nucleic Acids Research, 53(D1):D609–D617, 01 2025. ISSN 1362-4962. doi: 10.1093/nar/gkae1010. URL 10.1093/nar/gkae1010.

[56] Paul Shannon, Andrew Markiel, Owen Ozier, Nitin S Baliga, Jonathan T Wang, Daniel Ramage, Nada Amin, Benno Schwikowski, and Trey Ideker. Cytoscape: a software environment for integrated models of biomolecular interaction networks. Genome Res., 13(11):2498–2504, November 2003.

[57] Pauli Virtanen, Ralf Gommers, Travis E. Oliphant, Matt Haberland, Tyler Reddy, David Cournapeau, Evgeni Burovski, Pearu Peterson, Warren Weckesser, Jonathan Bright, Stéfan J. van der Walt, Matthew Brett, Joshua Wilson, K. Jarrod Millman, Nikolay Mayorov, Andrew R. J. Nelson, Eric Jones, Robert Kern, Eric Larson,J J Carey, İlhan Polat, Yu Feng, Eric W. Moore, Jake VanderPlas, Denis Laxalde, Josef Perktold, Robert Cimrman, Ian Henriksen, E. A. Quintero, Charles R. Harris, Anne M. Archibald, Antônio H. Ribeiro, Fabian Pedregosa, Paul van Mulbregt, and SciPy 1.0 Contributors. SciPy 1.0: Fundamental Algorithms for Scientific Computing in Python. Nature Methods, 17:261–272, 2020. doi: 10.1038/s41592-019-0686-2.

[58] Charles R. Harris, K. Jarrod Millman, Stéfan J. van der Walt, Ralf Gommers, Pauli Virtanen, David Cournapeau, Eric Wieser, Julian Taylor, Sebastian Berg, Nathaniel J. Smith, Robert Kern, Matti Picus, Stephan Hoyer, Marten H. van Kerkwijk, Matthew Brett, Allan Haldane, Jaime Fernández del Río, Mark Wiebe, Pearu Peterson, Pierre Gérard-Marchant, Kevin Sheppard, Tyler Reddy, Warren Weckesser, Hameer Abbasi, Christoph Gohlke, and Travis E. Oliphant. Array programming with NumPy. Nature, 585(7825):357–362, September 2020. doi: 10.1038/s41586-020-2649-2. URL 10.1038/s41586-020-2649-2.

[59] Florian Charlier, Marc Weber, Dariusz Izak, Emerson Harkin, Marcin Magnus, Joseph Lalli, Louison Fresnais, Matt Chan, Nikolay Markov, Oren Amsalem, Sebastian Proost, Agamemnon Krasoulis, getzze, and Stefan Repplinger. Statannotations, October 2022. URL 10.5281/zenodo.7213391.

[60] Yoav Benjamini and Yosef Hochberg. Controlling the false discovery rate: A practical and powerful approach to multiple testing. Journal of the Royal Statistical Society: Series B (Methodological), 57(1):289–300, 01 1995. ISSN 0035-9246. doi: 10.1111/j.2517-6161.1995.tb02031.x. URL 10.1111/j.2517-6161.1995.tb02031.x.

[61] Ricky T. Q. Chen, Yulia Rubanova, Jesse Bettencourt, and David Duvenaud. Neural ordinary differential equations, 2019. URL https://arxiv.org/abs/1806.07366.

[62] Le Cong, F. Ann Ran, David Cox, Shuailiang Lin, Robert Barretto, Naomi Habib, Patrick D. Hsu, Xuebing Wu, Wenyan Jiang, Luciano A. Marraffini, and Feng Zhang. Multiplex genome engineering using crispr/cas systems. Science, 339(6121):819–823, 2013. doi: 10.1126/science.1231143. URL https://www.science.org/doi/abs/10.1126/science.1231143.

[63] Martin Jinek, Krzysztof Chylinski, Ines Fonfara, Michael Hauer, Jennifer A. Doudna, and Emmanuelle Charpentier. A programmable dual-rna–guided dna endonuclease in adaptive bacterial immunity. Science, 337(6096):816–821, 2012. doi: 10.1126/science.1225829. URL https://www.science.org/doi/abs/10.1126/science.1225829.

