## Supplementary manuscript for "GeroEngine: Generative single-cell aging trajectories reveal a bidirectionally traversable identity core and direction-specific inflammatory remodeling"

### SM 1. Supplementary Methods

#### SM 1.1. Dataset Acquisition, Single-Cell Preprocessing, and Cross-Validation Design

Raw transcriptomic count matrices were acquired from the Tabula Muris Senis atlas.[1] To mitigate cross-platform technical batch effects, downstream analyses were strictly restricted to the FACS-sorted dataset. Rigorous quality control was enforced by discarding cells exhibiting total transcript counts or unique gene counts exceeding 5 Median Absolute Deviations (MAD) above the population median. To establish a rigorous, sex-controlled baseline, the primary training and testing cohort was exclusively restricted to 3-month-old (Young) and 24-month-old (Old) male mice. After quality control filtering, the primary male cohort comprised 6434 microglial cells and 2113 HSCs. For subsequent OOD generalization testing, we established two independent validation cohorts: an unseen age cohort (18-month-old males; 1946 microglia, 277 HSCs) and an unseen sex cohort (3-month and 18-month-old females; 4613 microglia, 887 HSCs).

To eliminate irrelevant background noise, olfactory (*Olfr*) and vomeronasal (*Vmn*) receptor genes were first removed from the raw count matrices. Because the 'seurat_v3' method requires unnormalized count data to accurately model mean-variance relationships, Highly Variable Genes (HVGs) were then identified directly on the raw primary male cohort using Scanpy's `highly_variable_genes()` function.[2, 3] Notably, rather than arbitrarily selecting a fixed number of top-ranked features, we enforced a strict standardized variance cutoff of 1.3 to objectively retain biologically informative genes, yielding 3182 microglial and 2908 HSC transcripts (SF 1). Following HVG identification, the full datasets were library-size normalized to 10,000 counts per cell and log-transformed ($\ln(x+1)$) using Scanpy's `normalize_total()` and `log1p()` functions, respectively. Finally, to prevent data leakage and preserve geometric consistency during latent projection, all normalized cohorts, including the OOD 18-month males and all females, were strictly subsetted to match the exact HVG feature space defined by the primary cohort (SF 2).

To ensure absolute computational reproducibility, all algorithms, initializations, and sampling processes were fixed with a global random seed of 42.

To evaluate model generalization and ensure robust biological discovery within the primary cohort, we employed a strict LOPO cross-validation scheme. For each of the 12 folds, data from one 24-month-old mouse and one 3-month-old mouse were held out entirely as a test set, while the remaining mice constituted the training set. This entire preprocessing, feature selection, and cross-validation pipeline was executed independently for microglial cells (Brain Myeloid tissue) and HSCs (Marrow tissue).

### SM 1.2. Variational Autoencoder (VAE) Architecture and Optimization

To model cellular aging patterns and simulate perturbations, we adopted the scGen framework, a VAE architecture specialized for single-cell perturbation analysis, implemented via the `scvi-tools` framework in PyTorch.[4–7] However, to adapt this architecture for the specific biological and technical constraints of longitudinal aging, the GeroEngine pipeline extends the base methodology by coupling it with strict LOPO consensus filtering and individual-agnostic trajectory weighting.

The network consists of an encoder and a decoder, each comprising two fully connected hidden layers with 800 units. The encoder maps the high-dimensional gene expression space to a 100-dimensional latent space ($N_{latent\_dim} = 100$), which was empirically determined to balance low dimensionality with retained prediction accuracy $R_{shift}$ (SF 3). The decoder reconstructs the original expression profile in the natural log domain ($\ln(x + 1)$). The decoder output utilized a linear layer (no activation function); during inference only, to enforce biological non-negativity, any negative reconstructed expression values were mathematically clamped to zero.

To prevent overfitting and ensure robust generalization across biological heterogeneity, we applied Batch Normalization ($\epsilon_{bn} = 0.001$, momentum=0.01), LeakyReLU activation (negative slope=0.01), and Dropout ($p_{dropout} = 0.2$) after each hidden layer. To systematically counteract confounding biases introduced by disproportionate cellular contributions across individual mice and age cohorts, mini-batches were stochastically assembled utilizing a weighted random sampling strategy with replacement.

Network parameters were optimized for 100 epochs using the Adam algorithm[8] with a baseline learning rate of $10^{-3}$ and a fixed batch size of 256 (SF 4 and 5). The generative model was trained by minimizing a composite objective function comprising a Mean Squared Error (MSE) reconstruction loss and a Kullback-Leibler (KL) divergence regularization term with the scaling factor $\beta_{KL}$.[5, 9]

To evaluate the stability of the GeroSimulator, we conducted a comprehensive hyperparameter ablation study across latent dimensionalities ($N_{latent\_dim} \in \{50, 100, 200\}$) and KL penalties ($\beta_{KL} \in \{10^{-5}, 10^{-4}, 10^{-3}, 10^{-2}, 10^{-1}\}$). The architecture demonstrated exceptional robustness, with the vast majority of configurations yielding highly predictive trajectory correlations ($R_{shift} > 0.70$) across unseen validation folds (SF 3). While higher KL penalties (e.g., $\beta_{KL} = 0.1$) yielded slightly higher absolute mathematical correlations by aggressively smoothing the latent manifold, such heavy regularization inherently forces the latent embeddings toward an isotropic prior, artificially crushing transcriptomic variance. To prevent this mathematical distortion and ensure the generated profiles reflect true biological heterogeneity, we conservatively selected a balanced architecture ($N_{latent\_dim} = 100, \beta_{KL} = 10^{-3}$) for all downstream GeroNetwork extractions. By deliberately avoiding over-regularization, the GeroSimulator's ability to simulate the observed transcriptomic contraction remains an emergent biological property of the learned manifold, rather than a forced consequence of hyperparameter tuning.

To embed the aging pattern into a latent dimensional vector, we calculated $\mathbf{g}$. During this calculation, to isolate the aging signature from inherent biological biases, we implemented a strict mouse-balanced averaging algorithm. By sequentially calculating the unweighted mean vector $\bar{\mathbf{z}} \in \mathbb{R}^{N_{latent\_dim}}$ at the individual mouse level within the latent space, we ensured that every mouse subject contributed equally to the systemic aging signature (SA 1).

Finally, the predicted transcriptomic profiles of aged cells (GeroSimOld) are calculated via:

$$\mathbf{Z}_{young,test,fold} = \text{En}(\mathbf{X}_{young,test,fold}) \tag{SEq 1}$$

$$\hat{\mathbf{X}}_{old,test,fold} = \max\left(0, \text{De}(\mathbf{Z}_{young,test,fold} + \mathbf{g})\right) \tag{SEq 2}$$

where $\mathbf{X}, \hat{\mathbf{X}} \in \mathbb{R}^{N_{cells} \times N_{hvg}}$, and $\mathbf{Z} \in \mathbb{R}^{N_{cells} \times N_{latent\_dim}}$. The trajectory vector $\mathbf{g}$ is broadcasted across the cell dimension to perform the latent addition.

**Algorithm 1** Mouse-Balanced Latent Space Averaging for GeroVector Extraction

**Input:**

- $\mathcal{D}_{train}$: Dataset of scRNA-seq profiles containing metadata for age and mouse ID.
- En(·): The encoder function of the trained VAE mapping preprocessed gene expression $\mathbf{X} \in \mathbb{R}^{N_{cells} \times N_{hvg}}$ to the latent space $\mathbf{Z} \in \mathbb{R}^{N_{cells} \times N_{latent\_dim}}$.

**Output:**

- $\mathbf{g}$: The individual-agnostic aging vector in the latent space.

**procedure** MouseBalancedMean($\mathcal{D}$)
  $\mathcal{M} \leftarrow [\,]$ ▷ Initialize list to store mean latent vectors ($\bar{\mathbf{z}}_m \in \mathbb{R}^{N_{latent\_dim}}$) for each mouse
  **for all** unique mouse $m \in \mathcal{D}$ **do**
    $\mathbf{X}_m \leftarrow \{x \in \mathcal{D} \mid \text{mouse\_id} = m\}$ ▷ Isolate all cells belonging to subject $m$
    $\mathbf{Z}_m \leftarrow \text{En}(\mathbf{X}_m)$ ▷ Project individual's cells into latent manifold
    $\bar{\mathbf{z}}_m \leftarrow \frac{1}{|\mathbf{Z}_m|} \sum_{\mathbf{z} \in \mathbf{Z}_m} \mathbf{z}$ ▷ Compute cell-level centroid for mouse $m$
    Append $\bar{\mathbf{z}}_m$ to $\mathcal{M}$
  $\bar{\mathbf{z}}_{global} \leftarrow \frac{1}{|\mathcal{M}|} \sum_{\bar{\mathbf{z}} \in \mathcal{M}} \bar{\mathbf{z}}$ ▷ Compute the unweighted mouse-level global centroid
  **return** $\bar{\mathbf{z}}_{global}$

**Execution:**
Partition $\mathcal{D}_{train,fold}$ into $\mathcal{D}_{young,train,fold}$ and $\mathcal{D}_{old,train,fold}$ based on temporal metadata
$\bar{\mathbf{z}}_{young,train,fold} \leftarrow$ MouseBalancedMean($\mathcal{D}_{young,train,fold}$)
$\bar{\mathbf{z}}_{old,train,fold} \leftarrow$ MouseBalancedMean($\mathcal{D}_{old,train,fold}$)
$\mathbf{g} \leftarrow \bar{\mathbf{z}}_{old,train,fold} - \bar{\mathbf{z}}_{young,train,fold}$ ▷ Calculate the aging trajectory

#### SM 1.3. Deterministic Baselines and In Silico Trajectory Generation

To rigorously evaluate the necessity of non-linear generative inference, we established two fundamental linear perturbation baselines: Algebraic Linear Shift (ALS) and PCA. To prevent data leakage across all models, aging trajectory vectors were calculated strictly within the training partitions of each cross-validation fold. Furthermore, to prevent trajectory bias driven by variable cellular recovery rates across individual subjects, all aging vectors were computed using a mouse-balanced mean. Specifically, the mean transcriptomic or latent state was first calculated independently for each mouse subject within the respective age group; these individual mouse centroids were subsequently averaged to establish the final, balanced group centroid.

For the ALS baseline, the aging trajectory vector was calculated directly in the $\ln(x+1)$-normalized expression space as

$$\mathbf{g}_{als} = \bar{\mathbf{x}}_{old,train,fold} - \bar{\mathbf{x}}_{young,train,fold} \tag{SEq 3}$$

where $\mathbf{g}_{als} \in \mathbb{R}^{N_{hvg}}$. Simulated aged cells were generated via direct vector addition:

$$\hat{\mathbf{X}}_{old,test,fold} = \mathbf{X}_{young,test,fold} + \mathbf{g}_{als}. \tag{SEq 4}$$

To maintain biological validity, any negative expression values in the ALS predictions were subsequently clamped to zero.

For the PCA baseline, the principal component model was fitted exclusively on the training set. The training and test count matrices were then projected into this linear subspace, retaining the first 100 principal components to mathematically align with the VAE latent dimensionality. The trajectory vector was computed utilizing the identical mouse-balanced centroid logic within this lower-dimensional space

$$\mathbf{g}_{pca} = \bar{\mathbf{z}}_{old,train,fold} - \bar{\mathbf{z}}_{young,train,fold} \tag{SEq 5}$$

where $\mathbf{g}_{pca} \in \mathbb{R}^{N_{latent\_dim}}$, applied to the young test cell embeddings, and inverse-transformed back to the original gene space:

$$\mathbf{Z}_{young,test,fold} = \text{PCA}(\mathbf{X}_{young,test,fold}) \tag{SEq 6}$$

$$\hat{\mathbf{X}}_{old,test,fold} = \max\left(0, \text{PCA}^{-1}(\mathbf{Z}_{young,test,fold} + \mathbf{g}_{pca})\right) \tag{SEq 7}$$

where $\mathbf{X}, \hat{\mathbf{X}} \in \mathbb{R}^{N_{cells} \times N_{hvg}}$, and $\mathbf{Z} \in \mathbb{R}^{N_{cells} \times N_{latent\_dim}}$. The trajectory vector $\mathbf{g}_{pca}$ is broadcasted across the cell dimension to perform the latent addition.

#### SM 1.4. Reverse-Direction (Old→Young) Trajectory Simulation

To evaluate whether the learned aging trajectory behaves as a coherent bidirectional axis, we performed reverse-direction simulation using the same fold-specific trained models and the same fold-specific training-derived aging vectors used in the forward young→old analysis. No model was re-trained for reverse simulation, and no held-out young or old test cells were used to recalculate trajectory vectors. Within each LOPO fold, the VAE aging vector was defined exclusively from the training cohort:

$$\mathbf{g}_{fold} = \overline{\mathbf{z}}_{old,train,fold} - \overline{\mathbf{z}}_{young,train,fold}. \tag{SEq 8}$$

Reverse VAE predictions were generated by encoding held-out old cells and applying the negated trajectory vector in latent space prior to decoding:

$$\mathbf{Z}_{old,test,fold} = \mathrm{En}_{fold}(\mathbf{X}_{old,test,fold}) \tag{SEq 9}$$

$$\hat{\mathbf{X}}^{VAE}_{young,test,fold} = \max(0,\ \mathrm{De}_{fold}(\mathbf{Z}_{old,test,fold} - \mathbf{g}_{fold}))\,, \tag{SEq 10}$$

yielding reverse-simulated young profiles, termed GeroSimYoung.

For PCA, held-out old cells were projected into the same fold-specific PCA space learned during the forward training fold, shifted by the negated PCA aging vector, and inverse-transformed back to gene-expression space:

$$\hat{\mathbf{X}}^{PCA}_{young,test,fold} = \max\Big(0,\ \mathrm{PCA}^{-1}_{fold}(\mathrm{PCA}_{fold}(\mathbf{X}_{old,test,fold}) - \mathbf{g}_{pca,fold})\Big)\,. \tag{SEq 11}$$

For ALS, the same training-derived expression-space vector was subtracted directly from held-out old cells:

$$\hat{\mathbf{X}}^{ALS}_{young,test,fold} = \max(0,\ \mathbf{X}_{old,test,fold} - \mathbf{g}_{als,fold})\,. \tag{SEq 12}$$

Because reverse simulation explicitly applies the negated forward vector, population-level movement toward the young centroid and mean-level sign reversal are expected by construction. We therefore interpreted reverse simulation using distributional metrics, continuous gene-effect concordance, and forward–reverse consensus-target overlap structure rather than centroid movement alone. All evaluation metrics, the 12-fold LOPO scheme, and the consensus GeroTarget extraction procedure were applied identically to the reverse direction, benchmarking GeroSimYoung against the held-out young reference cells.

#### SM 1.5. Aging-axis alignment for forward–reverse GeroTarget analysis

Although reverse simulation generates GeroSimYoung from real-old cells, all target-level signs were expressed on a common young→old aging axis. For forward simulation, the aging-axis effect was defined as:

$$\Delta^{F}_{aging,fold} = \overline{\hat{\mathbf{x}}}_{old,test,fold} - \overline{\mathbf{x}}_{young,test,fold}, \tag{SEq 13}$$

where $\overline{\hat{\mathbf{x}}}_{old,test,fold}$ denotes the pseudo-bulked GeroSimOld profile and $\overline{\mathbf{x}}_{young,test,fold}$ denotes the paired real-young source profile.

For reverse simulation, the old→young generated state was re-expressed on the same young→old axis:

$$\Delta^{R}_{aging,fold} = \overline{\mathbf{x}}_{old,test,fold} - \overline{\hat{\mathbf{x}}}_{young,test,fold}, \tag{SEq 14}$$

where $\overline{\mathbf{x}}_{old,test,fold}$ denotes the real-old source profile and $\overline{\hat{\mathbf{x}}}_{young,test,fold}$ denotes the pseudo-bulked GeroSimYoung profile.

The corresponding fold-specific log-fold changes were:

$$\log_2 \mathrm{FC}^{F}_{aging,fold} = \frac{\Delta^{F}_{aging,fold}}{\ln 2}, \qquad \log_2 \mathrm{FC}^{R}_{aging,fold} = \frac{\Delta^{R}_{aging,fold}}{\ln 2}. \tag{SEq 15}$$

Under this convention, positive values indicate genes higher on the aged side of the axis, and negative values indicate genes lower on the aged side of the axis. Bidirectionally shared GeroTargets were therefore considered sign-consistent when $\log_2 \mathrm{FC}^{F}_{aging,fold}$ and $\log_2 \mathrm{FC}^{R}_{aging,fold}$ had the same direction after 12-fold consensus filtering.

#### SM 1.6. Out-of-Distribution (OOD) Validation on Unseen Age and Sex

To assess the generalizability of the generated trajectories, the models were subjected to rigorous OOD validation on unseen intermediate ages (18-month) and an unseen sex (female mice). To simulate intermediate chronological aging, the 24-month aging vector was linearly interpolated using a temporal scaling factor of $\alpha = \frac{18-3}{24-3} \approx 0.714$.

For all OOD analyses, no PCA basis or trajectory vector was estimated from OOD evaluation cells. The VAE encoder/decoder weights and GeroVector $\mathbf{g}$, the PCA basis and PCA aging vector $\mathbf{g}_{pca}$, and the ALS expression-space aging vector $\mathbf{g}_{als}$ were all derived exclusively from the corresponding LOPO training cohort of the primary male forward analysis. OOD cells (18-month males and all females) were used only as inputs transformed through these training-derived mappings prior to vector application—scaled by $\alpha$ for the 18-month interpolation, or applied directly for cross-sex evaluation—and reconstruction. This ensures that the OOD comparison between VAE and linear baselines reflects only the action of each model's pre-trained components on unseen biological inputs, with no fold-internal re-estimation that could differentially advantage any model.

Directional alignment between the primary male GeroVector and each evaluation condition was quantified fold-wise by encoding both the fixed primary training cohort and the corresponding held-out or OOD cohort through the same primary male forward-trained VAE. Cosine similarity was computed between the training-derived vector $\mathbf{g}_{fold}^{train} = \overline{\mathbf{z}}_{old}^{train} - \overline{\mathbf{z}}_{young}^{train}$ and the evaluation-derived vector $\mathbf{g}_{fold}^{eval} = \overline{\mathbf{z}}_{target}^{eval} - \overline{\mathbf{z}}_{source}^{eval}$ as $\cos(\theta_{fold}) = \frac{\mathbf{g}_{fold}^{train} \cdot \mathbf{g}_{fold}^{eval}}{\|\mathbf{g}_{fold}^{train}\| \|\mathbf{g}_{fold}^{eval}\|}$; fold-level similarities were summarized as bars with individual folds overlaid as strip dots.

#### SM 1.7. Quantitative Evaluation Metrics (Full Definitions)

To systematically assess the fidelity of the simulated longitudinal trajectories, we employed a comprehensive suite of quantitative metrics evaluating macroscopic alignment, directional accuracy, technical-artifact compression, preservation of trajectory heterogeneity, and global manifold topology. A consolidated visual and algebraic summary of all evaluation metrics is provided in SF 7.

**Coefficient of Determination ($R^2$) of Mean Expression.** We utilized the $R^2$ score on pseudo-bulked expression profiles to measure macroscopic alignment, a highly robust approach for evaluating population-level transcriptomic shifts.[10] This metric evaluates the deterministic accuracy of the model in predicting the average expression levels of the aged population.[4] Let $\overline{\mathbf{x}}$ be the pseudo-bulked mean expression vector derived from the empirical held-out aged-cell reference state ($\mathbf{X}_{old,test,fold}$), and let $\hat{\overline{\mathbf{x}}}$ be the predicted mean expression vector derived from the simulated age state ($\hat{\mathbf{X}}_{old,test,fold}$). The score is calculated across all HVGs retained during preprocessing:

$$R^2 = 1 - \frac{\sum_{i=1}^{N_{hvg}} (\overline{x}_i - \hat{\overline{x}}_i)^2}{\sum_{i=1}^{N_{hvg}} (\overline{x}_i - \mu_{\overline{\mathbf{x}}})^2} \quad \text{(SEq 16)}$$

where $N_{hvg}$ is the total number of HVGs, and $\mu_{\overline{\mathbf{x}}}$ is the global mean of the target expression vector across this subset. A higher $R^2$ score indicates superior deterministic accuracy in capturing the active transcriptomic profile of the final aged state.

**Pearson Correlation ($R$) of Transcriptomic Shifts.** To verify that the generative model captured the true aging trajectory rather than merely reproducing a static cellular identity, we evaluated the Pearson correlation ($R$) between the predicted and empirical transcriptomic shifts.[11] Let $\overline{\mathbf{x}}_{young}$ and $\overline{\mathbf{x}}_{old}$ be the pseudo-bulked mean expression vectors derived from the empirical baseline ($\mathbf{X}_{young,test,fold}$) and target ($\mathbf{X}_{old,test,fold}$) states, respectively. Similarly, let $\hat{\overline{\mathbf{x}}}_{old}$ be the predicted mean expression vector derived from the simulated aged state ($\hat{\mathbf{X}}_{old,test,fold}$). The shift vectors were isolated by subtracting the young baseline centroid from both the predicted and empirical aged states:

$$\hat{\mathbf{\Delta}} = \hat{\overline{\mathbf{x}}}_{old} - \overline{\mathbf{x}}_{young} \quad \text{(SEq 17)}$$

$$\mathbf{\Delta} = \overline{\mathbf{x}}_{old} - \overline{\mathbf{x}}_{young} \quad \text{(SEq 18)}$$

where $\mathbf{\Delta} \in \mathbb{R}^{N_{hvg}}$, calculated strictly across the HVGs retained during preprocessing. This subtraction effectively eliminates invariant cell-type markers, isolating the dynamic transcriptomic signature of cellular senescence. A higher correlation coefficient ($R_{shift} = \mathrm{PCC}(\hat{\mathbf{\Delta}}, \mathbf{\Delta})$) indicates that the model successfully learned the true biological directionality of aging, correctly upregulating and downregulating the appropriate regulatory networks.

**Mean of Gene Variances.** To evaluate the extent of technical artifact compression, while separately monitoring biological heterogeneity, we quantified the mean of gene variances across the simulated cell populations. For a given predicted aged expression matrix $\hat{\mathbf{X}}_{old,test,fold}$ (comprising $N_{cells}$), the absolute variance of each HVG $j$ across all simulated cells ($\hat{\sigma}_j^2$) was calculated. The mean absolute gene variance was then computed strictly across the $N_{hvg}$ retained features as:

$$\bar{\sigma}^2 = \frac{1}{N_{hvg}} \sum_{j=1}^{N_{hvg}} \hat{\sigma}_j^2 \qquad \text{(SEq 19)}$$

Because raw single-cell data is inherently zero-inflated, it exhibits an artificially inflated absolute variance.[12] A generative model that compresses technical structural noise through its latent bottleneck would be expected to yield lower mean gene variance, while a model that simply propagates source-cell artifacts would not.[13] Therefore, lower mean gene variance is interpreted only as evidence of reduced technical artifact carryover when accompanied by preserved shift variance and manifold fidelity; it should not be taken to mean that age-associated biological variability has been removed or that a deterministic ground-truth manifold has been isolated. Rather, this metric is intended to detect propagation of technical source noise, a known failure mode of shift-invariant linear projections.

**Mean of Gene Shift Variances.** To detect algorithmic mode collapse and evaluate trajectory heterogeneity, we calculated the mean of gene shift variances. First, a cell-wise predicted transcriptomic shift matrix ($\hat{\mathbf{\Delta}}$) was established by subtracting the empirical young source matrix from the predicted aged matrix:

$$\hat{\mathbf{\Delta}} = \hat{\mathbf{X}}_{old,test,fold} - \mathbf{X}_{young,test,fold} \qquad \text{(SEq 20)}$$

where $\hat{\mathbf{\Delta}} \in \mathbb{R}^{N_{cells} \times N_{hvg}}$. We then calculated the variance of these transcriptomic shifts for each individual HVG $j$ across all simulated cells, denoted as $\hat{\sigma}_j^2(\hat{\mathbf{\Delta}})$. The mean shift variance was subsequently computed strictly across the retained features as:

$$\bar{\hat{\sigma}}^2_{shift} = \frac{1}{N_{hvg}} \sum_{j=1}^{N_{hvg}} \hat{\sigma}_j^2(\hat{\mathbf{\Delta}}) \qquad \text{(SEq 21)}$$

A shift variance approaching zero indicates severe mode collapse, wherein a model forces all cells along an artificially rigid, uniform trajectory (such as a standard shift-invariant linear projection). Conversely, a higher mean shift variance indicates a robust generative model that successfully avoids collapse, predicting complex, non-linear trajectories that preserve the natural cell-to-cell stochasticity inherent to biological aging.

**Gene-Gene Correlation Matrix for Technical-Artifact-Aware Network Reconstruction.** To evaluate the fidelity of the reconstructed Gene Regulatory Networks (GRNs) and quantify whether simulated profiles preserve multivariate gene-gene relationships while reducing technical artifact carryover, we generated pairwise gene-gene co-expression matrices, a well-established benchmarking approach for single-cell data recovery.[14, 15] For both the predicted aged state ($\hat{\mathbf{X}}_{old,test,fold}$) and the empirical held-out aged-cell reference state ($\mathbf{X}_{old,test,fold}$), we computed their respective Pearson correlation matrices, $\hat{\mathbf{C}}$ and $\mathbf{C} \in \mathbb{R}^{N_{hvg} \times N_{hvg}}$. Each element $\mathbf{C}_{i,j}$ represents the Pearson correlation coefficient between the log-normalized expression of gene $i$ and gene $j$ across the cellular population. To isolate unique regulatory edges and eliminate redundant self-correlations (the matrix diagonal), we extracted the strictly upper-triangular elements ($i < j$) of both matrices into flattened, one-dimensional network edge vectors:

$$\hat{\boldsymbol{e}}_{old,test,fold} = \{\hat{\mathbf{C}}_{i,j} \mid 1 \leq i < j \leq N_{hvg}\} \qquad \text{(SEq 22)}$$

$$\boldsymbol{e}_{old,test,fold} = \{\mathbf{C}_{i,j} \mid 1 \leq i < j \leq N_{hvg}\} \quad \text{(SEq 23)}$$

The global structural concordance of the simulated and real aged networks was then definitively quantified by calculating the Pearson correlation coefficient between these flattened edge vectors: $R_{ggc} = \text{PCC}(\hat{\boldsymbol{e}}_{old,test,fold}, \boldsymbol{e}_{old,test,fold})$. A higher $R_{ggc}$ score indicates that the simulated population faithfully preserves the complex, multivariate regulatory dependencies of true aged tissue rather than predicting biologically disjointed gene expressions. Crucially, unlike linear models that achieve baseline correlation scores by artificially propagating high-variance technical dropouts, a higher $R_{ggc}$ score for the generative model suggests better preservation of coordinated regulatory structure after reducing dropout-associated artifacts, rather than definitive recovery of a dropout-free biological ground truth, consistent with the denoising capabilities of non-linear autoencoders.[13]

**Maximum Mean Discrepancy (MMD) for Global Manifold Alignment.** Maximum Mean Discrepancy (MMD) evaluates the distance between two probability distributions by mapping them into a Reproducing Kernel Hilbert Space (RKHS).[16] To robustly capture manifold topology across both local and global scales, we utilized a multi-scale Gaussian Radial Basis Function (RBF) kernel.[17] For a predicted aged expression matrix $\hat{\mathbf{X}}_{old,test,fold}$ (comprising $m$ cells) and the empirical held-out aged-cell reference matrix $\mathbf{X}_{old,test,fold}$ (comprising $n$ cells), let $\hat{\mathbf{x}}_i$ and $\mathbf{x}_j$ represent their respective individual cellular transcriptomic vectors. The base bandwidth parameter ($\sigma_{base}$) was established using the median heuristic based on pairwise Euclidean distances between the two distributions [16]:

$$2\sigma_{base}^2 = \text{median}(||\hat{\mathbf{x}}_i - \mathbf{x}_j||^2) \quad \text{(SEq 24)}$$

To evaluate topological alignment simultaneously at fine, intermediate, and coarse resolutions, the final kernel $k(\mathbf{u}, \mathbf{v})$ was defined as an unweighted mixture of three RBF kernels with scaled bandwidths $S = \{0.5\sigma_{base}, \sigma_{base}, 2.0\sigma_{base}\}$:

$$k(\mathbf{u}, \mathbf{v}) = \sum_{s \in S} \exp\left(-\frac{||\mathbf{u} - \mathbf{v}||^2}{2s^2}\right) \quad \text{(SEq 25)}$$

The empirical MMD was computed using the biased V-statistic estimator,[16] which is asymptotically consistent and widely used in deep generative modeling for its computational simplicity and negligible bias at large sample sizes:

$$\text{MMD}(\hat{\mathbf{X}}_{old,test,fold}, \mathbf{X}_{old,test,fold}) = \frac{1}{m^2}\sum_{i=1}^{m}\sum_{j=1}^{m} k(\hat{\mathbf{x}}_i, \hat{\mathbf{x}}_j) + \frac{1}{n^2}\sum_{i=1}^{n}\sum_{j=1}^{n} k(\mathbf{x}_i, \mathbf{x}_j) - \frac{2}{mn}\sum_{i=1}^{m}\sum_{j=1}^{n} k(\hat{\mathbf{x}}_i, \mathbf{x}_j) \quad \text{(SEq 26)}$$

This quantitative manifold distance was evaluated against two reference distributions: the Raw Target, representing empirical *in vivo* old cells containing both biological heterogeneity and scRNA-seq technical artifacts, and the VAE-Reconstructed Target, representing empirical aged cells passed through the same VAE reconstruction procedure to reduce dropout- and zero-inflation-associated technical artifacts. A lower MMD score indicates closer alignment to the corresponding reference distribution, rather than definitive recovery of an absolute biological ground truth.

**Holistic Interpretation and the Confounding Effect of Technical Noise.** It is critical to evaluate these quantitative metrics holistically rather than in isolation due to the severe confounding effects of single-cell technical noise. Because raw transcriptomic data is highly zero-inflated, deterministic linear projections (ALS and PCA) inherently propagate this technical artifact and sparsity structure directly from the source cells into the simulated predictions. Consequently, linear baselines can achieve artificially high scores in deterministic bulk alignment ($R^2$) and global network correlation ($R_{ggc}$), not because they have successfully inferred dynamic aging biology, but simply because they perfectly preserved the static technical artifacts of the starting population. To definitively distinguish true biological trajectory inference from artifact propagation, a model cannot simply maximize $R^2$; it must demonstrate the dual capacity to maintain accurate dynamic directionality (high $R_{shift}$ and $\bar{\sigma}^2_{shift}$) while reducing technical artifact carryover, preserving non-collapsed trajectory heterogeneity, and converging upon a technical-artifact-reduced topological reference (low VAE-reconstructed-reference MMD).

#### SM 1.8. Consensus Aging Genes and Pathway Enrichment Analysis

To overcome the magnitude bias inherent in traditional cross-sectional transcriptomics, we implemented a rank-based cross-validation consensus approach. Because the underlying single-cell expression matrices were preprocessed in the natural log domain ($\ln(x+1)$), the empirical young and predicted old expressions were first pseudo-bulked by their unweighted means. The longitudinal transcriptomic shift was then calculated and mathematically converted to the standard $\log_2$ Fold Change ($\log_2$ FC) scale by dividing the mean difference by $\ln(2)$.

Recognizing the biological asymmetry of aging, we bypassed arbitrary static magnitude thresholds (e.g., $|\log_2 \mathrm{FC}| > 1.0$) and segregated the transcriptomic shifts by absolute directionality. Crucially, a strict baseline threshold of 0 was enforced prior to ranking. This guarantees that only genes exhibiting a genuine biological increase ($\log_2 \mathrm{FC} > 0$) or decrease ($\log_2 \mathrm{FC} < 0$) are eligible for selection, preventing the algorithmic misclassification of low-magnitude, opposite-shift genes during fixed-size subset extraction. Within these validated directional pools, we ranked the HVG space to extract the top 100 up-regulated and top 100 down-regulated genes per fold. Only genes that ranked within this top 100 signature across all 12 independent hold-out folds (100% consensus requirement) were finalized as definitive GeroTargets.

Downstream GSEA was performed using the Enrichr API (gseapy) querying the KEGG_2019_Mouse and GO_Biological_Process_2021 libraries.[18–22] Statistical significance was defined using an adjusted p-value cutoff of $< 0.05$ (BH-FDR). Critically, for all enrichment tests, the complete set of tissue-specific HVGs was utilized as the statistical background to prevent artificial inflation of significance.

#### SM 1.9. Master Regulator Inference and Aging Network Construction

To isolate true GeroRegulators from background topological noise, we implemented a sequential, multi-stage algorithmic filtering pipeline grounded in literature-curated interaction networks (OmniPath and DoRothEA).[23, 24] First, a reverse-directed PPR algorithm was executed on these directed graphs.[25, 26] By utilizing the consensus GeroTargets as seed nodes (personalization = 1.0) and reversing the directionality of the network edges, the algorithm mathematically prioritized upstream structural hubs based on their path connectivity to the downstream aging signature. Crucially, Topological GeroRegulators were not defined by static network topology alone; they were subsequently filtered through a strict biological intersection. A candidate network hub was retained only if it successfully intersected with the active pool of potential aging genes, ensuring the regulator was both structurally central to the network and dynamically involved in the simulated aging trajectory.

Following this intersection, to prevent unmapped computational proteins from skewing the mathematical distribution of the topological scores, we utilized regular expressions (e.g., `^Gm`, `^AOA`) to discard initial computational artifacts. We then applied the Kneedle algorithm to the descending topological scores.[27] By calculating the maximum perpendicular distance from the score curve to a linear secant line, this algorithm objectively identifies the mathematical elbow point, allowing us to drop all candidate genes below this dynamic noise threshold (SF 6).

Subsequently, an advanced structural regular expression (`[A-Z0-9]{6,}`) was applied to purge any remaining raw UniProt accessions, restricting the candidate pool strictly to structurally validated, official MGI mouse gene symbols.[28] We then enforced a strict structural path constraint: candidate regulators were retained only if they possessed a directed regulatory path of length $L \leq 2$ (representing direct or single-intermediary regulation) to the core consensus GeroTargets.

Finally, to prevent "hub bias" while adapting to the unique topological sparsity of each cell type, we applied network-size adaptive thresholds. For the dense HSC regulatory network, we enforced strict constraints, requiring a minimum out-degree of $\geq 4$ distinct GeroTargets controlled alongside an absolute PageRank score $> 0.01$. However, applying these identical, stringent constraints to the Microglia network resulted in severe over-filtering, yielding an artificially small regulatory network. Consequently, we relaxed the thresholds for Microglia to an out-degree of $\geq 3$ and a PageRank score $> 0.005$ to preserve its biologically meaningful regulatory architecture. The resulting GeroNetworks were subsequently visualized using Cytoscape.[29]

#### SM 1.10. Statistical Analysis

All statistical analyses and quantitative metric evaluations were performed using Python (SciPy, NumPy, and statannotations libraries).[30–32] Because generative and linear model evaluations were strictly paired across the 12 independent LOPO cross-validation folds, statistical significance between comparative models (e.g., VAE vs. PCA, VAE vs. ALS) was determined using a paired Wilcoxon signed-rank test. To strictly control for multiple hypothesis testing across concurrent model comparisons, all derived p-values were subsequently adjusted utilizing the Benjamini-Hochberg False Discovery Rate (BH-FDR) correction.[33] For all quantitative benchmarks, data are presented as the mean $\pm$ standard deviation across the 12 folds. Statistical significance thresholds displayed in all comparative figures reflect these BH-FDR adjusted p-values and were defined *a priori* as * $p < 0.05$, ** $p < 0.01$, *** $p < 0.001$, and **** $p < 0.0001$. For downstream GSEA, statistical significance was similarly determined utilizing a BH-FDR adjusted p-value threshold of $< 0.05$.

### SR 2. Supplementary Results

#### SR 2.1. Detailed Pathway Enrichment Comparison in Aging Microglia

GSEA of the consensus aging transcripts revealed a profound divergence in how linear versus non-linear models capture age-related transcriptomic shifts in microglia. The VAE-derived signatures exhibited significantly higher sensitivity for lineage-specific, upstream functional terms. While the VAE also captured the age-associated MHC response, it additionally prioritized myeloid-centric pathways such as Osteoclast differentiation (OR = 14.9) and Leishmaniasis (OR = 16.8), suggesting that the model did not reduce the aging signature to a single high-variance inflammatory axis (ST 1).

Crucially, gene-level analysis demonstrated that the VAE achieved this specificity by comprehensively capturing upstream regulatory networks rather than merely identifying downstream structural effectors. For example, while linear models successfully isolated partial downstream markers, they completely failed to enrich the 'Cytokine-cytokine receptor interaction' pathway. In contrast, the VAE uniquely captured this critical upstream cascade (OR = 6.5) alongside the 'JAK-STAT signaling pathway' (OR = 10.2). The VAE achieved this by uniquely prioritizing critical upstream cytokine receptors, including *Lifr*, *Il13ra1*, and *Jak1*, which were entirely missed or de-prioritized by linear baselines (ST 1). Taken together, these results indicate that the VAE's non-linear representation reduces technical artifact carryover and mitigates over-reliance on MHC-dominated downstream signals, revealing a highly specific and topologically robust microglial aging network.

#### SR 2.2. Detailed Pathway Enrichment Comparison in Aging HSCs

Evaluation of the HSC trajectories revealed an analytical paradigm parallel to the microglial findings. While all methods successfully captured the downstream inflammatory signatures of age-related myeloid skewing (e.g., Leishmaniasis, Th1/Th2 cell differentiation), the linear models (PCA, ALS) were disproportionately driven by broad systemic inflammatory shifts. In contrast, the VAE retained the inflammatory component while additionally prioritizing putative upstream drivers of stem cell exhaustion as its top structural hits.

Specifically, the VAE isolated 'DNA replication' as a top biological hit, achieving a massive odds ratio (OR = 84.6) compared to PCA (OR = 52.0) and ALS (OR = 41.5). This pathway captured the comprehensive downregulation of the MCM helicase complex (*Mcm2*, *Mcm3*, *Mcm5*, *Mcm6*) and DNA polymerase (*Pold1*), which are essential for preventing replicative senescence. Furthermore, the VAE robustly prioritized the 'Hematopoietic cell lineage' pathway (OR = 10.6), accurately identifying the loss of critical upstream stemness and progenitor markers, including *Kit* (c-Kit) and *Flt3*, which were assigned lower statistical priority in linear models.

Crucially, the VAE strongly elevated pathways related to pre-leukemic transformation ('Acute myeloid leukemia', OR = 16.6 vs. PCA OR = 7.0), successfully capturing the severe depletion of the key stem cell regulators *Flt3* and *Kit*. Notably, the VAE successfully retained *Mpo* (Myeloperoxidase), a critical granulocyte-macrophage progenitor (GMP) marker essential for early myeloid commitment [34, 35], which was completely dropped by the PCA baseline. While excessive *Mpo* activity is associated with oxidative damage in stress or disease states,[36] its physiological downregulation in the aged trajectory reflects a critical loss of functional GMP commitment, contributing to age-related immunosenescence.[37] This demonstrates that the VAE's non-linear manifold can prioritize core molecular hallmarks of replicative senescence and progenitor exhaustion alongside the broader inflammatory remodeling that accompanies HSC aging.

#### SR 2.3. Forward–Reverse Target Symmetry and Clamp-Ablation Analysis (Full)

The GeroNetwork architecture predicts a topological hierarchy in which lineage- and replication-specific identity programs (Pillar I) act as deep upstream drivers, while the inflammatory and MHC remodeling cascade (Pillar III) emerges as a downstream, state-dependent consequence. A direct gene-level test of this hierarchy is whether the deep identity programs are more bidirectionally traversable under the learned aging trajectory than the downstream inflammatory programs. To address this, we compared consensus targets between GeroSimOld and GeroSimYoung after expressing both gene-effect signatures on the common young→old aging axis. Forward effects were computed as $\Delta^F_{aging} = \hat{\bar{\mathbf{x}}}_{old} - \bar{\mathbf{x}}_{young}$, whereas reverse effects were computed as $\Delta^R_{aging} = \bar{\mathbf{x}}_{old} - \hat{\bar{\mathbf{x}}}_{young}$. Genes were then classified as bidirectionally shared with sign-consistent aging-axis direction, forward-only, reverse-only, or shared with conflicting aging-axis sign (Figure 4A,B). Jaccard overlap was calculated as shared sign-consistent genes divided by the union of forward and reverse consensus target sets..

Under the standard clamped pipeline, VAE forward and reverse signatures shared a sign-consistent but incomplete core: 22 shared targets in microglia and 29 in HSCs, corresponding to a 38% Jaccard overlap in both tissues. No sign-inconsistent shared genes were detected in any model, tissue, or clamp condition (Figure 4A,B; red category empty in every panel), confirming that the learned GeroVector defines a sign-coherent bidirectional aging axis.

This partial overlap should not be interpreted as inferior reversibility relative to linear baselines; rather, it distinguishes the VAE from models whose same-axis forward–reverse behavior is constrained by shift-invariant or nearly linear reconstruction geometry. To establish this rigorously, we performed two complementary analyses designed to exclude alternative explanations for the VAE–linear gap. First, we performed a no-clamp ablation in which the non-negativity rectification was removed prior to consensus extraction (Figure 4B,C). ALS recovered to complete forward–reverse symmetry (Jaccard 61% → 100% Microglia; 70% → 100% HSC), confirming that the rectification was the *sole* source of asymmetry in this baseline, consistent with the algebraic fact that $\mathbf{X}_{young} + \mathbf{g}_{als}$ and $\mathbf{X}_{old} - \mathbf{g}_{als}$ produce exactly anti-symmetric pseudo-bulk shifts in the absence of clamping. PCA recovered partially (58% → 67% Microglia; 64% → 66% HSC), with the residual asymmetry reflecting the lossy, state-dependent nature of truncated linear projection: young and old cells project differently into the same fixed PCA basis, so the forward and reverse reconstruction residuals differ. The VAE was essentially unaffected by clamp removal (38% → 43% Microglia; 38% → 35% HSC), identifying intrinsic non-linear encoder–decoder geometry, rather than the shared rectification, as the dominant source of the VAE's direction-specific recovery.

Second, to exclude consensus thresholding as the source of the asymmetry, we computed a continuous, threshold-free forward–reverse $\log_2$FC concordance (Pearson $r$) across all HVGs per LOPO fold (Figure 4D,E). Because this metric operates on the full HVG space without any top-100 or 12/12 selection step, it cannot be affected by thresholded target selection by construction. The continuous metric reproduced the same model ordering and the same clamp-ablation pattern: under the standard clamp, ALS and PCA showed high same-axis forward–reverse concordance (Microglia: ALS $r = 0.84$, PCA $r = 0.82$; HSC: ALS $r = 0.90$, PCA $r = 0.87$), while the VAE remained substantially lower (Microglia: $r = 0.59$; HSC: $r = 0.65$); after clamp removal, ALS reached the algebraic ceiling $r = 1.00$ in both tissues, PCA increased to $r = 0.91$, and the VAE remained at $r = 0.61/0.65$. Together, the no-clamp ablation and the all-HVG continuous concordance rule out both output rectification and thresholded target selection as the cause of the VAE's lower forward–reverse overlap. In contrast, ALS reaches perfect no-clamp symmetry precisely because direct expression-space translation cannot encode state-dependent return paths; PCA approaches but does not reach this ceiling because its lossy projection introduces only a mild form of state dependence; the VAE retains substantial direction-specificity because its non-linear encoder–decoder geometry is genuinely state-dependent by design.

The biological structure of the VAE partitions was concordant with the GeroNetwork hierarchy. The HSC shared bidirectional core was dominated by replication and progenitor-identity machinery, comprising the MCM helicase complex (*Mcm2*, *Mcm3*, *Mcm5*, *Mcm6*), the DNA polymerase *Pold1*, and the stemness regulator *Flt3*, together with protein-folding/ER components (*P4hb*, *Pdia4*, *Serp1*) and nuclear-envelope/chromatin regulators (*Xab2*, *Sun2*, *Ddx54*). This is precisely the Pillar I program topologically anchored by the *E2f/Tp53/Mycn* regulatory module, indicating that the deepest replicative–identity component of HSC aging is the most state-independent and bidirectionally traversable layer of the learned trajectory. The HSC forward-only set was instead dominated by the Pillar III inflammatory remodeling cascade: MHC class II components (*H2-Aa*, *H2-Ab1*, *H2-Eb1*, *Cd74*), AP-1 / immediate-early / NF-$\kappa$B regulators (*Jun*, *Junb*, *Egr1*, *Ier3*, *Dusp1*, *Nfkbia*, *Rela*), *Apoe*, and the surface stem-cell markers *Kit*, *Mki67*, and *Mpo*. In microglia, the same dissociation appeared more partially: the shared core contained the Pillar I cytokine-receptor / AP-1 identity axis (*Jun*, *Junb*, *Lifr*, *Il13ra1*, *Jak1*), constitutive MHC class I components (*H2-D1*, *H2-K1*), and the classical microglial aging hallmarks *Apoe* and *Lyz2*, while the forward-only set captured MHC class II (*Cd74*, *H2-Ab1*, *H2-Q6*) and homeostatic phagocytic receptors (*C3ar1*, *Rab31*). Across both tissues, MHC class I components were bidirectionally reversible while MHC class II was forward-specific, reflecting the biological distinction between constitutive class I presentation and the inducible, specialized class II antigen-presentation program that engages during aging.

Because forward-only and reverse-only assignments depend on strict 12/12 consensus filtering at a fixed top-100 rank cutoff, individual genes near this boundary may shift classification through fold-to-fold ranking variance rather than true direction-specificity. We therefore interpret these categories primarily at the level of coherent modules—for example, the joint co-classification of all four MCM subunits and *Pold1* as shared in HSC, or the joint co-classification of the entire MHC-II + AP-1 + NF-$\kappa$B block as forward-only—rather than over-interpreting any individual boundary gene. Because GeroTargets were defined using a strict 12/12 LOPO consensus threshold, model-specific set sizes reflect both biological reproducibility and each model's reconstruction geometry; therefore, overlap was evaluated using Jaccard normalization rather than raw shared counts alone.

Taken together, this gene-level analysis provides convergent support for the GeroNetwork tripartite hierarchy: Pillar I lineage and replicative identity programs constitute the deep, bidirectionally traversable core of the learned aging trajectory, while Pillar III inflammatory and antigen-presentation programs emerge as state-dependent, direction-specific downstream consequences. The lower VAE forward–reverse overlap should therefore be interpreted cautiously: rather than indicating a simple failure of reversibility, it may provide a diagnostic readout of state-dependent manifold geometry that reflects the depth structure of the aging trajectory. The quantitative summary of this analysis is presented in main-text Figure 4.

### SDi 3. Supplementary Discussion

#### SDi 3.1. Forward–Reverse Asymmetry Reveals State-Dependent Non-linear Manifold Geometry

A central observation from our bidirectional analysis is that the VAE exhibits substantially lower forward–reverse gene overlap than the linear baselines, and that this gap persists after removing both the non-negativity rectification and the consensus threshold. The clamp ablation and the threshold-free continuous concordance metric together rule out output rectification and thresholded target selection as explanations, leaving non-linear, state-dependent decoder geometry as the remaining source of the asymmetry. We interpret this lower overlap not as inferior reversibility, but as the diagnostic signature of a model that explicitly encodes state-dependent biology.

The three models we benchmarked occupy a clean ladder of state dependence under the common aging-axis convention. ALS is state-independent by construction: the same expression-space vector is added in the forward direction and subtracted in the reverse direction. Therefore, after reverse effects are re-expressed as real old minus GeroSimYoung and the shared non-negativity rectification is removed, the forward and reverse gene effects converge to the same aging-axis vector. PCA introduces a mild form of state dependence through its lossy, finite-dimensional reconstruction: young and old cells project differently into the same fixed basis, and the reconstruction residuals therefore differ between directions, but this contribution is small (continuous $r \approx 0.91$ after clamp removal). The VAE is strongly state-dependent:

$$\hat{\mathbf{X}}^F_{old} = \max\left(0,\ \mathrm{De}\left[\mathrm{En}(\mathbf{X}_{young}) + \mathbf{g}\right]\right), \qquad \hat{\mathbf{X}}^R_{young} = \max\left(0,\ \mathrm{De}\left[\mathrm{En}(\mathbf{X}_{old}) - \mathbf{g}\right]\right). \tag{SEq 27}$$

Because these two simulations begin from different regions of the latent manifold and are decoded through locally distinct non-linear neighborhoods, a fixed latent displacement $\mathbf{g}$ can produce different gene-space effects depending on the starting cellular state. Encoder lossiness compounds this effect: gene-level structure compressed away by the encoder cannot be perfectly restored by the decoder during reverse traversal. Consequently, after both directions are expressed on the common aging axis,

$$\Delta^F_{aging} = \overline{\hat{\mathbf{x}}}^F_{old} - \overline{\mathbf{x}}_{young}, \qquad \Delta^R_{aging} = \overline{\mathbf{x}}_{old} - \overline{\hat{\mathbf{x}}}^R_{young}, \tag{SEq 28}$$

a shared bidirectional core can coexist with direction-specific peripheral programs.

Importantly, this asymmetry is a feature rather than a defect. Linear baselines achieve full forward/reverse symmetry only because they cannot encode state-dependent biology in the first place; the VAE's lower overlap is the price—and the diagnostic value—of an explicitly state-dependent generative model. Biologically, the asymmetry is moreover plausible: once a cell has acquired the aged inflammatory state (Pillar III), simply negating the population-level aging vector does not deterministically restore every gene to its young state, because the inflammatory program engages on top of, and downstream of, the more state-independent identity machinery (Pillar I). The deeper a program lies in the regulatory hierarchy, the less its reverse traversal depends on the specific aged-cell starting state, and the more bidirectionally reversible it appears at the gene level. This is precisely what the consensus partition shows: the MCM helicase complex, *Pold1*, and *Flt3* (Pillar I replication/stemness identity) constitute the bidirectionally shared core in HSCs, while the entire MHC class II + AP-1 + NF-$\kappa$B module (Pillar III) is recovered only in the forward direction.

This interpretation reframes the partial overlap as a depth-of-hierarchy readout rather than a model limitation. Genes recovered with the same sign after common aging-axis alignment define the most state-independent component of the trajectory; genes recovered in only one direction are state-dependent satellites of that core. Read alongside the GeroNetwork topology, this dissociation provides converging gene-level support for prioritizing Pillar I master regulators (*E2f*, *Mycn* for HSC replication; *Spi1*, *Tal1* for microglial identity) as therapeutic candidates whose modulation is more likely to translate into broad, coherent reversal of aged cellular programming than targeting downstream inflammatory hubs alone—a hypothesis consistent with the therapeutic argument developed in the main-text Discussion. We interpret the reverse simulation throughout as a computational analog of rejuvenation-like latent traversal, not as evidence of experimentally validated biological rejuvenation.

Finally, we note that even within the VAE framework, the perturbation is applied as a first-order linear shift in latent space; higher-order, curved, or time-varying latent trajectories may modify the precise structure of the direction-specific component. Exploring such trajectories—for instance via Neural Ordinary Differential Equations directly in the latent space, as noted in the main-text Limitations—would test whether the residual direction-specificity is intrinsic to the underlying biology or partially attributable to the latent-linearity assumption of the present implementation.

### SF 4. Supplementary Figures

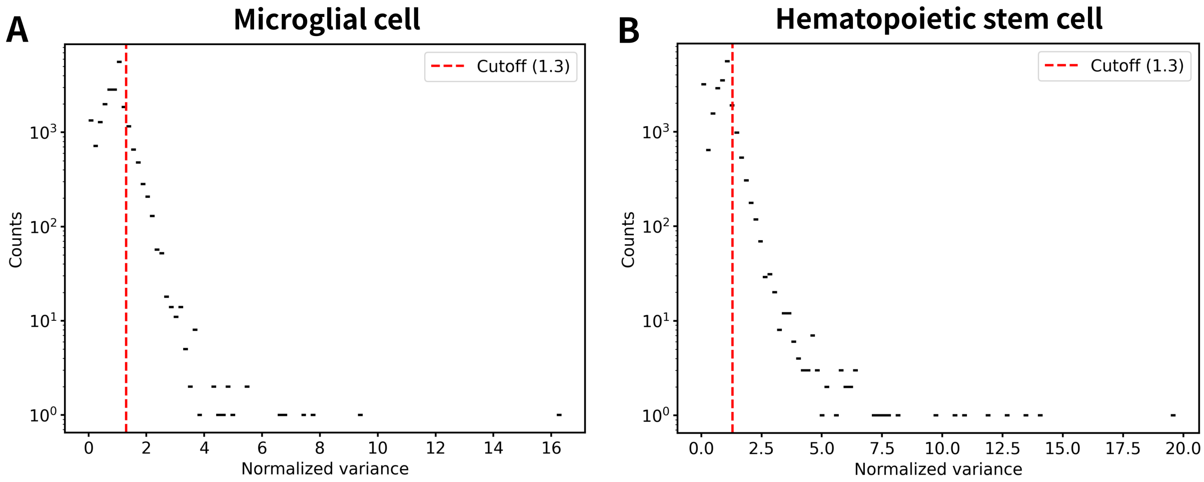


Figure 1. **Distribution of normalized gene variances and feature selection.** Scatter plot illustrating the normalized variance for all genes in the primary male cohort. A strict normalized variance cutoff of 1.3 (indicated by the threshold line) was applied to objectively retain Highly Variable Genes (HVGs) while discarding stable background transcripts, yielding the finalized geometric feature space for downstream latent modeling.

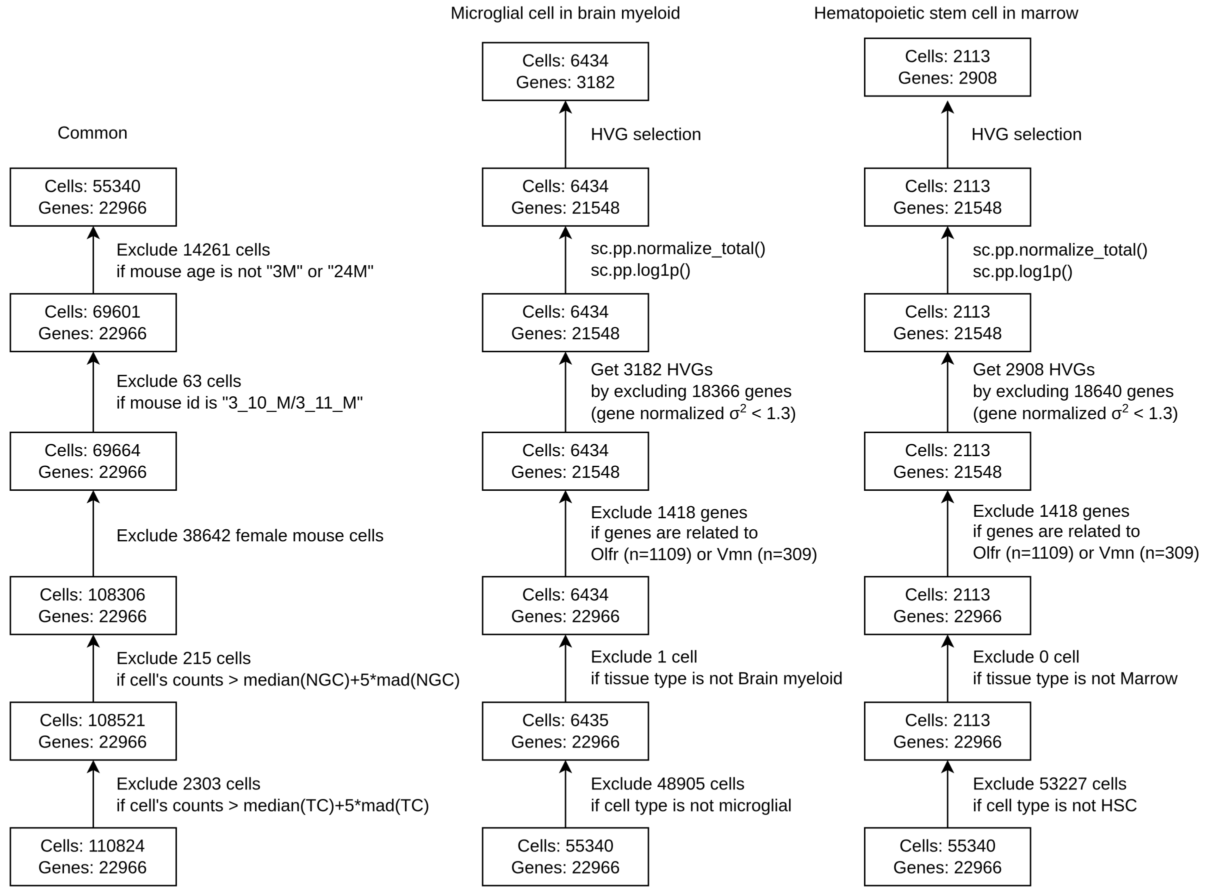


Figure 2. **Detailed single-cell transcriptomic preprocessing and feature alignment pipeline.** Overview of the computational workflow applied to the Tabula Muris Senis dataset. The flowchart details initial quality control (QC) filtering, library-size normalization, log-transformation ($\ln(x+1)$), and the strict geometric feature subsetting required to map Out-Of-Distribution (OOD) cohorts (18-month males and all females) onto the identical Highly Variable Gene (HVG) coordinate space defined by the primary male training cohort. TC, Total Counts; NGC, Number of Genes by Counts; MAD, Median Absolute Deviation

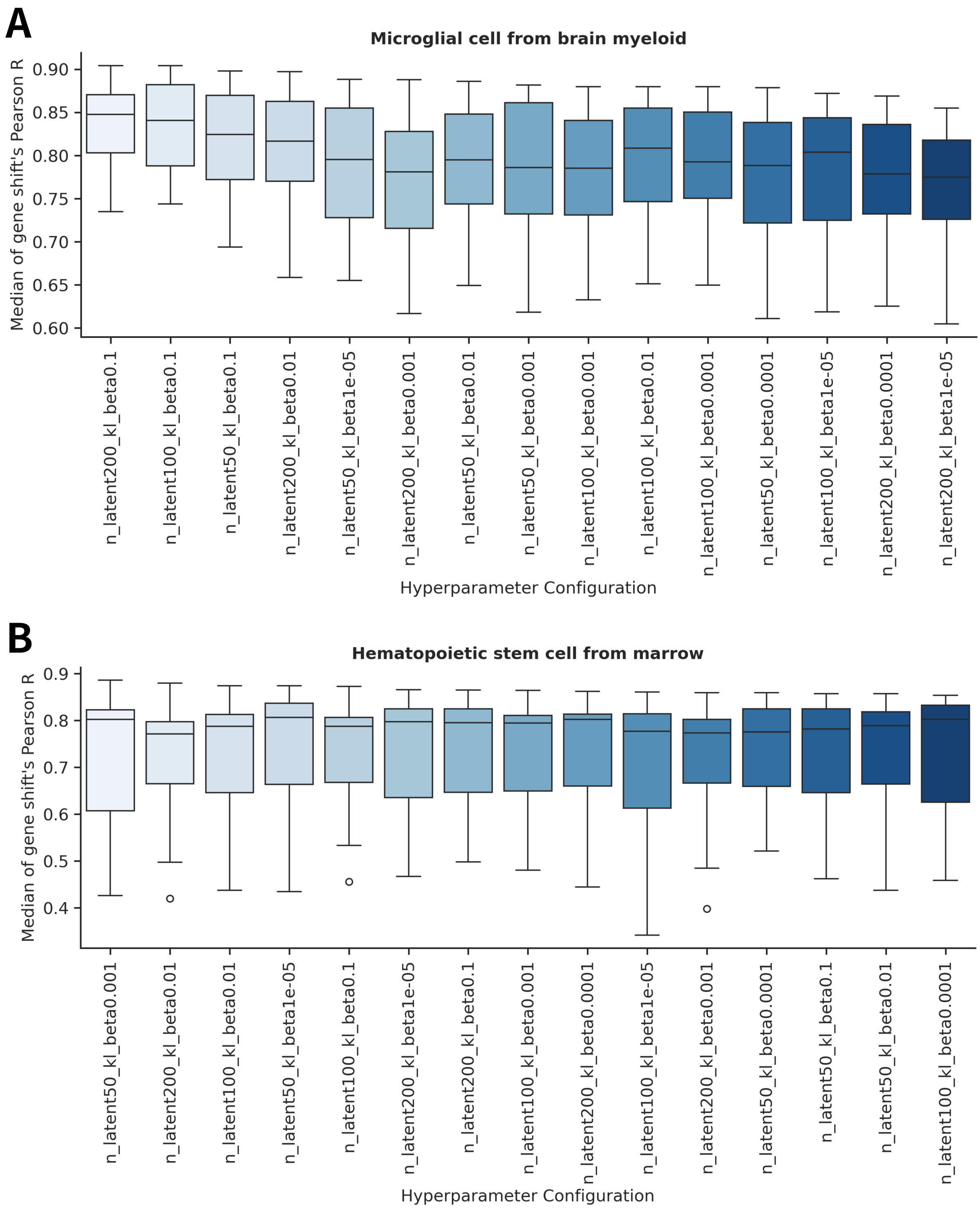


Figure 3. **Ablation study of Variational Autoencoder (VAE) hyperparameters.** Grid search evaluating the stability of the GeroSimulator across varying latent dimensionalities ($N_{latent\_dim} \in \{50, 100, 200\}$) and Kullback-Leibler (KL) divergence penalty scaling factors ($\beta_{KL} \in \{10^{-5}, 10^{-4}, 10^{-3}, 10^{-2}, 10^{-1}\}$). Boxes display the resulting trajectory prediction accuracy as measured by the transcriptomic shift correlation ($R_{shift}$). The conservatively selected configuration ($N_{latent\_dim} = 100$, $\beta_{KL} = 10^{-3}$) demonstrates an optimal balance, ensuring robust directional accuracy without artificially crushing transcriptomic heterogeneity through over-regularization.

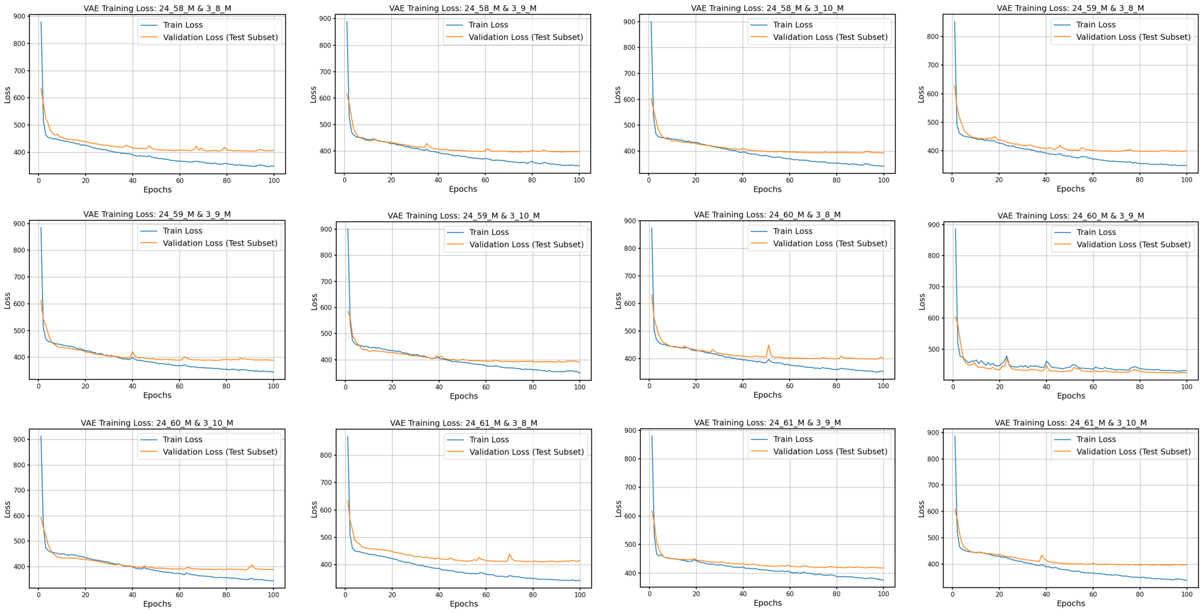


Figure 4. **Training and validation loss trajectories for the Microglia GeroSimulator.** Convergence plots detailing the composite objective function (MSE reconstruction loss and KL divergence regularization) over 100 training epochs for the microglial cohort. The curves demonstrate stable optimization dynamics and convergence across the leave-one-pair-out cross-validation folds, with no indications of severe overfitting.

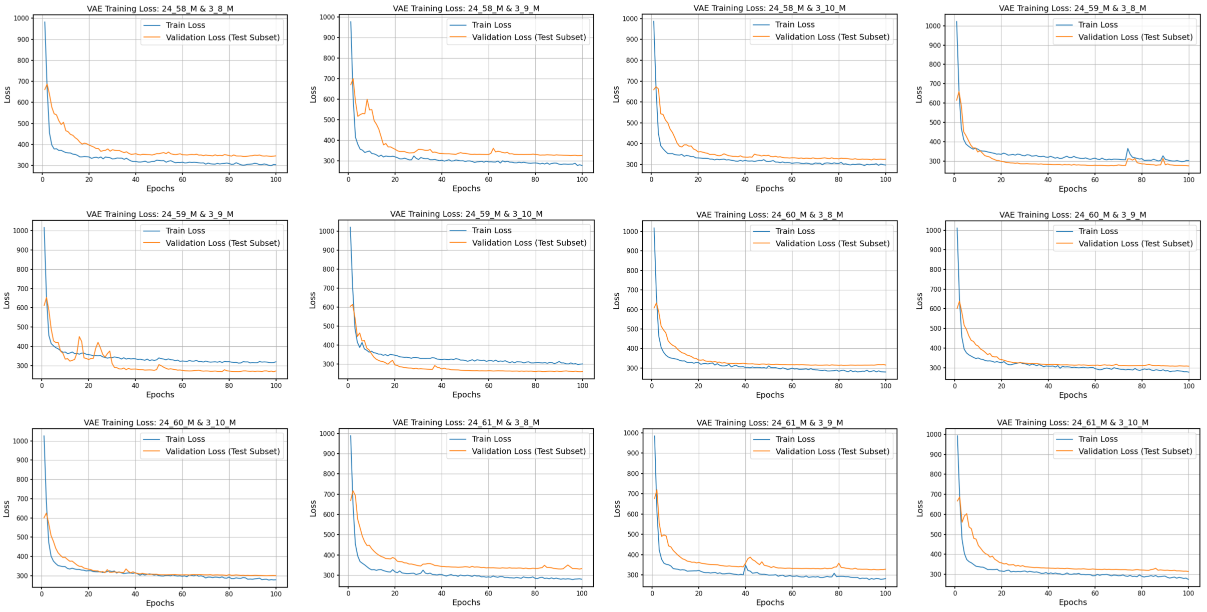


Figure 5. **Training and validation loss trajectories for the Hematopoietic Stem Cell (HSC) GeroSimulator.** Convergence plots detailing the composite objective function over 100 training epochs for the HSC cohort. Similar to the microglial model, the training dynamics exhibit stable convergence across the cross-validation folds without signs of mode collapse or significant generalization gaps.

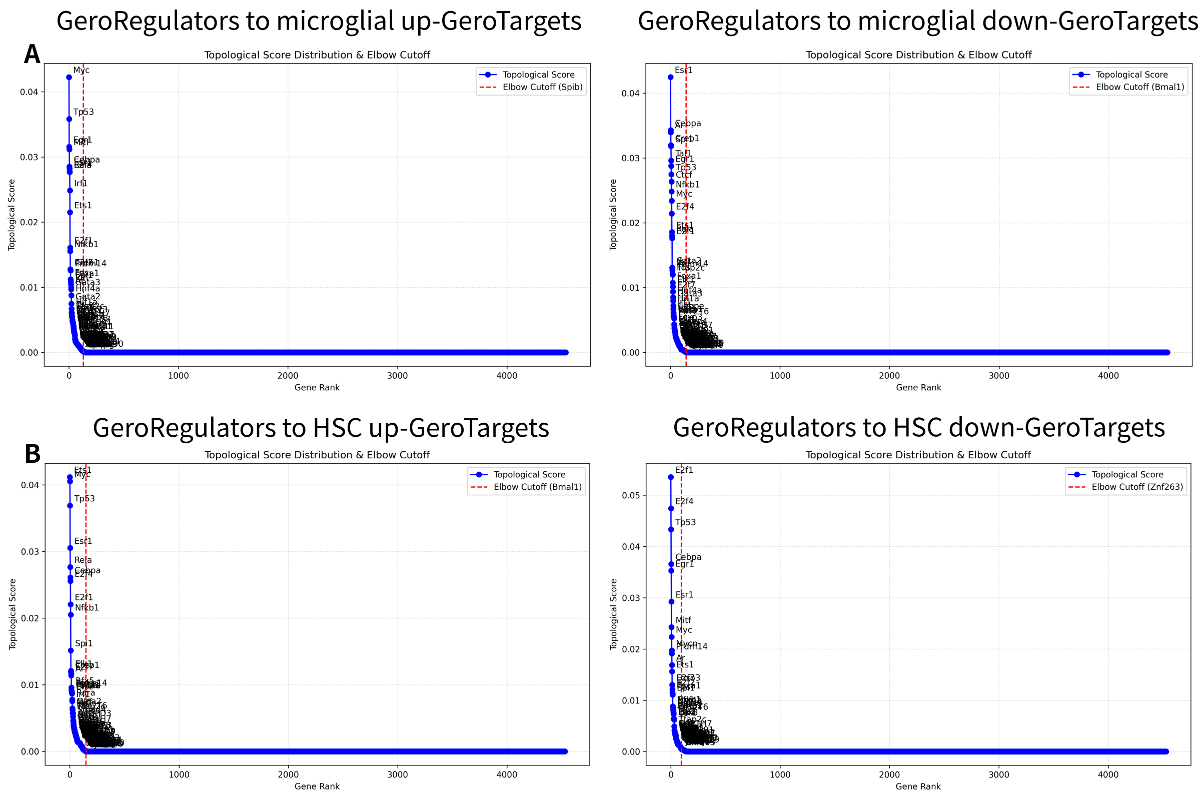


Figure 6. **Objective dynamic thresholding of Topological GeroRegulators via the Kneedle algorithm.** Distribution of reverse-directed Personalized PageRank (PPR) topological scores for candidate upstream master regulators. Genes are ranked along the x-axis in descending order of structural centrality. The Kneedle algorithm was applied to mathematically define the elbow point of the curve (indicated by the threshold), allowing for the objective exclusion of low-impact background network noise prior to the application of tissue-adaptive structural filtering constraints.

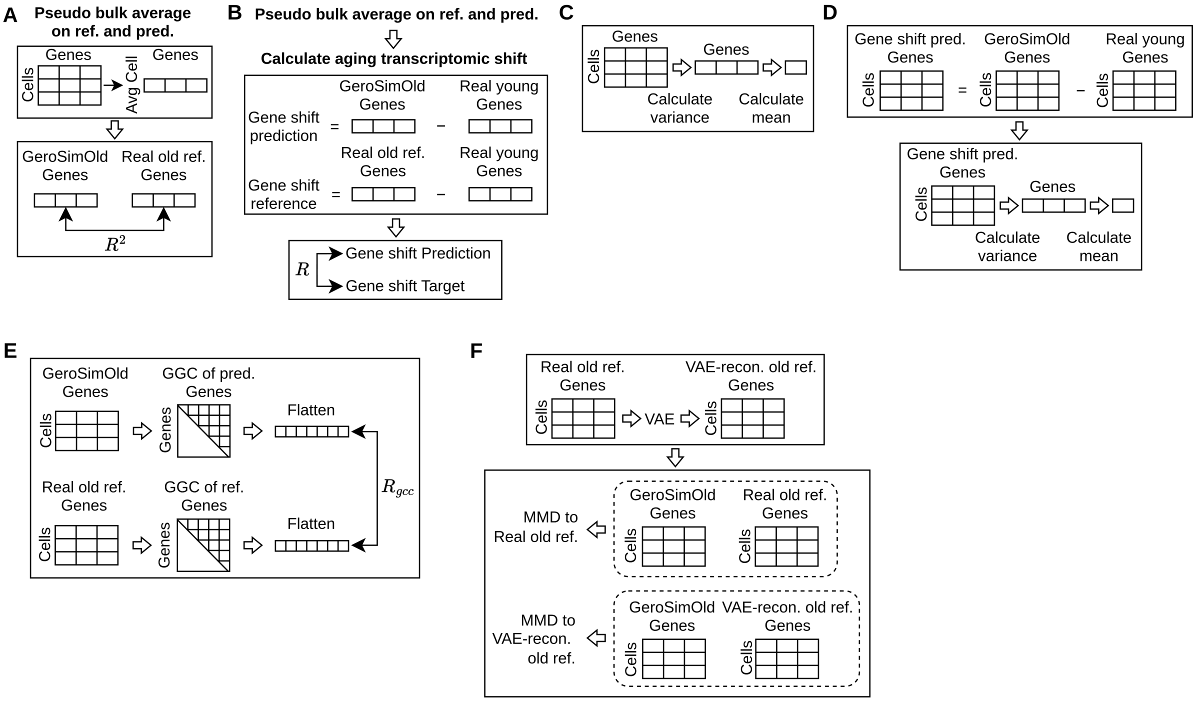


Figure 7. **Overview of quantitative evaluation metrics for longitudinal trajectory simulation.** (A) Coefficient of Determination ($R^2$) for Macroscopic Alignment. The predicted aged expression matrix and the empirical biological old target matrix are pseudo-bulked by averaging across the cellular dimension. The $R^2$ score is calculated between the resulting macroscopic gene expression vectors to evaluate baseline deterministic accuracy. (B) Pearson Correlation of Transcriptomic Shifts ($R_{\text{shift}}$). Pseudo-bulked transcriptomic shift vectors are isolated by subtracting the empirical biological young baseline from both the predicted and target old states. The Pearson correlation ($R$) between these isolated shift vectors measures the directional accuracy of the dynamic aging trajectory, removing invariant cell-type markers. (C) Mean of Gene Variances ($\bar{\sigma}^2$) for Transcriptomic Denoising. To evaluate structural noise compression, the variance of each gene is calculated across the simulated cell population, and the global mean of these gene variances is computed. (D) Mean of Gene Shift Variances ($\bar{\sigma}^2_{\text{shift}}$) for Trajectory Heterogeneity. A cell-wise transcriptomic shift matrix is established by subtracting the biological young baseline from the predicted aged cells. The variance of these individual shifts is calculated per gene, and the global mean is computed to evaluate the retention of natural cell-to-cell stochasticity and to detect algorithmic mode collapse. (E) Gene-Gene Correlation ($R_{\text{ggc}}$) for Network Denoising. Pairwise gene-gene correlation (GGC) matrices are generated for both the predicted and empirical target aged states. The strictly upper-triangular elements are flattened into one-dimensional network edge vectors, and their Pearson correlation ($R_{\text{ggc}}$) quantifies the global preservation of multivariate regulatory networks. (F) Maximum Mean Discrepancy (MMD) for Global Manifold Alignment. The empirical biological old target is passed through the VAE latent bottleneck to establish a mathematically denoised biological target manifold. The unbiased empirical MMD of the predicted aged state is subsequently evaluated against both the raw (noisy) target and this denoised target to quantify global topological fidelity.

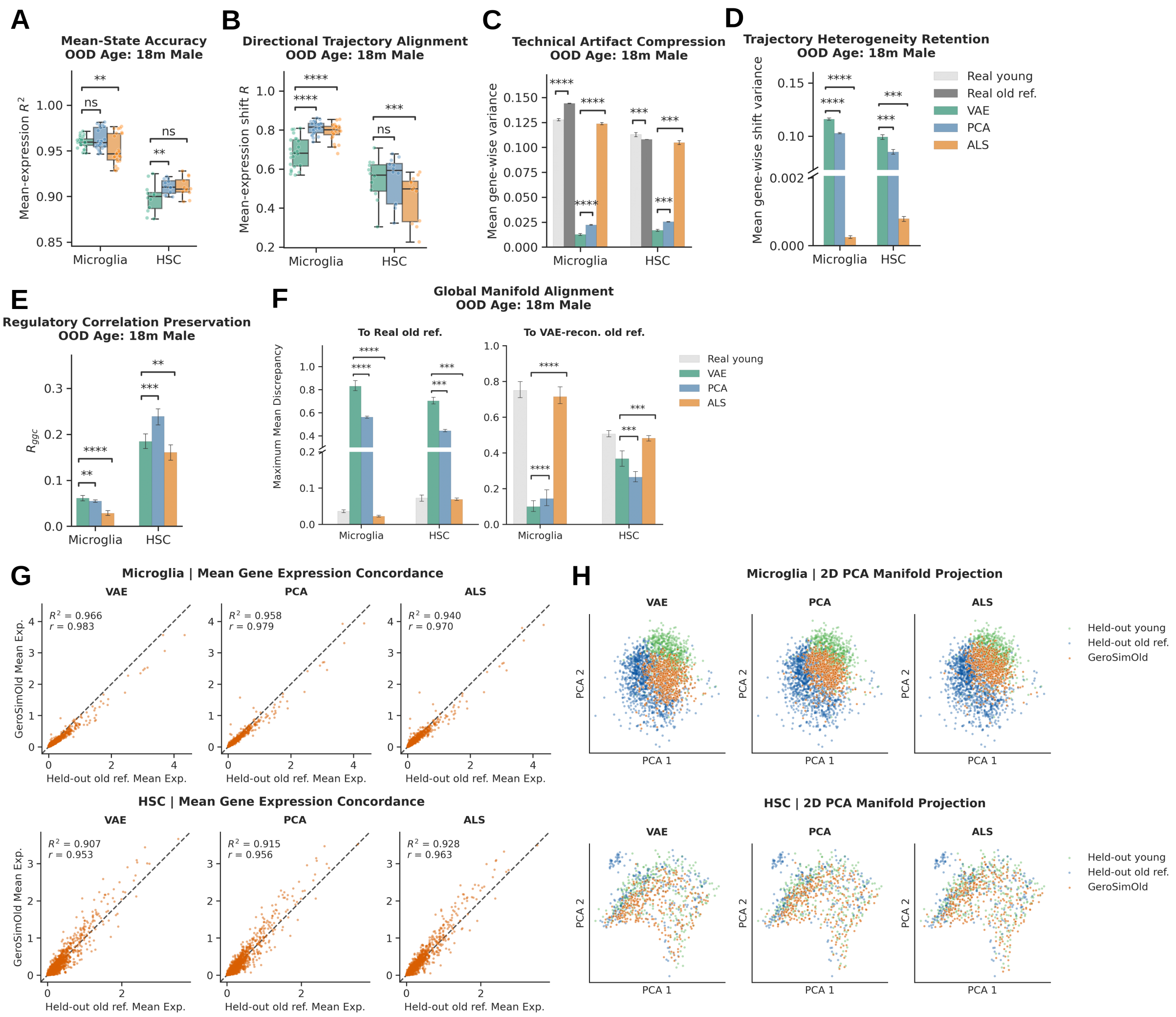


Figure 8. **OOD validation across unseen chronological age.** Performance when interpolating the 24-month learned vector to predict the unseen 18-month male state; metrics correspond to Figure 1. The VAE demonstrates robust temporal scalability while maintaining artifact compression, heterogeneity preservation, and manifold alignment. Panels (G,H) display representative simulations using the 3_10_M/18_53_M pair for HSCs (fold 12) and 3_9_M/18_45_M for microglia (fold 15).

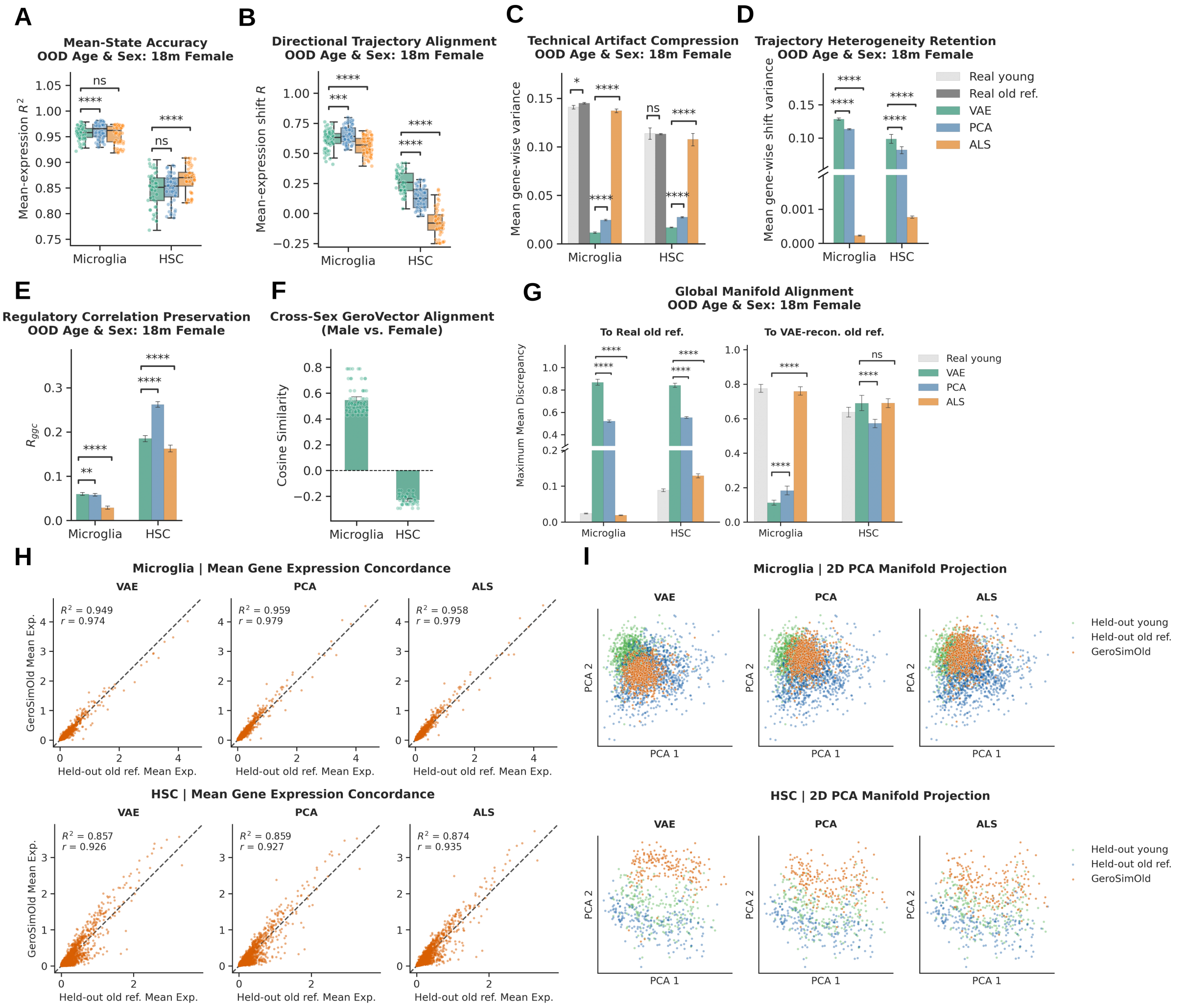


Figure 9. **OOD validation across unseen chronological age and biological sex.** Performance when applying the male-derived vector to an unseen young female baseline. The VAE maintains superior directional accuracy ($R_{shift}$) despite biological orthogonality, but reveals decoder limitations in MMD alignment when reconstructing highly dimorphic baseline topologies (HSCs). Panels (H,I) display representative simulations using the 3_39_F/18_46_F pair for microglia (fold 14) and 3_39_F/18_47_F for HSCs (fold 20).

### ST 5. Supplementary Tables

Table 1. **Pathway Enrichment Comparison in Aging Microglia.** GSEA reveals that while linear models (PCA, ALS) are disproportionately influenced by downstream MHC-associated signals, the VAE non-linear manifold additionally prioritizes upstream lineage-specific regulators.
*Bold numbers indicate the highest Odds Ratio (OR) across models for a given pathway. Bold key enriched genes denote targets captured by the VAE that were missed by at least one of the linear baseline models.

| Category | Pathway / GO Term | VAE | PCA | ALS | Key Enriched Genes |
|---|---|---|---|---|---|
| **Lineage & Differentiation** | Osteoclast differentiation | $8.7 \times 10^{-3}$ OR: 14.9 | $1.7 \times 10^{-4}$ **OR: 17.7** | $5.5 \times 10^{-3}$ OR: 8.1 | *Jun, Junb, Jak1* |
| | Leishmaniasis | $7.1 \times 10^{-3}$ **OR: 16.8** | $5.6 \times 10^{-4}$ OR: 15.6 | $5.9 \times 10^{-4}$ OR: 12.0 | *Jun,* ***H2-Ab1****, Jak1* |
| **Upstream Regulation** | JAK-STAT signaling | $1.8 \times 10^{-2}$ **OR: 10.2** | $1.5 \times 10^{-2}$ OR: 6.8 | $2.1 \times 10^{-2}$ OR: 5.3 | *Lifr, Il13ra1, Jak1* |
| | Cytokine receptor interaction | $4.3 \times 10^{-2}$ **OR: 6.5** | N/A N/A | N/A N/A | ***Lifr, Il13ra1, Tnfsf13b*** |
| **MHC / Downstream Inflammatory Response** | Allograft rejection | $4.0 \times 10^{-4}$ **OR: 30.7** | $3.0 \times 10^{-4}$ OR: 19.8 | $3.9 \times 10^{-4}$ OR: 15.6 | *H2-Q6, H2-K1, H2-D1,* ***H2-Ab1*** |
| | Autoimmune thyroid disease | $4.0 \times 10^{-4}$ **OR: 30.7** | $3.0 \times 10^{-4}$ OR: 19.8 | $3.9 \times 10^{-4}$ OR: 15.6 | *H2-Q6, H2-K1, H2-D1,* ***H2-Ab1*** |

Table 2. **Pathway Enrichment Comparison in Aging HSCs.** GSEA indicates that linear models (PCA, ALS) are disproportionately skewed by downstream inflammatory myeloid skewing. Conversely, the VAE manifold isolates the upstream drivers of stem cell exhaustion, particularly replicative senescence and progenitor marker loss.
*Bold numbers indicate the highest OR across models for a given pathway. Bold key enriched genes denote targets captured by the VAE that were missed by at least one of the linear baseline models.

| Category | Pathway / GO Term | VAE | PCA | ALS | Key Enriched Genes |
|---|---|---|---|---|---|
| **Stemness & Replication** | DNA replication | $2.5 \times 10^{-5}$ **OR: 84.6** | $3.9 \times 10^{-5}$ OR: 52.0 | $1.1 \times 10^{-4}$ OR: 41.5 | *Pold1, Mcm2, Mcm3, Mcm5, Mcm6* |
| | Hematopoietic cell lineage | $1.1 \times 10^{-3}$ **OR: 10.6** | $1.7 \times 10^{-4}$ OR: 9.5 | $1.1 \times 10^{-4}$ OR: 8.9 | *H2-Eb1, Flt3, Kit, Cd9, H2-Aa, H2-Ab1* |
| **Pre-Leukemic** | Acute myeloid leukemia | $2.3 \times 10^{-3}$ **OR: 16.6** | $4.5 \times 10^{-2}$ OR: 7.0 | $1.4 \times 10^{-2}$ OR: 8.2 | *Flt3, Kit,* ***Mpo****, Rela* |
| **Downstream Myeloid Skewing** | Leishmaniasis | $3.1 \times 10^{-5}$ **OR: 20.4** | $6.0 \times 10^{-6}$ OR: 18.3 | $9.7 \times 10^{-6}$ OR: 14.5 | *Nfkbia, H2-Eb1, Jun, Fos, H2-Aa, Rela, H2-Ab1* |
| | Th1/Th2 differentiation | $1.2 \times 10^{-4}$ **OR: 14.6** | $3.0 \times 10^{-5}$ OR: 12.7 | $9.7 \times 10^{-6}$ OR: 11.8 | *Nfkbia, H2-Eb1, Jun, Fos, H2-Aa, Rela, H2-Ab1* |

### Terms

#### Abbreviations

**ALS**: Algebraic Linear Shift
**BH-FDR**: Benjamini-Hochberg False Discovery Rate
**ER**: Endoplasmic Reticulum

**FACS**: Fluorescence-Activated Cell Sorting
**GGC**: Gene-Gene Correlation
**GMP**: Granulocyte-Macrophage Progenitor
**GO**: Gene Ontology
**GRN**: Gene Regulatory Network
**GSEA**: Gene Set Enrichment Analysis
**HSC**: Hematopoietic Stem Cell
**HVG**: Highly Variable Gene
**KEGG**: Kyoto Encyclopedia of Genes and Genomes
**KL**: Kullback-Leibler (divergence)
$\log_2$ **FC**: Logarithm base 2 Fold Change
**LOPO**: Leave-One-Pair-Out
**MAD**: Median Absolute Deviation
**MGI**: Mouse Genome Informatics
**MHC**: Major Histocompatibility Complex
**MMD**: Maximum Mean Discrepancy
**MSE**: Mean Squared Error
**NGC**: Number of Genes by Counts
**ODE**: Ordinary Differential Equation
**OOD**: Out-Of-Distribution
**OR**: Odds Ratio
**PCA**: Principal Component Analysis
**PCC**: Pearson Correlation Coefficient
**PPR**: Personalized PageRank
**QC**: Quality Control
**RBF**: Radial Basis Function
**RKHS**: Reproducing Kernel Hilbert Space
**SA**: Supplementary Algorithm
**scRNA-seq**: Single-cell RNA sequencing
**SDi**: Supplementary Discussion
**SEq**: Supplementary Equation
**SF**: Supplementary Figure
**SM**: Supplementary Methods
**SR**: Supplementary Results
**ST**: Supplementary Table
**TC**: Total Counts
**VAE**: Variational Autoencoder

### Mathematical Notation

#### Constants and Hyperparameters

$N_{hvg}$: Constant. The number of highly variable genes retained during preprocessing
$N_{cells}$: Constant. The number of cells in a given matrix
$N_{latent\_dim}$: Constant. The dimensionality of the VAE latent space (and PCA subspace)
$\beta_{KL}$: Constant. Scaling factor for the Kullback-Leibler divergence regularization term
$p_{dropout}$: Constant. Dropout probability regularization in the VAE model
$\epsilon_{bn}$: Constant. Epsilon value for batch normalization in the VAE model
$\alpha$: Constant. Temporal scaling factor for chronological aging interpolation

#### Expression Space (Vectors and Matrices)

$\mathbf{X} \in \mathbb{R}^{N_{cells} \times N_{hvg}}$: Matrix. Empirical preprocessed gene expression of cells
$\hat{\mathbf{X}} \in \mathbb{R}^{N_{cells} \times N_{hvg}}$: Matrix. Predicted (simulated) gene expression of cells (e.g., GeroSimOld in the forward

direction, or GeroSimYoung in the reverse direction)
$\overline{\mathbf{x}}, \hat{\overline{\mathbf{x}}} \in \mathbb{R}^{N_{hvg}}$: Vectors. Empirical and predicted pseudo-bulked mean expression vectors
$\mu_{\overline{\mathbf{x}}}$: Scalar. Global mean of the target expression vector across the evaluated gene subset
$\hat{\boldsymbol{\Delta}}, \boldsymbol{\Delta} \in \mathbb{R}^{N_{hvg}}$: Vectors. Predicted and empirical mean transcriptomic shift vectors (aged minus young centroid)
$\Delta^F_{aging}, \Delta^R_{aging} \in \mathbb{R}^{N_{hvg}}$: Vectors. Forward and reverse mean transcriptomic shift vectors expressed on the common young→old aging axis, defined as $\Delta^F_{aging} = \hat{\overline{\mathbf{x}}}_{old} - \overline{\mathbf{x}}_{young}$ (GeroSimOld − real young) and $\Delta^R_{aging} = \overline{\mathbf{x}}_{old} - \hat{\overline{\mathbf{x}}}_{young}$ (real old − GeroSimYoung)
$\log_2 \mathrm{FC}^F_{aging}, \log_2 \mathrm{FC}^R_{aging}$: Vectors/Scalars. Forward and reverse aging-axis log-fold changes, obtained by dividing $\Delta^F_{aging}$ and $\Delta^R_{aging}$ by $\ln 2$
$\hat{\mathbf{X}}^F_{old}, \hat{\mathbf{X}}^R_{young} \in \mathbb{R}^{N_{cells} \times N_{hvg}}$: Matrices. Forward-simulated aged profiles (GeroSimOld) and reverse-simulated young profiles (GeroSimYoung)
$\hat{\boldsymbol{\Delta}} \in \mathbb{R}^{N_{cells} \times N_{hvg}}$: Matrix. Cell-wise predicted transcriptomic shift matrix
$\mathbf{g}_{als} \in \mathbb{R}^{N_{hvg}}$: Vector. Algebraic Linear Shift aging trajectory vector in the original gene expression space

#### Latent Space (Vectors and Matrices)

$\mathbf{Z} \in \mathbb{R}^{N_{cells} \times N_{latent\_dim}}$: Matrix. Encoded gene expression of cells in the latent (or PCA) space
$\mathbf{g} \in \mathbb{R}^{N_{latent\_dim}}$: Vector. GeroVector; the fold-specific, population-level transcriptomic aging direction in the VAE latent space
$\mathbf{g}_{pca} \in \mathbb{R}^{N_{latent\_dim}}$: Vector. Aging trajectory direction in the linear Principal Component subspace
$\mathbf{g}^{train}_{fold}, \mathbf{g}^{eval}_{fold} \in \mathbb{R}^{N_{latent\_dim}}$: Vectors. Fold-specific latent aging directions (aged−young, i.e. target−source, centroid difference) for the fixed primary male training cohort and for a given held-out or OOD evaluation cohort, respectively, both encoded by the same fold-specific primary male forward-trained VAE
$\overline{\mathbf{z}}_m \in \mathbb{R}^{N_{latent\_dim}}$: Vector. Cell-level centroid (mean latent vector) isolated for a specific mouse $m$
$\overline{\mathbf{z}}_{global} \in \mathbb{R}^{N_{latent\_dim}}$: Vector. Unweighted mouse-level global centroid in the latent space

#### Evaluation Metrics (Variances and Networks)

$R^2$: Scalar. Coefficient of Determination measuring macroscopic alignment of mean expression profiles
$R_{shift}$: Scalar. Pearson correlation coefficient between the predicted ($\hat{\boldsymbol{\Delta}}$) and empirical ($\boldsymbol{\Delta}$) transcriptomic shift vectors
$\hat{\sigma}^2_j$: Scalar. Absolute variance of gene $j$ across simulated cells
$\overline{\hat{\sigma}}^2$: Scalar. Mean absolute gene variance strictly across the $N_{hvg}$ retained features
$\hat{\sigma}^2_j(\hat{\boldsymbol{\Delta}})$: Scalar. Variance of the transcriptomic shifts for individual gene $j$
$\overline{\hat{\sigma}}^2_{shift}$: Scalar. Mean shift variance across the $N_{hvg}$ retained features
$\hat{\mathbf{C}}, \mathbf{C} \in \mathbb{R}^{N_{hvg} \times N_{hvg}}$: Matrices. Predicted and empirical pairwise gene-gene Pearson correlation matrices
$\hat{\boldsymbol{e}}, \boldsymbol{e}$: Vectors. Flattened, one-dimensional network edge vectors containing strictly upper-triangular elements of $\hat{\mathbf{C}}$ and $\mathbf{C}$
$R_{ggc}$: Scalar. Pearson correlation coefficient between flattened network edge vectors $\hat{\boldsymbol{e}}$ and $\boldsymbol{e}$
$\cos(\theta_{fold})$: Scalar. Per-fold cosine similarity between $\mathbf{g}^{train}_{fold}$ and $\mathbf{g}^{eval}_{fold}$, quantifying directional alignment of the learned aging axis between the primary male training cohort and each evaluation condition

#### Topology, Algorithms, and Sets

$\mathrm{En}(\cdot)$: Function. The encoder of the trained Variational Autoencoder
$\mathrm{De}(\cdot)$: Function. The decoder of the trained Variational Autoencoder
$\mathrm{PCA}(\cdot)$: Function. Projection of expression data into the fitted PCA subspace
$\mathrm{PCA}^{-1}(\cdot)$: Function. Inverse projection from the PCA subspace back to the original gene expression space
$\mathcal{D}$: Dataset of single-cell RNA-seq profiles containing associated metadata (age, mouse ID)
$\mathcal{M}$: List. Stores the calculated mean latent vectors ($\overline{\mathbf{z}}_m$) for each mouse subject
$k(\mathbf{u}, \mathbf{v})$: Function. Multi-scale Gaussian Radial Basis Function (RBF) kernel
$\sigma_{base}$: Scalar. Base bandwidth parameter for the RBF kernel, established using the median heuristic
MMD: Scalar. Empirical Maximum Mean Discrepancy (biased V-statistic estimator).

### References


[1] Tabula Muris Consortium. A single-cell transcriptomic atlas characterizes ageing tissues in the mouse. *Nature*, 583(7817):590–595, July 2020.

[2] F Alexander Wolf, Philipp Angerer, and Fabian J Theis. SCANPY: large-scale single-cell gene expression data analysis. *Genome Biology*, 19(1):15, February 2018.

[3] Tim Stuart, Andrew Butler, Paul Hoffman, Christoph Hafemeister, Efthymia Papalexi, William M Mauck, III, Yuhan Hao, Marlon Stoeckius, Peter Smibert, and Rahul Satija. Comprehensive integration of Single-Cell data. *Cell*, 177(7):1888–1902.e21, June 2019.

[4] Mohammad Lotfollahi, F Alexander Wolf, and Fabian J Theis. scgen predicts single-cell perturbation responses. *Nature Methods*, 16(8):715–721, August 2019.

[5] Diederik P Kingma and Max Welling. Auto-encoding variational bayes, 2022. URL https://arxiv.org/abs/1312.6114.

[6] Adam Gayoso, Romain Lopez, Galen Xing, Pierre Boyeau, Valeh Valiollah Pour Amiri, Justin Hong, Katherine Wu, Michael Jayasuriya, Edouard Mehlman, Maxime Langevin, Yining Liu, Jules Samaran, Gabriel Misrachi, Achille Nazaret, Oscar Clivio, Chenling Xu, Tal Ashuach, Mariano Gabitto, Mohammad Lotfollahi, Valentine Svensson, Eduardo da Veiga Beltrame, Vitalii Kleshchevnikov, Carlos Talavera-López, Lior Pachter, Fabian J Theis, Aaron Streets, Michael I Jordan, Jeffrey Regier, and Nir Yosef. A python library for probabilistic analysis of single-cell omics data. *Nature Biotechnology*, 40(2):163–166, February 2022.

[7] Adam Paszke, Sam Gross, Francisco Massa, Adam Lerer, James Bradbury, Gregory Chanan, Trevor Killeen, Zeming Lin, Natalia Gimelshein, Luca Antiga, Alban Desmaison, Andreas Köpf, Edward Yang, Zach DeVito, Martin Raison, Alykhan Tejani, Sasank Chilamkurthy, Benoit Steiner, Lu Fang, Junjie Bai, and Soumith Chintala. Pytorch: An imperative style, high-performance deep learning library, 2019. URL https://arxiv.org/abs/1912.01703.

[8] Diederik P. Kingma and Jimmy Ba. Adam: A method for stochastic optimization, 2017. URL https://arxiv.org/abs/1412.6980.

[9] Irina Higgins, Loic Matthey, Arka Pal, Christopher Burgess, Xavier Glorot, Matthew Botvinick, Shakir Mohamed, and Alexander Lerchner. beta-VAE: Learning basic visual concepts with a constrained variational framework. In *International Conference on Learning Representations*, 2017. URL https://openreview.net/forum?id=Sy2fzU9gl.

[10] Jordan W Squair, Matthieu Gautier, Claudia Kathe, Mark A Anderson, Nicholas D James, Thomas H Hutson, Rémi Hudelle, Taha Qaiser, Kaya J E Matson, Quentin Barraud, Ariel J Levine, Gioele La Manno, Michael A Skinnider, and Grégoire Courtine. Confronting false discoveries in single-cell differential expression. *Nat. Commun.*, 12(1):5692, September 2021.

[11] Yusuf Roohani, Kexin Huang, and Jure Leskovec. Predicting transcriptional outcomes of novel multigene perturbations with GEARS. *Nature Biotechnology*, 42(6):927–935, June 2024.

[12] Peng Qiu. Embracing the dropouts in single-cell RNA-seq analysis. *Nat. Commun.*, 11(1):1169, March 2020.

[13] Gökcen Eraslan, Lukas M Simon, Maria Mircea, Nikola S Mueller, and Fabian J Theis. Single-cell RNA-seq denoising using a deep count autoencoder. *Nat. Commun.*, 10(1):390, January 2019.

[14] David van Dijk, Roshan Sharma, Juozas Nainys, Kristina Yim, Pooja Kathail, Ambrose J Carr, Cassandra Burdziak, Kevin R Moon, Christine L Chaffer, Diwakar Pattabiraman, Brian Bierie, Linas Mazutis, Guy Wolf, Smita Krishnaswamy, and Dana Pe'er. Recovering gene interactions from single-cell data using data diffusion. *Cell*, 174(3):716–729.e27, July 2018.

[15] Wenpin Hou, Zhicheng Ji, Hongkai Ji, and Stephanie C Hicks. A systematic evaluation of single-cell RNA-sequencing imputation methods. *Genome Biol.*, 21(1):218, August 2020.

[16] Arthur Gretton, Karsten M. Borgwardt, Malte J. Rasch, Bernhard Schölkopf, and Alexander Smola. A kernel two-sample test. *Journal of Machine Learning Research*, 13(25):723–773, 2012. URL http://jmlr.org/papers/v13/gretton12a.html.

[17] Mingsheng Long, Yue Cao, Jianmin Wang, and Michael Jordan. Learning transferable features with deep adaptation networks. In Francis Bach and David Blei, editors, *Proceedings of the 32nd International Conference on Machine Learning*, volume 37 of *Proceedings of Machine Learning Research*, pages 97–105, Lille, France, 07–09 Jul 2015. PMLR. URL https://proceedings.mlr.press/v37/long15.html.

[18] Zhuoqing Fang, Xinyuan Liu, and Gary Peltz. Gseapy: a comprehensive package for performing gene set enrichment analysis in python. *Bioinformatics*, 39(1):btac757, 01 2023. ISSN 1367-4811. doi: 10.1093/bioinformatics/btac757. URL https://doi.org/10.1093/bioinformatics/btac757.

[19] M Kanehisa and S Goto. KEGG: kyoto encyclopedia of genes and genomes. *Nucleic Acids Res.*, 28 (1):27–30, January 2000.

[20] Carol J Bult, Judith A Blake, Cynthia L Smith, James A Kadin, Joel E Richardson, and Mouse Genome Database Group. Mouse genome database (MGD) 2019. *Nucleic Acids Res.*, 47(D1): D801–D806, January 2019.

[21] Michael Ashburner, Catherine A Ball, Judith A Blake, David Botstein, Heather Butler, J Michael Cherry, Allan P Davis, Kara Dolinski, Selina S Dwight, Janan T Eppig, Midori A Harris, David P Hill, Laurie Issel-Tarver, Andrew Kasarskis, Suzanna Lewis, John C Matese, Joel E Richardson, Martin Ringwald, Gerald M Rubin, and Gavin Sherlock. Gene ontology: tool for the unification of biology. *Nature Genetics*, 25(1):25–29, May 2000.

[22] The Gene Ontology Consortium. The gene ontology knowledgebase in 2026. *Nucleic Acids Research*, 54(D1):D1779–D1792, 01 2026. ISSN 1362-4962. doi: 10.1093/nar/gkaf1292. URL https://doi.org/10.1093/nar/gkaf1292.

[23] Dénes Türei, Tamás Korcsmáros, and Julio Saez-Rodriguez. OmniPath: guidelines and gateway for literature-curated signaling pathway resources. *Nat. Methods*, 13(12):966–967, November 2016.

[24] Luz Garcia-Alonso, Christian H. Holland, Mahmoud M. Ibrahim, Denes Turei, and Julio Saez-Rodriguez. Benchmark and integration of resources for the estimation of human transcription factor activities. *Genome Research*, 29(8):1363–1375, 2019. doi: 10.1101/gr.240663.118. URL http://genome.cshlp.org/content/genome/29/8/1363.

[25] Lawrence Page, Sergey Brin, Rajeev Motwani, and Terry Winograd. The pagerank citation ranking : Bringing order to the web. In *The Web Conference*, 1999. URL https://api.semanticscholar.org/CorpusID:1508503.

[26] Aric A. Hagberg, Daniel A. Schult, and Pieter J. Swart. Exploring network structure, dynamics, and function using networkx. In Gaël Varoquaux, Travis Vaught, and Jarrod Millman, editors, *Proceedings of the 7th Python in Science Conference*, pages 11 – 15, Pasadena, CA USA, 2008.

[27] Ville Satopaa, Jeannie Albrecht, David Irwin, and Barath Raghavan. Finding a "kneedle" in a haystack: Detecting knee points in system behavior. In *Proceedings of the 2011 31st International Conference on Distributed Computing Systems Workshops*, ICDCSW '11, page 166–171, USA, 2011. IEEE Computer Society. ISBN 9780769543864. doi: 10.1109/ICDCSW.2011.20. URL https://doi.org/10.1109/ICDCSW.2011.20.

[28] The UniProt Consortium. Uniprot: the universal protein knowledgebase in 2025. *Nucleic Acids Research*, 53(D1):D609–D617, 01 2025. ISSN 1362-4962. doi: 10.1093/nar/gkae1010. URL https://doi.org/10.1093/nar/gkae1010.

[29] Paul Shannon, Andrew Markiel, Owen Ozier, Nitin S Baliga, Jonathan T Wang, Daniel Ramage, Nada Amin, Benno Schwikowski, and Trey Ideker. Cytoscape: a software environment for integrated models of biomolecular interaction networks. *Genome Res.*, 13(11):2498–2504, November 2003.

[30] Pauli Virtanen, Ralf Gommers, Travis E. Oliphant, Matt Haberland, Tyler Reddy, David Cournapeau, Evgeni Burovski, Pearu Peterson, Warren Weckesser, Jonathan Bright, Stéfan J. van der Walt, Matthew Brett, Joshua Wilson, K. Jarrod Millman, Nikolay Mayorov, Andrew R. J. Nelson, Eric Jones, Robert Kern, Eric Larson, C J Carey, İlhan Polat, Yu Feng, Eric W. Moore, Jake VanderPlas, Denis Laxalde, Josef Perktold, Robert Cimrman, Ian Henriksen, E. A. Quintero, Charles R. Harris, Anne M. Archibald, Antônio H. Ribeiro, Fabian Pedregosa, Paul van Mulbregt, and SciPy 1.0 Contributors. SciPy 1.0: Fundamental Algorithms for Scientific Computing in Python. *Nature Methods*, 17:261–272, 2020. doi: 10.1038/s41592-019-0686-2.

[31] Charles R. Harris, K. Jarrod Millman, Stéfan J. van der Walt, Ralf Gommers, Pauli Virtanen, David Cournapeau, Eric Wieser, Julian Taylor, Sebastian Berg, Nathaniel J. Smith, Robert Kern, Matti Picus, Stephan Hoyer, Marten H. van Kerkwijk, Matthew Brett, Allan Haldane, Jaime Fernández del Río, Mark Wiebe, Pearu Peterson, Pierre Gérard-Marchant, Kevin Sheppard, Tyler Reddy, Warren Weckesser, Hameer Abbasi, Christoph Gohlke, and Travis E. Oliphant. Array programming with NumPy. *Nature*, 585(7825):357–362, September 2020. doi: 10.1038/s41586-020-2649-2. URL https://doi.org/10.1038/s41586-020-2649-2.

[32] Florian Charlier, Marc Weber, Dariusz Izak, Emerson Harkin, Marcin Magnus, Joseph Lalli, Louison Fresnais, Matt Chan, Nikolay Markov, Oren Amsalem, Sebastian Proost, Agamemnon Krasoulis, getzze, and Stefan Repplinger. Statannotations, October 2022. URL https://doi.org/10.5281/zenodo.7213391.

[33] Yoav Benjamini and Yosef Hochberg. Controlling the false discovery rate: A practical and powerful approach to multiple testing. *Journal of the Royal Statistical Society: Series B (Methodological)*, 57(1):289–300, 01 1995. ISSN 0035-9246. doi: 10.1111/j.2517-6161.1995.tb02031.x. URL https://doi.org/10.1111/j.2517-6161.1995.tb02031.x.

[34] Franziska Paul, Ya'ara Arkin, Amir Giladi, Diego Adhemar Jaitin, Ephraim Kenigsberg, Hadas Keren-Shaul, Deborah Winter, David Lara-Astiaso, Meital Gury, Assaf Weiner, Eyal David, Nadav Cohen, Felicia Kathrine Bratt Lauridsen, Simon Haas, Andreas Schlitzer, Alexander Mildner, Florent Ginhoux, Steffen Jung, Andreas Trumpp, Bo Torben Porse, Amos Tanay, and Ido Amit. Transcriptional heterogeneity and lineage commitment in myeloid progenitors. *Cell*, 163(7):1663–1677, December 2015.

[35] Ian G Cowell and Caroline A Austin. Myeloperoxidase inhibition protects bone marrow mononuclear cells from DNA damage induced by the TOP2 poison anti-cancer drug etoposide. *FEBS Open Bio*, 14(6):1001–1010, June 2024.

[36] Gjin Ndrepepa. Myeloperoxidase - a bridge linking inflammation and oxidative stress with cardiovascular disease. *Clin. Chim. Acta*, 493:36–51, June 2019.

[37] Hartmut Geiger, Gerald de Haan, and M Carolina Florian. The ageing haematopoietic stem cell compartment. *Nat. Rev. Immunol.*, 13(5):376–389, May 2013.
